# Regulatory logic of neuronal differentiation in the *Drosophila* visual system

**DOI:** 10.1101/2025.09.01.673531

**Authors:** McKenzie Treese, Yen-Chung Chen, Abigail Tyree, Rose Coyne, Cathleen Lake, Ojong Besong Tabi, Raghuvanshi Rajesh, Yu-Chieh David Chen, Huzaifa Hassan, Hua Li, Claude Desplan, Mehmet Neset Özel

## Abstract

Combinations of terminal selector transcription factors (tsTFs) are thought to establish and maintain the unique identities of the numerous cell types found in nervous systems. However, it remains largely unclear how tsTF combinations are specified during development, and how they then coordinate the type-specific differentiation programs of each neuron. To investigate these regulatory mechanisms, we performed simultaneous single-cell RNA and ATAC sequencing on the *Drosophila* optic lobes at four stages of their development and identified over 250 distinct cell types. We characterized the common cis-regulatory features of neuronal enhancers and performed comprehensive inference of gene regulatory networks across cell types and stages. Our results reveal cell-type and stage-specific enhancers of many neuronal genes and the cooperative actions of tsTFs, pan-neuronal and ecdysone-responsive TFs on these enhancers. We show that the same effector genes are often regulated by different tsTF combinations acting through distinct enhancers in different neurons. During neurogenesis, tsTF codes are established within a brief critical period in newborn neurons, often through cell-type-specific enhancers that are not accessible in their progenitors. Accordingly, when neuroblast temporal patterning TFs are re-utilized as tsTFs in neurons, they are regulated independently through separate enhancers. Therefore, neuronal identity specification and differentiation is a multi-step regulatory program, wherein the same TFs enact distinct regulatory codes at different steps and across cell types.

## Introduction

Neurons are by far the most heterogeneous cell type in animals. Their vast diversity in connectivity and function directly underlies the intricacies of sensory processing and the precision of cognitive and motor functions. While the wide morphological diversity of neurons has been appreciated since the early days of Cajal (1), rapid popularization of single-cell genomic (in particular transcriptomic (2, 3)) technologies over the last decade has delivered a renewed interest and expanded understanding of neuronal diversity across species. The most recent single-cell RNA sequencing (scRNA-seq) studies of the mouse brain identified over 5,000 molecularly distinct neurons (4, 5). Different combinations of transcription factors (TFs) expressed in various neuronal types encode their distinct identities, instructing cell-type-specific morphological (6–9), synaptic (10–13) and electrophysiological (14) features. However, little is known about the gene regulatory mechanisms that link differential TF expression to the specific morphology, connectivity and activity of each neuron. It is also unclear how the genomes instruct the emergence of such a wide variety of cell types from a limited pool of neural stem cells.

*Drosophila* has long been a premier model system for neurobiology. The complete connectome of its brain at electron microscopy (EM) resolution has recently become available (15–17), revealing over 5,000 neuronal types. The visual system in particular offers distinct advantages to study neuronal diversity because its development is thought to be genetically hardwired (18, 19), *i.e.,* it does not require experience. Despite constituting two-thirds of the fly brain, the optic lobes contain only a few hundred cell types (16, 17) due to the repetitive nature of their neural circuits, a characteristic of all visual systems (20). This is in contrast to the rest of the brain where thousands of different cell types have been identified, many of which exist as only one (or a few) neuron(s) per hemibrain (15, 21). Each optic lobe receives direct input from photoreceptors of the retina and processes visual information in its four neuropils: lamina, medulla, lobula and lobula plate (22), which have a layered and columnar structure analogous to the vertebrate retina and cortex (20). We previously published a large scRNA-seq atlas of the *Drosophila* optic lobe, resolving ∼200 distinct cell-types across 6 developmental stages (23). This characterization revealed remarkable consistency between the molecular and classical (22) (defined morphologically by light microscopy) cell types. Although some of our transcriptomic clusters contained multiple morphologically distinct neurons (24–26), these were all due to limited coverage of rarer cell types.

Optic lobe neurons are generated during late larval and early pupal stages from two main neurogenic regions: the inner and the outer proliferation centers (IPC and OPC). Neuroepithelial (NE) progenitors in both regions produce neural stem cells (neuroblasts, NB), which undergo a series of asymmetric divisions generating intermediate progenitors known as ganglion mother cells (GMC). GMCs then divide just once to produce two different neurons (or glia). The IPC primarily produces the C/T neurons (except T1) that are positioned near the lobula plate, while the OPC is more complex. Lamina neurons are generated from the lateral OPC, while the medial OPC primarily produces medulla neurons. At least 3 distinct axes of patterning mechanisms contribute to generating neuronal diversity from the OPC: *(i)* spatial patterning of NE progenitors by a combination of signaling molecules and 5 (known) spatial TFs (27–29); *(ii)* temporal patterning with a series of >12 temporal TFs (tTFs) expressed sequentially in NBs (30); and *(iii)* Notch signaling between sister neurons after the asymmetric terminal division of GMCs (31).

Interestingly, expression of spatial and temporal TFs is not commonly maintained in postmitotic neurons (30). Instead, unique combinations of terminal selectors (tsTFs) that are stably expressed in each neuron throughout its lifetime have been proposed to implement and maintain neuronal type identity (32–35). We recently deciphered the codes of putative tsTFs in all optic lobe neurons using scRNA-seq, and were able to genetically modify some of these codes to induce complete and predictable changes in identity between different neuronal types (36). Therefore, tsTFs link the early cell fate determination events in neural progenitors to the acquisition of specific morphology, connectivity and physiology of each neuron. Using correlative models, we have additionally shown that the combination of spatial domain and temporal window of origin, together with the Notch status of each neuron, can fully account for the diversity of tsTF expression in OPC neurons (37). However, little is known about the gene regulatory networks upstream and downstream of tsTFs, which hold the key for understanding the molecular genetic mechanisms that underlie neuronal diversity.

To provide a comprehensive description of these regulatory networks in the *Drosophila* visual system, we generated a large single-cell multiomic (simultaneous RNA+ATAC-seq) dataset of the optic lobes collected at four different stages from neurogenesis to adulthood. With ∼250 clusters consistently resolved across these stages, the cell-type resolution of this new atlas exceeds that of our scRNA-seq dataset (23) and provides a robust framework for regulatory analyses. We comprehensively characterized the cis-regulatory features of neuronal enhancers (accessible in all or some neurons) with various developmental dynamics, as well as the key TF networks across all optic lobe cell types and stages. Our results indicate that most neuronal genes are regulated by different tsTF combinations, acting through distinct enhancers in different optic lobe cell types. Pan-neuronal, terminal selector and ecdysone-regulated TFs cooperate extensively on neuronal enhancers to ensure that each effector gene is expressed in the correct cell type(s) and developmental stage(s). Our analysis revealed a particularly dramatic shift in the chromatin landscape upon the terminal division of GMCs, indicating a dynamic gene regulatory network that operates in newborn neurons and integrates all the upstream (spatial, temporal, Notch) patterning inputs to establish the permanent tsTF combination of each neuron. Consistently, we found that the OPC temporal TFs that are also expressed post-mitotically as tsTFs are almost always associated with different enhancers in NBs vs. neurons, indicating their independent deployment in different regulatory programs.

## Results

### A multiomic single-cell sequencing atlas of the developing *Drosophila* visual system

We performed single-cell RNA and ATAC sequencing on nuclei isolated from *Drosophila* optic lobes at multiple stages of their development: P0 (neurogenesis), P24 (axon guidance), P48 (synaptogenesis) and Adult (functional circuit), using *Chromium Single Cell Multiome* kits (*10x Genomics*). After doublet removal and other quality controls, we retained a total of 232,251 cell barcodes with high-quality transcriptomes and epigenomes (chromatin accessibility) jointly available for further analyses. The UMAP reductions calculated separately on RNA and ATAC modalities are shown for each stage we sequenced (**Fig. S1**), but for all other figures we used the Weighted Nearest Neighbor (WNN) algorithm implemented in the *Seurat* and *Signac* packages (38, 39), which takes advantage of both modalities to calculate a joint UMAP reduction (**Fig. 1**). Key quality control metrics for these cells (number of UMIs, genes, fragments, as well as nucleosome signal and TSS enrichment values) are shown per library and strain (**Fig. S2-3**), as well as overlayed on the UMAPs of each stage (**Figs. S4-5**). Most notably, the ATAC modality of this new atlas provides a 5-10x improvement on the sequencing depth (unique fragments per cell) of the previously published scATAC-seq atlas of the fly brain (40).

**Figure 1:**
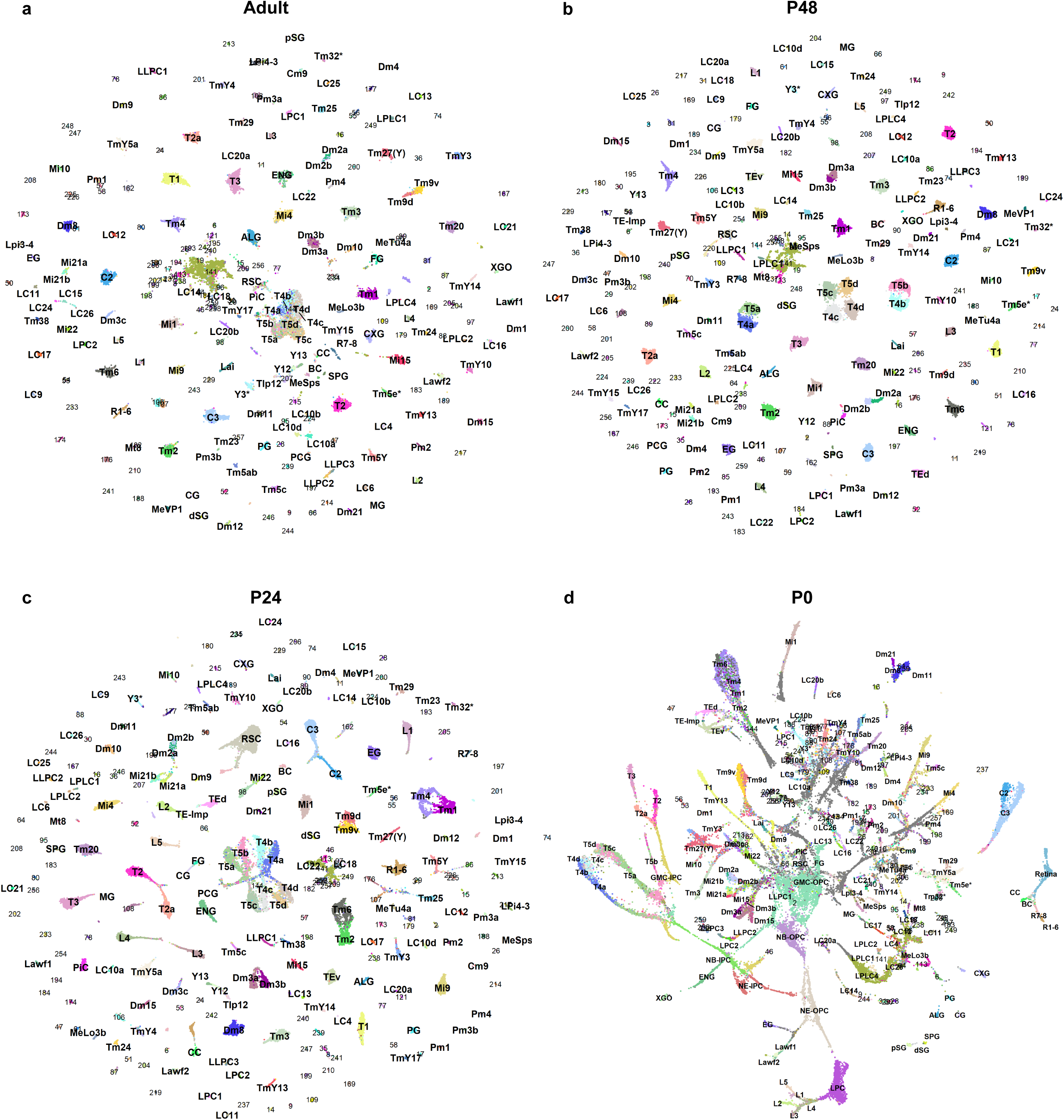
A multiomic single-cell sequencing atlas of the *Drosophila* visual system. **a-d**, UMAP reductions of single-cell multiomic profiles from the developing *Drosophila* optic lobe, shown separately for each developmental stage: (**a**) Adult, (**b**) P48, (**c**) P24, each calculated on 150 PCA and LSI components, and (**d**) P0, calculated on 70 PCA and LSI components. All embeddings were generated using weighted nearest neighbor (WNN) integration of RNA (SCT) and ATAC modalities.

We classified these cells into cell types using two methods. First, we took a supervised approach (**Fig. S5f**), using a neural network classifier we had trained on the marker gene expression profiles of 206 clusters that we previously resolved with scRNA-seq (23). In addition, we performed unsupervised clustering using the WNN algorithm on the 63,726 cells of the P48 stage, as we and others have previously shown that neurons display the highest molecular diversity at this stage (23, 41). Upon comparing the supervised and unsupervised classification results (**Fig. S6**), we found that WNN clustering divided 32 clusters of our previous scRNA-seq atlas into 2-6 subgroups each, amounting in fact to 85 clusters in this new dataset. In contrast, in only 3 cases the new clustering failed to resolve a previously known subdivision, which could be attributed to the very small sizes of these clusters. We integrated the results of both approaches into a new reference atlas containing 259 clusters (**Table S1**), which we then applied to other stages (**Fig. 1**) using a multimodal label transfer algorithm (SI Methods). As we described previously (23, 30) and discuss further in later sections, only the relatively differentiated cells (tips of the trajectories in **Fig. 1d**) could be classified into these clusters at the P0 stage, i.e. during active neurogenesis.

The improved cell-type resolution of this dataset (233 neuronal clusters, compared to 185 in the previous atlas) is largely attributable to higher depth in the ATAC modality (**Fig. S1**). However, our data do not support any molecular diversity among the postmitotic neurons that could only be detected in chromatin accessibility, since all newly identified subclusters had also differentially expressed genes among them. Thus, we surmise that these clusters would have been resolvable with transcriptomics given enough coverage (number of cells) and depth (reads/cell), which the published optic lobe scRNA-seq atlases (23, 42) did not reach.

Because of this very high granularity of our cell-type classification, most of our clusters do not contain enough cells to reliably determine their ATAC peaks when treated individually. Thus, we generated a hierarchical tree of all postmitotic clusters (those consistently assigned from P24 to Adult) based on their transcriptomic similarity at P24 and grouped them into 37 neuronal and 5 non-neuronal ‘metaclusters’, each containing 2 to 13 related clusters (**Fig. S7a-b**). The P0 dataset was independently divided into 19 metaclusters based on WNN clustering at this stage (**Fig. S7d**). We called the ATAC peaks separately on each of these 61 metaclusters using *MACS2* (43), and then combined these into 76,777 (500 bp) ‘consensus’ peaks (SI Methods) for optic lobe-wide analyses of differential accessibility.

### Cell type annotations

In our previous scRNA-seq atlas (23), we had annotated ∼70 clusters as specific neuronal types; glial clusters were also annotated in another study (44). Subsequent work has defined molecular markers for many additional optic lobe neurons, enabling their assignment to clusters (24–26, 45, 46), which we have incorporated into our new atlas (detailed in **Table S1**). Following the gene-specific *Gal4*/split-*Gal4* strategy described previously (46), we annotated 11 additional neuronal types: LC20b, MeLo3b, MeVP1, Cm9 (**Fig. S8**), Dm3c, Mi10, Mi22, MeTu4a, Y12, Y13 and Tlp12 (**Fig. S9**). We also used differential Bifid expression to separate the Mi21 cluster into Mi21a and Mi21b (**Fig. S8f**). Full details of each annotation, including the relevant split-*Gal4* intersections, are provided in the SI Methods and figure legends. Altogether, we were able to assign a cell-type identity to 17 of the 26 non-neuronal clusters (**Fig. S7b-c**), and to 122 of the 233 neuronal clusters (**Table S1**). Importantly, all morphologically distinct types of neurons that we have so far annotated in our datasets mapped to unique clusters without exception, further highlighting the very high resolution of this new atlas. In a few cases, molecularly distinct subtypes have been identified within morphologically homogenous neurons. We had previously described this phenomenon for Tm4, Tm6, Tm9 and TE neurons that differ when present dorsally or ventrally (23); and here we found a similar subdivision within the Dm2 cluster (**Fig. S8a**). We validated by HCR *in situ* hybridization experiments that these subclusters (Dm2a/b) represent mutually exclusive, but morphologically indistinguishable, subgroups of Dm2 neurons (**Fig. S8h**).

Finally, we trained a new multi-task neural network classifier on the marker gene expression profiles of our clusters at P24, P48 and Adult stages (SI Methods). This machine learning model can be used to automatically assign cell-type identities to any scRNA-seq dataset derived from the optic lobes. We applied it to reclassify the cells in two previously published scRNA-seq atlases of the developing optic lobes (23, 42), and both datasets (with newly assigned identities based on this new reference) can be explored using the web app scMarco (46) (https://apps.ycdavidchen.com/scMarco/).

### Dynamic gene regulatory landscapes of differentiating neurons

We previously identified tsTFs that maintain neuronal identity through their continuous expression from neuronal birth to adulthood (36). However, how these stable factors generate both cell-type-specific and dynamic gene expression programs throughout development and adulthood (23) remained unresolved. To understand these regulatory dynamics during neuronal maturation, we examined the differentially expressed genes (DEGs) and differentially accessible regions (DARs) among 229 neuronal clusters throughout development (excluding the 3 clusters of TE neurons that are not present in adults and cluster 141, which is a highly heterogeneous mixture of central brain contaminants). Across P24, P48 and Adult stages, we identified a total of 2,256 DEGs that were markers of one or more neuronal types in at least one stage (highly expressed in a cluster compared to average expression in all other neurons at the same stage, with a minimum fold-change of 2, **Table S2**). 976 of these were DEGs at all three stages, and 674 (30% of total) were consistent markers of at least one neuron throughout these stages (**Fig. 2a**). However, on a per-cluster basis, on average only ∼15% of marker genes were constant throughout development (**Fig. 2c**), implying that many genes that are continuously expressed in some cell types are dynamically regulated in others (i.e. Constant & Dynamic, **Fig. 2a**). Similarly, we found a total of 53,637 DARs among neurons across these stages (**Table S3**), with 29,190 found at all stages and 21,819 (40% of total) being a consistent marker for at least one neuron (**Fig. 2b**). But again, only ∼10% of marker peaks for any given cluster were consistent across all 3 stages (**Fig. 2c**). In contrast to most consensus peaks that were DARs among neurons, we found only 1,986 peaks that display relatively uniform accessibility among neurons with low or no accessibility in non-neuronal cells (pan-neuronal) in at least one of the three stages; 1,358 peaks maintained this pattern from P24 to Adult (consistent pan-neuronal). In summary, the majority of accessible regions we identified in the optic lobe are in fact differentially accessible both among neuronal types, and across developmental stages.

**Figure 2:**
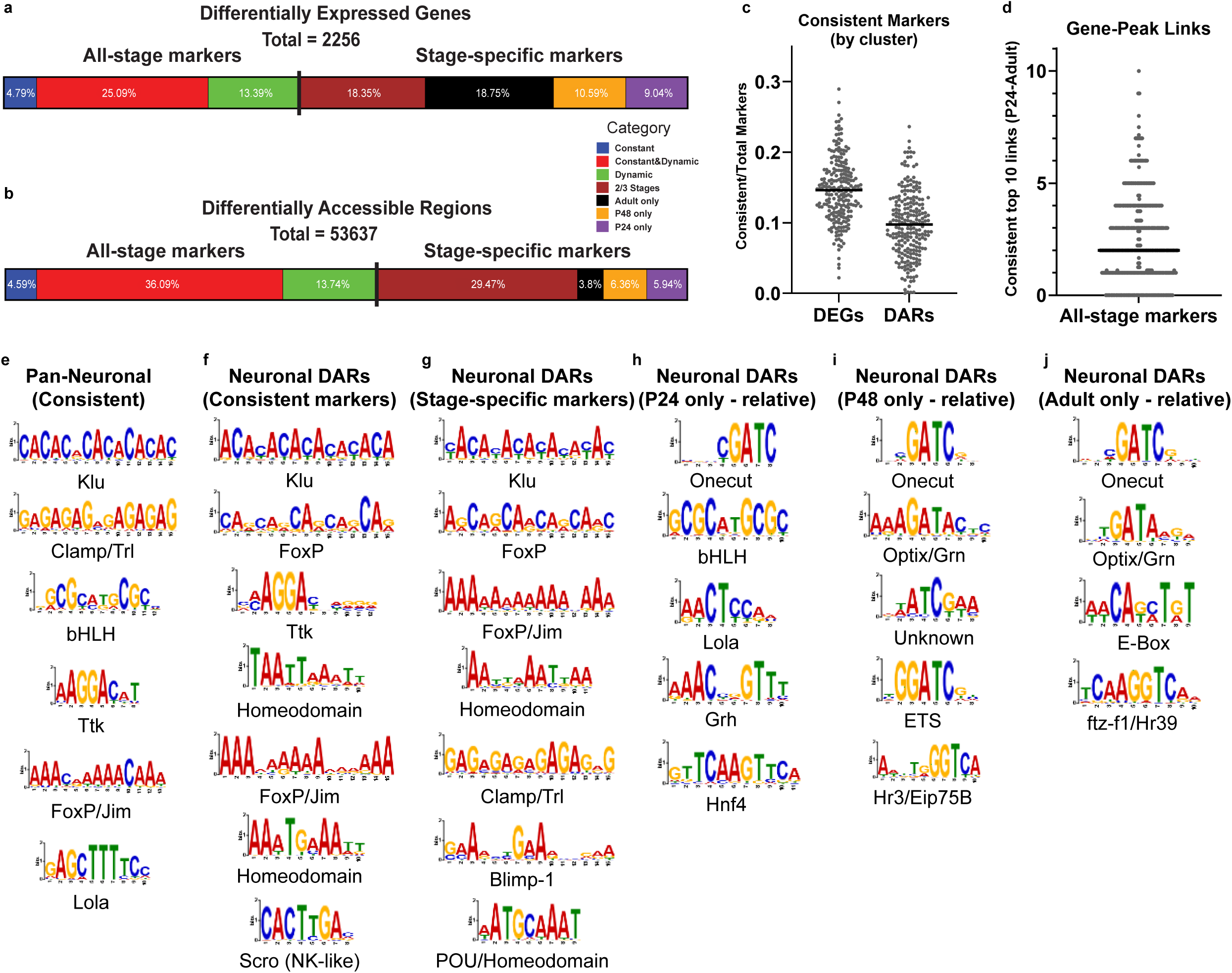
Dynamic gene regulatory landscapes of differentiating neurons. **a**, Distribution of the 2,256 DEGs captured across development. Left, “All-stage markers”: 976 markers found at all stages. These markers are further grouped as Dynamic (Green; dynamically regulated in all cell types), Constant and Dynamic (Red; continuously expressed in some cell types, but dynamically regulated in others), and Constant (Blue; continuously expressed in all cell types that express them). Right, “Stage-specific markers”: The remaining 1280 DEGs, which were captured as markers in 1 or 2 stages, but not all 3. Colors reflect the stages that these DEGs were found as neuronal markers. **b**, Same as in (a), but for DARs. 29,190 all-stage DARs are on the left, and 24,447 stage-specific DARs are on the right. **c**, Scatter plot depicting the proportion of markers (DEGs, left and DARs, right) shared across all 3 stages on a per-cluster basis. **d**, Strip plot depicting the counts of top-10 gene-peak links that are consistently found across development (P24-Adult), corrected for genes with <10 peaks linked. **e-j**, STREME results depicted as sequence logos, ordered (top to bottom) based on E-value of enrichment (see Table S5). The names were assigned based on Tomtom results matching these PWMs to the known *Drosophila* TF motifs. **e**, Motifs enriched in consistent pan-neuronal peaks (not DARs). **f**, Motifs enriched in consistent neuronal DARs. **g**, Motifs enriched in DARs found at only one stage (P24, P48, or Adult). **h-j**, Motifs relatively enriched in DARs found at P24 (**h**), P48 (**i**) or Adult (**j**). Shuffled input sequences (**e-g**) or the sequences of consistent neuronal DARs (**h-j**) were used background for motif enrichment.

To further explore developmental enhancer usage, we calculated gene-enhancer links (47) by correlating gene expression and nearby peak accessibility (50kb window, SI Methods) across all clusters at each stage (**Table S4**). Of the 976 DEGs that were markers of at least one neuron at all three stages, 969 had at least one linked peak at each of the 3 stages. Therefore, we compared the peaks linked to these genes across development. On average (median) for each gene, only 2 out of the top 10 linked peaks (ranked by absolute z-score and corrected for genes with fewer than 10 total linked peaks) were common across stages (**Fig. 2d**), suggesting highly dynamic enhancer usage. Half (14/28) of the genes with stable enhancer usage (at least 7 of the top 10 linked peaks common across stages) were tsTFs, which constitute <10% of the consistent DEGs, highlighting their unique role in maintenance of cell type identity. This nevertheless represents a minority of all tsTFs, implying that they are also, and often, subject to differential enhancer usage during development.

To identify common cis-regulatory features of neuronal enhancers, we performed *de novo* motif discovery (using STREME (48)) on consistent pan-neuronal peaks (**Fig. 2e**), consistent neuronal DARs (**Fig. 2f**), and stage-restricted DARs (**Fig. 2g**), comparing the top motifs to known *Drosophila* TF motifs using Tomtom (49) (**Table S5**). Three motif classes were broadly enriched across these regions: CA-repeats, consistent with the binding preference of the progenitor-restricted repressor Klumpfuss (50); GA-repeats, consistent with the pioneer GAGA factors Clamp and Trl, both transiently expressed and required in progenitors (51, 52); and AGGA motifs, matching Tramtrack, which is expressed exclusively in non-neuronal clusters (**Figs. S10b-c**, **S11b,d**). Together, these features suggest that many neuronal enhancers are pioneered by GAGA factors and kept inactive by Klu in progenitors, and by Ttk in non-neuronal cells.

In addition to these putative pioneering and repressive TF binding sites, both pan-neuronal peaks and neuronal type-specific DARs were enriched in motifs likely corresponding to pan-neuronal TFs FoxP (**Figs. S10b, S11b,d**), Jim and Lola. Combined with the findings above, this supports a layered model of regulation for these enhancers, whereby they are first rendered “neuronal” by these common factors, with additional type-specificity provided by tsTFs. Indeed, only the neuronal DARs (both consistent and stage-specific), but not pan-neuronal peaks, were strongly enriched for homeodomain binding motifs. These motifs belong to the TF family that is most highly represented among terminal selectors (36, 53), and include variants such as NK-like and POU homeodomains (**Fig. 2f-g**). We found that stage-specific, but not consistent, neuronal DARs were additionally enriched in Blimp-1 motifs. *Blimp-1* encodes an ecdysone-responsive TF, highly expressed in all neurons during early pupal stages, which is turned off after P36 before being upregulated again in adult neurons (54).

To identify additional regulators of temporally dynamic chromatin accessibility, we investigated the motifs *relatively* enriched (using as background the sequences of consistent DARs) in peaks that were found as cluster markers only at P24 (3,187 peaks), P48 (3,413 peaks) or Adult (2,039 peaks) stages (**Fig. 2h-j**). Interestingly, ‘GATC’ motif of the pan-neuronal Onecut (**Figs. S10b, S11b,d**) was the top hit in all 3 stages. The contribution of the Cut family of TFs to regulation of neuron type-specific genes (in addition to pan-neuronal genes) has been established in *C. elegans* (*55*) and mouse spinal cord (56). Our findings further suggest that despite its constant expression in neurons, Onecut specializes in dynamically regulated enhancers (i.e. those that are accessible in a subset of neurons during only one stage) rather than consistent neuronal DARs. Finally, we identified a relative enrichment of different nuclear hormone-receptor (Hr) class of motifs that act downstream of ecdysone in stage-specific peaks, which further highlights the crucial role of this signaling pathway in temporally dynamic gene expression (57). While the core ‘GGTCA’ that is enriched at P48 is most consistent with the known motif of Hr3 that is upregulated during mid-pupal stages, the variant observed in the Adult-specific peaks is more similar to the known Ftz-f1 motif, another effector of ecdysone expressed at late pupal stages. These findings are generalizable to the entire optic lobe and reinforce the previous conclusion made in lamina neurons (54) that, despite their pan-neuronal expression, downstream targets of ecdysone-regulated TFs are predominantly cell-type specific.

Lastly, we used the ChromVAR algorithm (58) to calculate ‘motif activity scores’ (based only on chromatin accessibility) for the 72 optic lobe tsTFs with available consensus motifs in the CisBP database. We then calculated the correlations between these activity scores and the (RNA) expression levels of the respective TFs across all neurons at P24, P48 and Adult stages (**Fig. S12**). Many of them correlate best (negatively or positively) with their own motifs (the diagonal); however, ambiguities or mismatches that result from TF families with very similar motifs (e.g. homeodomains) or from strong co-expression of certain tsTFs among optic lobe neurons (e.g. *Dll* and *Ets65A*) are also observable. Overall, our global analyses of DARs reveal the extensive cooperation between various classes of pan-neuronal, terminal selector and ecdysone-responsive TFs for shaping spatiotemporally specific chromatin accessibility in developing neurons.

### Computational inference of developmental gene regulatory networks

In order to comprehensively understand neuron type-specific regulation, we performed computational gene regulatory network (GRN) inference using the SCENIC+ framework (59). To increase our statistical power, in addition to the multiome data from P24, P48 and Adult stages, we also utilized the scRNA-seq datasets of optic lobes previously published by us (23) and others (42) from similar development stages. These ‘RNA-only’ cells were included alongside the multiome cells for the steps that do not utilize cis-regulatory information, *i.e.* calculation of TF-target gene expression correlations with GRNBoost2, while all other inference steps were run with the multiome data only. Because the target genes regulated by a TF can be highly context-dependent, we applied GRN inference at the metacluster level (**Fig. S7a**), rather than to the whole optic lobe. However, since some of our metaclusters contain as few as 2 cell types and thus have limited biological variation for GRN inference, we merged some metaclusters containing related neurons (detailed in SI Methods). We inferred 28 different SCENIC+ networks (23 neuronal) covering all optic lobe cell types, which we provide in **Table S6**.

We first used these SCENIC+ results to characterize the context dependency of TF-target specificity in neuronal gene regulation (**Fig. 3a**). 33 TFs had regulons in at least 10 of the 23 neuronal GRNs; for all target genes of these TFs, across all networks, we determined the number of GRNs in which a TF was predicted to be regulating the same target gene (**Fig. 3b**). We observed that the majority of predicted targets for all 33 TFs were found in only one or two different GRNs, indicating that the same TFs in different metaclusters regulate highly divergent sets of target genes. In fact, out of 23,846 unique TF-target interactions predicted for these 33 TFs across all 23 GRNs, only 159 were found in more than 50% of the networks which include that TF as a regulator (to the right of the dashed line in **Fig. 3b**). Visualization of these 159 interactions as a network (**Fig. S13**) highlights that 69% of them are mediated by the (pan-neuronal) ecdysone-responsive TFs Eip75B, Hr3, Hr4 and Ftz-f1. These results indicate that while the ecdysone-regulated TFs occasionally regulate the same genes across a variety of different neurons in the optic lobe, downstream targets of tsTFs (and other DE TFs) are almost always context-specific, *i.e.,* limited to one (or a few) metacluster(s).

**Figure 3:**
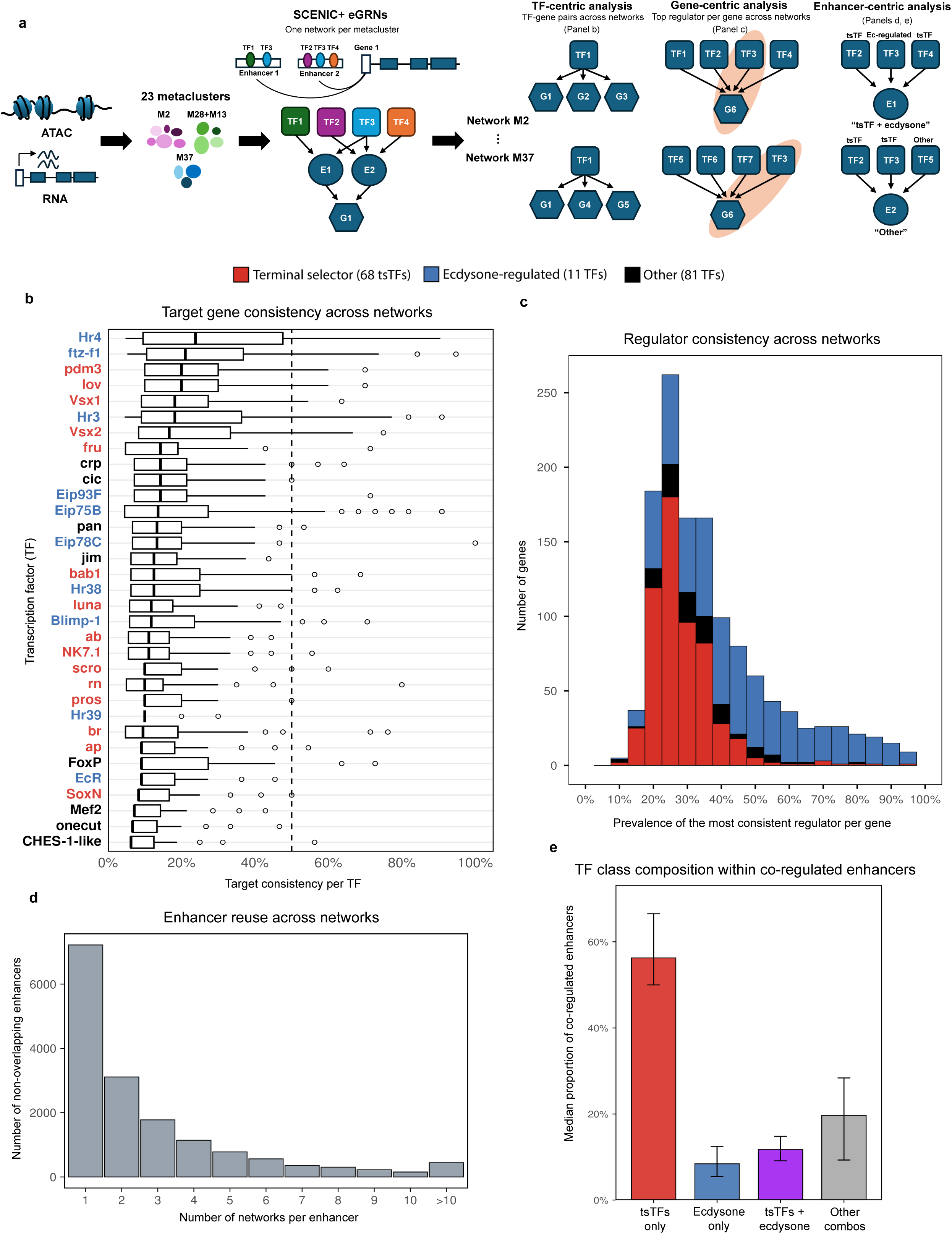
Context specificity of TF-target interactions. **a**, Schematic depicting SCENIC+ analysis strategy. The multiome data from each metacluster (after the merges as specified in SI Methods) were used to infer context-specific enhancer gene regulatory networks (eGRNs) using SCENIC+. Networks were generated separately for each of the 23 neuronal metaclusters, capturing TF-target relationships mediated through accessible enhancer regions. Neuronal networks were then analyzed from three complementary perspectives: “TF-centric analysis” quantifies how consistently TFs regulate the same genes across networks, considering only the TFs present in at least 10 networks. “Gene-centric analysis” identifies the most consistent regulator for each gene, considering only the (target) genes present in at least 10 networks. Finally, “Enhancer-centric analysis” examines both the enhancer reuse across networks, as well as the combinations of TFs predicted to co-operate on the same enhancers. **b**, Boxplots portraying the prevalence (%) of TF-gene pairs predicted across SCENIC+ networks, ordered by the median prevalence. The TFs shown on the y-axis were identified as regulators in at least 10 distinct neuronal GRNs and are colored by functional class: Terminal selector (red), Ecdysone-regulated (blue), or Other (black). **c**, Histogram depicting the persistence of top regulators (*i.e.,* the TF most consistently predicted to regulate each gene) across genes present in at least 10 different GRNs. Each stacked bar represents proportions of regulator classes, as colored in **b**. **d**, Histogram portraying the number of unique non-overlapping enhancers as a function of how many neuronal GRNs they appear in. Because each GRN was inferred using a different peak set, overlapping enhancer regions were first merged across networks, and the resulting merged regions were then mapped back to their corresponding regulons. **e**, Bar plot summarizing the median proportion of enhancer-target gene pairs (per network) that are regulated by terminal selector TFs only (tsTFs only, red), ecdysone-regulated TFs only (Ecdysone only, blue), a combination of terminal selector TFs with ecdysone-regulated TFs (tsTFs + ecdysone, purple), or any other combination involving other TFs (gray). Error bars indicate the interquartile range (IQR; 25^th^-75^th^ percentiles) around the median.

We next asked this question from the perspective of target genes. For 1,287 genes that were found in at least one regulon in at least 10 different GRNs, we determined which TFs were most consistently predicted to regulate each of these genes, and how often. On average, even the most consistent regulators were linked to a specific gene only in a minority (<50%) of GRNs that include this gene (**Fig. 3c**). These findings suggest that the same DEGs are most often regulated by different TFs in different cell types. Similarly, it reveals that the vast majority of highly consistent regulators for any given gene (found in >50% of GRNs including that target gene) were ecdysone-regulated TFs (**Fig. 3c**). We emphasize, however, that such consistently found interactions were rare, and thus cell type-specific regulation appears to be a general feature of all classes of neuronal TFs. These results suggest that most neuron type-specific genes are under highly modular regulation, whereby different TF combinations acting through distinct enhancers control their expression in a variety of different neurons.

To directly address the modularity of neuronal gene regulation, we compared the predicted enhancers across our SCENIC+ networks. Among the 23 neuronal GRN models, 16,058 non-overlapping regions were linked to the expression of a gene. Strikingly, 7,226 (45%) of these putative enhancers were identified (predicted to regulate a gene) in only one GRN, and only 441 (<3%) were found in more than 10 networks (**Fig. 3d**). We note that these different GRNs were inferred using our metacluster-specific peaks (rather than consensus peaks), and thus we counted any overlapping regions across different GRNs as potentially the same enhancer. However, in cases of partial overlaps, these may still represent different cis-regulatory regions, and thus neuronal enhancer usage is likely even more context-dependent than it appears in **Figure 3d**. In addition, we showed above that neuronal enhancers are strongly enriched in motifs for both tsTFs and ecdysone-responsive TFs (**Fig. 2g-j**), suggesting that effector genes are often regulated jointly by these different classes of TFs. To understand if they co-operate directly on specific enhancers, we investigated cases where at least two TFs were predicted to regulate the same target gene through the same enhancer in each of our 23 neuronal GRNs. Indeed, on average across these networks, ∼12% of all such enhancers were predicted to be bound by both tsTFs and ecdysone TFs (**Fig. 3e**), confirming that they indeed work together to control neural development.

We also used our SCENIC+ networks to identify the dominant regulators of different gene expression programs in developing optic lobes. For the 103 TFs with at least 50 unique targets, we ranked their fold-enrichment of target gene sets for the GO terms previously found among optic lobe cluster and stage markers (23). Regulators of cluster-marker categories (neurotransmitter receptors, synaptic target recognition, axon guidance, and ion channels; **Fig. S14a-d**) were uniformly dominated by tsTFs, providing strong support for the idea that most neuronal effector genes are directly regulated by terminal selectors. Ecdysone-responsive TFs (notably EcR and Hr3) also ranked among the top regulators of axon-guidance and ion-channel genes (respectively), reinforcing the joint tsTF/hormonal regulation described above. In contrast, stage-marker categories (translation in early pupal stages, oxidative phosphorylation in adults) were dominated by ubiquitous or pan-neuronal regulators (BtbVII, Crol, FoxP, CG4404, CrebA, Hr38; **Fig. S14e-f**), some of which have previously been implicated in activity-dependent transcription, potentially linking the onset of electrical activity in the optic lobe to the upregulation of energy-metabolism genes in maturing neurons.

Lastly, we plotted the accessibility and expression profiles of multiple loci across select cell types and stages (**Fig. S15**) to highlight specific examples of the different modes of neuronal gene regulation inferred by our genome-wide analyses above. The gene that encodes for the pan-neuronal active zone protein Bruchpilot (*brp*), which is upregulated in the optic lobe neurons during mid-pupal stages, is accordingly predicted to be under joint regulation of pan-neuronal TFs and the ecdysone TF Hr3 in multiple networks, supporting shared regulation across different contexts. Whereas the tsTF genes *fd59A* and *knot*, as well as the dopamine receptor *Dop2R* and the classical cholinergic genes *ChAT*/*VAChT* (transcribed from a shared locus) are regulated modularly through different enhancers (and upstream TFs) in different neurons, as typical of most neuronal genes. Some enhancers of *Dop2R* and *ChAT* are also predicted to be regulated by ecdysone TFs (by Blimp-1 and Hr38/Ftz-f1, respectively), which likely underlies the stage-specific upregulation of these genes in the indicated neurons.

It is important to emphasize that the regulatory interactions we inferred here using SCENIC+ are only predictions supported by our multiome data, with varying degrees of confidence; they should not be taken as ground truth in future studies. It should also be noted that the way we grouped our cell-types into metaclusters (*i.e.* based on transcriptomic similarity) to infer these GRNs introduces certain biases that need to be considered while interpreting these results. For example, the TFs or targets whose expression do not significantly vary either among the cell types (within a metacluster) or across the stages considered are unlikely to be captured. This could explain why the vast majority of the most consistently predicted interactions were downstream of the dynamically expressed ecdysone TFs, rather than the continuously expressed pan-neuronal TFs like Onecut or FoxP.

### Patterning regulators play mutually exclusive roles before and after cell cycle exit

While the multiome data from P24 to Adult stages enable us to explore the GRNs of postmitotic neurons during their terminal differentiation and circuit formation, the P0 dataset captures the events of neurogenesis that is ongoing until P15 (23). Since the same cell types are produced continuously in the optic lobe from late larval to early pupal stages (31, 60), a single snapshot at P0 can reconstruct the entire trajectory of neuronal identity specification (30), including neuroepithelia (NE), neuroblasts (NB) and ganglion mother cells (GMC), as well as young neurons (**Fig. 1d**). In addition, our dataset captures neurogenesis in both the retina and lamina. Thus, we separated these lineages into different UMAPs for additional analysis: retina (**Fig. S16a**), lamina (or lateral OPC, **Fig. S16b**), IPC (**Fig. S16c**), and medulla (medial OPC, **Fig. 4a-b**). Molecular mechanisms of cell-type specification are relatively well understood in the retina and lamina, and we will address IPC neurogenesis in a separate manuscript. Thereby we are focusing in this section on the medial OPC lineages that produce mostly medulla neurons.

**Figure 4:**
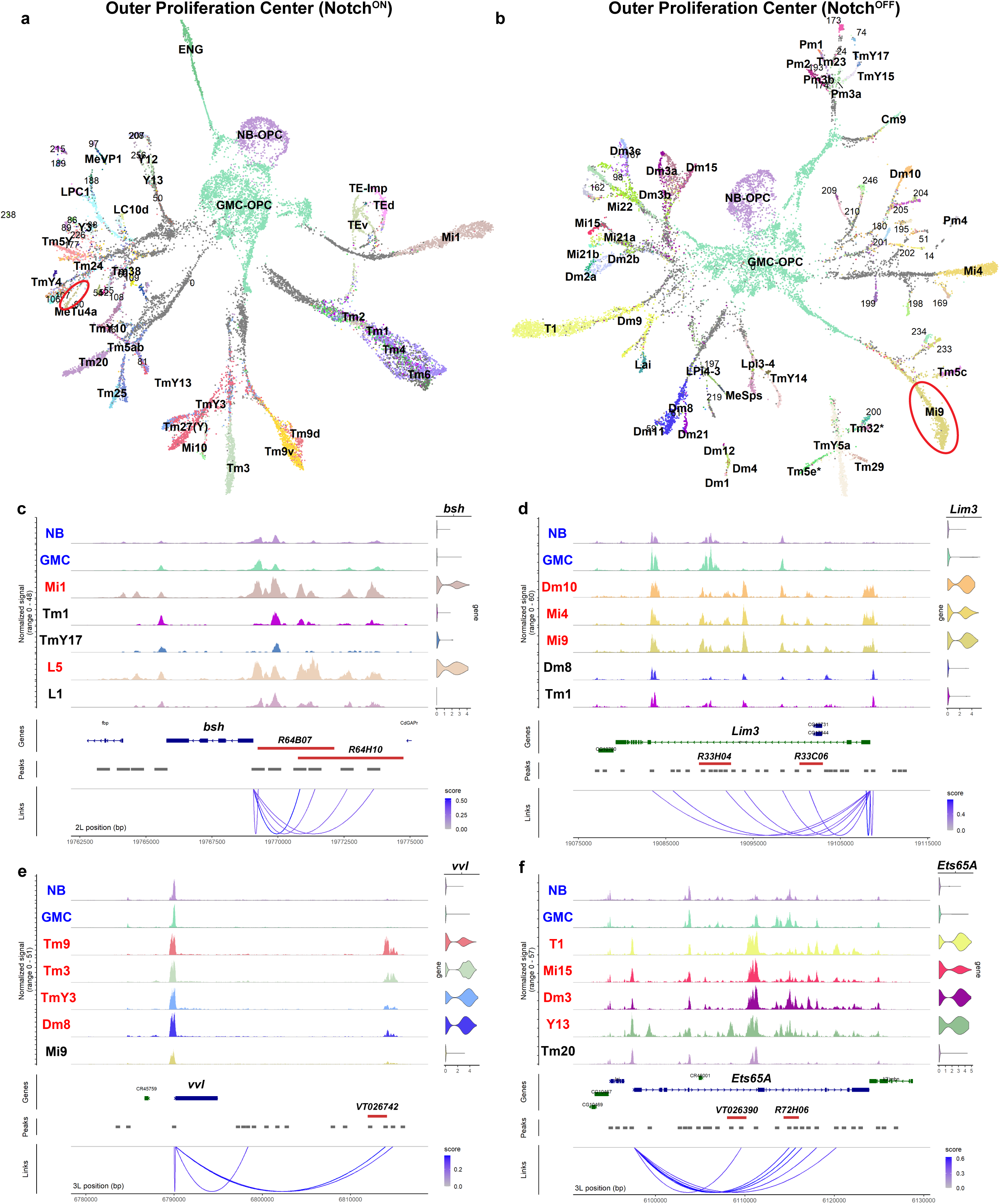
Differentiation trajectories during optic lobe neurogenesis. **a-b**, UMAP reductions of single-cell multiomic profiles of the medial OPC lineages in P0 optic lobes (from Fig. 1d). (**a**) Notch^ON^ hemilineage, calculated on 33 PCA and 29 LSI components; and (**b**) Notch^OFF^ lineage, calculated on 31 PCA and 38 LSI components. Neuronal identities were assigned based on label transfer from the P24 dataset (Fig. 1c). Young neurons for which these could not be confidently determined are shown in grey. The red ellipse in **a** marks MeTu4a neurons without *ap* expression despite their N^ON^ status (also in Fig. S10d). The red ellipse in **b** (also in Fig. S18) indicates Mi9 neurons. **c-f**, Coverage plots showing the accessibility of the *bsh* (c), *Lim3* (d), *vvl* (e) and *Ets65A* (f) loci at P0. For each gene, the progenitor clusters (NB and GMC) are shown together with representative neurons. Blue indicates the progenitor (NB and GMC) clusters; red, the neurons that express the gene as a tsTF; and black, the neurons that do not. The corresponding mRNA expression is shown as violin plots on the right. Red bars mark FlyLight and Vienna Tile enhancer fragments that overlap the indicated peaks (*bsh*: R64B07, R64H10; *Lim3*: R33H04, R33C06; *vvl*: VT026742; *Ets65A*: VT026390, R72H06; see Fig. S17 for reporter expression micrographs), and arcs (“Links”) connect each gene to its correlated peaks (putative enhancers), colored by link *score*. The links were filtered for: *z-score*>5 for *bsh*; *z-score*>5 and *score*>0.5 for *Lim3* and *Ets65A*; *score*>0.25 for *vvl*. All consensus peaks with links passing these filters fall within the depicted genomic ranges.

Of the three patterning mechanisms we introduced earlier, Notch status has the most significant effect on terminal identity of OPC neurons, such that sister (progeny of the same GMC) Notch^ON^/Notch^OFF^ (N^ON^/N^OFF^) neurons are more distinct from each other than other neurons with the same Notch status but different spatial or temporal origin (30). Thus, we separated the OPC progenies at P0 into N^ON^ (**Fig. 4a**) and N^OFF^ (**Fig. 4b**) hemilineages based on the immediate effectors of Notch signaling (*Hey* and *E(spl)* genes), which are only transiently expressed in postmitotic N^ON^ cells (**Fig. S10d-e**), and analyzed them separately. Expression of the tsTF Apterous (ap) is regulated by Notch (31) and has been treated as a universal marker of N^ON^ OPC neurons (37) (**Fig. S10d**). It is also the main regulator of their cholinergic identity (12), although this is not generalizable to the IPC or other brain regions. We identified one exception to this pattern in the OPC: MeTu4a neurons are cholinergic N^ON^ neurons (likely from a Slp/D temporal window), but do not express *ap* (**Figs. 4a** and **S10d**, red ellipse).

As representative examples, we plotted the chromatin accessibility and RNA expression of four tsTF genes (*bsh*, *vvl*, *Lim3* and *Ets65A*) at P0, in both the OPC progenitor clusters (NB and GMC) and in selected neuronal types that either do or do not express each gene (**Fig. 4c–f**). All four are likely regulated by multiple enhancers whose accessibility strongly correlates with gene expression (blue ‘links’). Several of these enhancers overlap fragments used to generate *Gal4* reporters in the FlyLight (61) and Vienna Tile (62) collections, confirming that they are active in optic lobe neurons at both late L3 (close to P0) and adult stages (**Fig. S17**). For *bsh*, two such fragments overlap a previously described set of enhancers (63) that drive expression in the lamina (L4/L5) and medulla (Mi1): both were active in lamina and medulla in adults, but during neurogenesis *R64B07* was specific to the lamina (**Fig. S17a**), whereas *R64H10* was enriched in the medulla (**Fig. S17b**). These tsTF enhancers were generally closed in progenitors. A few regions associated with *Lim3* (**Fig. 4d**) and *Ets65A* (**Fig. 4f**) were already accessible in NBs and GMCs; however, neither gene was detectably expressed, and both loci lacked promoter accessibility in the NB and GMC clusters. Consistent with this, the reporter fragments *R33H04* (in *Lim3*; **Fig. S17c**) and *R72H06* (in *Ets65A*; **Fig. S17g**), which overlap some of these pre-accessible regions, were inactive in OPC progenitors and active only in neurons. And even at these loci, most candidate enhancers become accessible only post-mitotically.

Different branches we observe on the UMAP trajectories of medial OPC hemilineages (**Fig. 4a-b**) roughly reflect the birth order of the depicted cell types (in clockwise direction). This allows their assignment to different NB temporal windows based on tTF expression patterns at the bases of these branches (**Fig. S18**), with a higher resolution than we had previously achieved with scRNA-seq (30). Interestingly, at least 6 tTFs (*hth*, *erm*, *ey*, *hbn*, *scro* and *D*) are also expressed as tsTFs in postmitotic neurons (30, 36). However, our trajectories show that none of these TFs are maintained in all neurons that originate from the corresponding temporal windows in either hemilineage (**Fig. S18**, arrows). And all of them are also independently activated in neurons originating outside of the corresponding temporal windows (**Fig. S18**, arrowheads), indicating little or no continuity in their regulation (summarized in **Fig. 5a**). Indeed, there is a strong (but still not perfect) correlation between the temporal window of origin and the neuronal expression of these genes for only two tTFs: *hth* (in both hemilineages) and *scro* (only in the N^OFF^ hemilineage). We thus suspected that even in cases of apparent maintenance (*i.e.,* a tTF remains expressed as a tsTF in a neuron that originates from its own temporal window), expression of these genes before and after cell cycle exit is likely to be controlled independently.

**Figure 5:**
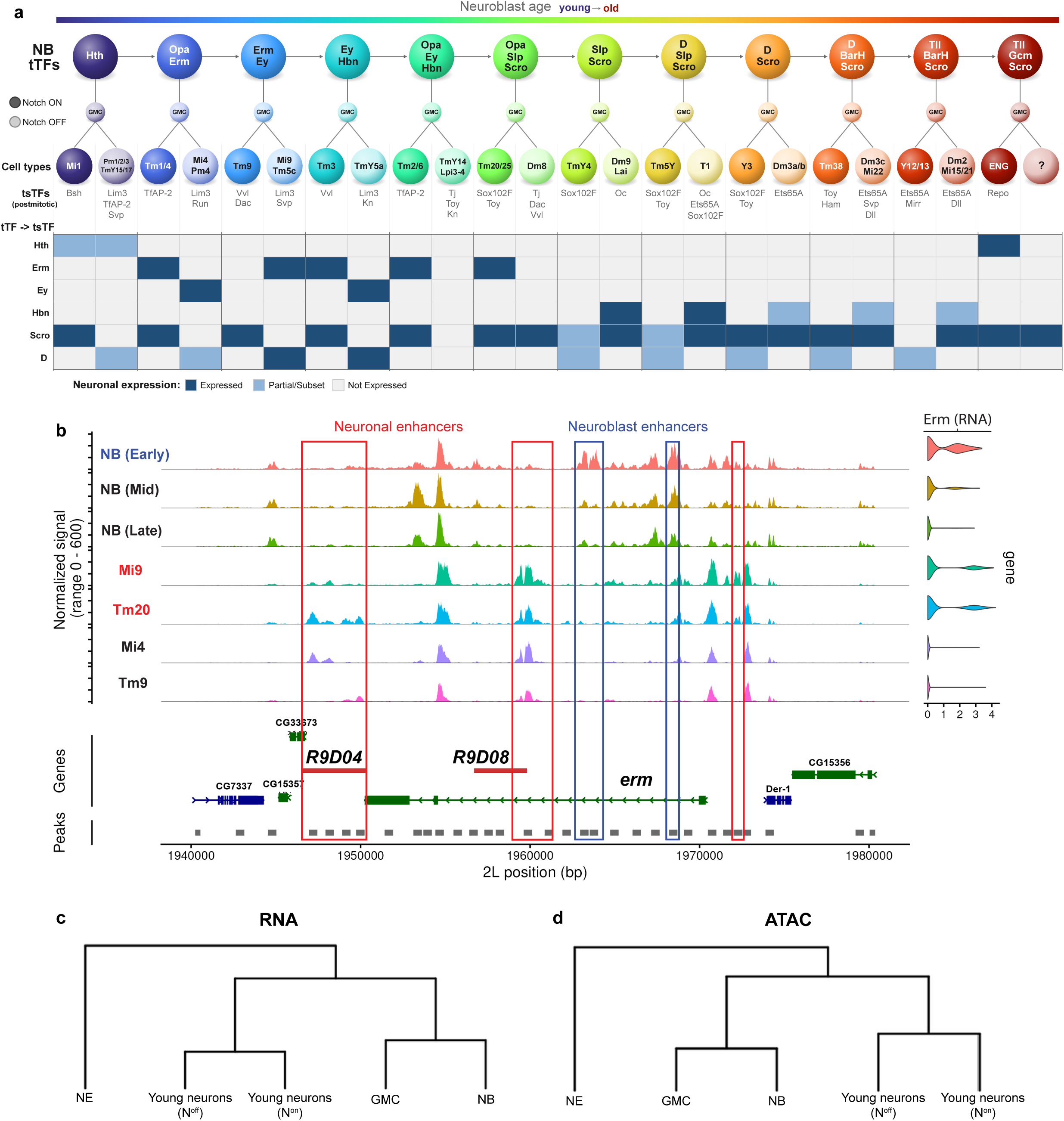
Independent regulation of temporal TFs in neuroblasts and neurons. **a**, Schematic summarizing the temporal origin and neuronal expression of OPC temporal TFs. Top: the series of medulla neuroblast (NB) temporal windows, ordered and colored by neuroblast age (young to old), each labeled with its tTF combination. Below, the neuronal types produced from each window are shown, separated by Notch status (ON/OFF), with a few characteristic tsTFs for each temporal window (concentric genes) listed underneath. The bottom matrix (“tTF → tsTF”) shows the neuronal expression of the 6 temporal TFs that are also expressed as tsTFs (*hth*, *erm*, *ey*, *hbn*, *scro* and *D*) across these temporal windows, colored as Expressed, Partial/Subset, or Not Expressed. The summary is based on the trajectories presented in Fig. S18 as well as previously published *in situ* protein expression data (30). **b**, Coverage plot showing the accessibility of *erm* locus at P0. NBs were subclustered into their respective temporal windows, and 4 representative neurons were selected based on presence or absence of *erm* expression. Blue indicates the NBs expressing *erm*, and red indicates the neurons that express *erm* as a tsTF. Boxes indicate putative enhancer regions for the respective group, colored accordingly. **c-d**, Hierarchical cluster trees plotted for the indicated P0 clusters based on the top 10 components of (**c**) the PCA reduction, calculated on SCT residuals of the RNA data, and (**d**) the LSI reduction of the ATAC data. The first LSI component correlates with sequencing depth and is always ignored. See SI Methods regarding the selection of “young neurons” based on P0 clustering. NE: Neuroepithelia; NB: Neuroblasts; GMC: ganglion mother cells.

To test this hypothesis, we first calculated the correlations between gene expression and nearby enhancer accessibility (*i.e.,* peak-gene linkages), separately for the progenitor cells (NB+GMC) and the postmitotic neurons of OPC at P0 (**Table S4**) and then compared the top 5 peaks (by absolute z-score) linked to each of these 6 tTF genes in both groups. Indeed, we found that none of the top-correlating enhancers were common between progenitors and neurons for any of these genes except *scro*, for which the accessibility of its proximal promoter region (but only that region) correlated strongly with its expression in both groups. We then directly examined chromatin accessibility in and around these tTF loci, both in NBs at the corresponding temporal windows, and in a selection of postmitotic neurons. We highlighted some of the distinct putative NB and neuronal enhancers for *erm* (**Fig. 5b**), *ey* and *D* (**Fig. S16d-e**), whose accessibility correlated with expression of the gene in either NBs or neurons, but never both. In most cases, neuronal enhancers were closed in NBs and vice versa. Two FlyLight fragments (*R9D04* and *R9D08*) that overlap with some of the *erm* neuronal enhancers (**Fig. 5b**) were indeed active both in the adult medulla as well as in some OPC neurons (but not progenitors) at late L3 (**Fig. S17h-i**).

Mi9 neurons (**Fig. 4b**, red ellipse) represent a particularly interesting case: they are produced during a temporal window when *erm* and *ey* expression overlap in NBs and GMCs and they also express *erm* (but not *ey*) as postmitotic neurons (**Fig. 5a-b**). Thus, Erm expression in Mi9 would appear to be “maintained” from progenitors; yet accessibility of nearby enhancers suggests that *erm* expression is instead reactivated during neuronal commitment by cis-regulatory elements distinct from those that regulate it in NBs. **Figure 5b** further suggests that N^OFF^ Mi9 and N^ON^ Tm20 neurons, which unlike Mi9 do not originate from the Erm temporal windows (**Fig. 5a**), may also be using different (postmitotic) enhancers to express *erm*.

Finally, we built hierarchical trees of OPC NE, NB, GMC clusters, together with N^ON^ and N^OFF^ neurons immediately after their terminal division at P0 (i.e. cells that are near the bases of branches in **Fig. 4a-b**, see SI Methods). We calculated their similarities based on the top 10 components of PCA (for the RNA modality, **Fig. 5c**) or LSI (for the ATAC modality, **Fig. 5d**) reductions, which should represent the largest sources of variation among these cells. In both modalities, the GMC cluster was much closer to the NB cluster as compared to neurons. In the ATAC modality (**Fig. 5d**), even the N^ON^ and N^OFF^ neurons were immediately less similar to each other compared to the distance (*i.e.,* branch length) between the NB and GMC clusters. Combined with the observations above, our results strongly suggest that the chromatin landscape undergoes a dramatic transformation during the first few hours of a neuron’s life after a GMC’s terminal division. Even the genes that are continuously expressed from progenitors appear to switch their enhancers upon neuronal commitment, allowing them to be re-utilized independently from their regulation in progenitors.

We posit that this brief window during which Notch signaling is active and temporal TFs inherited from neuroblasts may still be available at the protein level is a particularly amenable regulatory state (*i.e.,* critical period) that enables each neuron to decode its developmental history into a stably maintained terminal selector code. In this model, cell-type identity is not directly ‘inherited’ from the progenitors but rather ‘computed’ (based on lineage inputs) by a dedicated gene regulatory network in newborn neurons. Our dataset is a unique resource to systematically investigate this process and understand how the joint actions of various patterning mechanisms instruct the generation of hundreds of cell types from neural stem cells.

## Discussion

By sequencing nearly a quarter million single cells in two modalities, we resolved over 250 cell types (233 neurons) of the *Drosophila* optic lobes consistently across development, the highest resolution of any single-cell atlas that includes this brain region (23, 30, 40, 42, 64, 65), and one of the most granular descriptions of any complex nervous system to date. While this is a significant improvement, only 32 of the 206 clusters of our previous scRNA-seq atlas (23) were found here to contain multiple cell types, generally owing to the rarity of the cell types involved or subtle differences between very similar neurons. Notably, in all cases where we resolved molecularly distinct subtypes among morphologically homogeneous neurons (albeit subtly), these fell within some of the most abundant optic lobe cell types (Dm2, Tm4, Tm6, Tm9), implying that comparable subdivisions within rarer types may remain unresolved. Overall, scRNA-seq characterizations of the optic lobe (23, 42) were already near saturation; our dataset extends this resolution to the ATAC modality and should contain independent clusters for all cell types present in at least ∼20 cells per optic lobe. This threshold explains the discrepancy with the >700 cell types recently defined by EM connectomics (17): >500 of those were found in <20 copies, whereas the top 230 most abundant types with >20 copies account for 97% of all optic lobe neurons. The same difficulty in resolving rare types mirrors the continued challenges for scRNA-seq of the central brain (66, 67), where neuronal types are rarely present in more than 20 copies per brain; we predict that 1–2 million single cells per stage will be required to reach a comparable resolution there.

The tens of thousands of putative neuronal enhancers we identified here are mostly type-specific yet share common sequence features. The presence of pan-neuronal TF motifs (even in type-specific enhancers), and of binding sites for Ttk that likely repress them in non-neuronal cells could keep these enhancers inactive elsewhere in the body, consistent with the neuroendocrine tumors (that upregulate some neuronal TFs) induced by the removal of *ttk* from intestinal stem cells (68). This may enable neuronal enhancers to evolve more freely without disrupting the expression of their associated genes in other tissues. The prevalence of Klu and GAF motifs, even though these TFs are expressed in NBs but not neurons, suggests that progenitors both pioneer these regions for later use and prevent their premature activation (51). Finally, the enrichment of homeodomain binding sites, which are overrepresented among tsTFs (69), in neuronal DARs but not pan-neuronal enhancers supports our previous conclusion that tsTFs are primarily responsible for neuron type-specific gene regulation (36).

Importantly, we show that tsTFs orchestrate a very dynamic gene regulatory program as neurons differentiate. Despite their continuous expression, many (if not most) tsTF targets are stage-specific, largely through their co-operation with ecdysone-regulated TFs, as previously observed in the lamina neurons (54). The unique, dynamic gene expression program of each neuron therefore requires the combinatorial action of at least two regulatory mechanisms: one stable across time but variable among cell types (tsTFs), and the other uniform among cell types but variable through time (hormones). These two mechanisms are best understood as orthogonal axes of regulatory variation (rather than independent or parallel pathways) that converge on shared enhancers: the expression of a given stage-specific effector often requires the simultaneous input of both a cell-type-specific TF combination and the appropriate hormone-responsive TF acting on the same regulatory element.

Our computationally inferred GRN models also reinforce the central role of tsTFs in regulating neuronal DEGs and provide support for three key features of terminal selectors that were originally proposed in *C. elegans* (33). First, tsTFs can only act in combinations: we found little overlap among the genes regulated by the same tsTFs across different metaclusters. Thus, target specificity is not conferred by individual tsTFs but is strictly dependent on the other TFs expressed in each cell-type. This combinatorial requirement is in fact a property of all regulators, including pan-neuronal TFs; we therefore argue that any notion of a single “master regulator” for a given neuronal feature is likely to be misleading, as all such interactions are context-dependent at some level. Second, tsTFs directly regulate at least some of their neuronal effector genes. All interactions inferred by SCENIC+ (59) are supported by cis-regulatory evidence, *i.e.* enrichment of the matching TF motifs in a peak linked to the target gene by accessibility-expression correlation. Nevertheless, tsTFs also commonly regulate other TFs, including other tsTFs, implying that a substantial fraction of their downstream effects may be indirect. Third, tsTFs do not specialize in distinct phenotypic features such as morphology or electrophysiology, likely as a consequence of their combinatorial nature. Consistently, the predicted targets of most tsTFs were enriched for all functional classes of cluster marker genes (**Fig. S14**). This does not imply that every effector in a neuron is jointly regulated by all tsTFs in that neuron. As we previously proposed (36, 37), our results are more consistent with the effector genes being controlled by different permutations of available tsTFs in each neuron, without any phenotypic specialization, but any given effector is not regulated by all tsTFs in that neuron.

We previously defined putative tsTFs in optic lobe neurons based on their continuous expression in specific cell types (36). This definition of “terminal selector” is useful because neuronal types can be interconverted simply by modifying these codes (disregarding other DE TFs that are not continuously expressed). This implies that these combinations contain within them the ‘minimal code’ of TFs required to specify each type. However, many of the same TFs are also transiently expressed in other cell types, where they are not tsTFs by this definition. This does not imply that these TFs cannot have important functions in those neurons: We identified regulons for some TFs in metaclusters where they would not be considered tsTFs (but they are in others). Such TFs may appear to have more phenotypic specialization if they are only expressed at specific stages in a given cell type, where their effects would be accordingly limited.

The *Drosophila* optic lobe is a powerful system for dissecting how these tsTF combinations are specified. In addition to the putative tsTF codes in every neuron (36), the spatial, temporal and Notch origin of most OPC neurons have been mapped (30, 37), as well as the correlations between these origins and the tsTF expression patterns (37). A particular strength is our ability to link neuronal differentiation trajectories at P0 to NB/GMC temporal windows (**Fig. 4a-b** and **Fig. S18**), enabled by the continuous production of same cell types over 2-3 days by sequentially generated NBs, a unique feature of the visual system. This assumption could be complicated by the recent description of a fourth patterning mechanism acting within the OPC NE (25). This long-range temporal patterning mediated by the RNA-binding proteins Imp and Syp (as in central brain NBs) implies that different neurons could be produced during early vs. late stages of neurogenesis from different NBs, even when they share the same spatial domain, temporal window and Notch status.

This ability to link neuronal lineages to specific temporal windows revealed that NB temporal TFs do not exert their effects on neuronal type identity by being maintained in neurons that originate from their respective windows. Most tTFs are not maintained in differentiated neurons at all (30); even when the same gene functions as both a tTF and a tsTF, there is no correlation between the temporal window of origin and expression of these TFs in neurons (with the exceptions of *hth* and *scro*). These genes are almost always associated with different open chromatin regions in NBs vs. neurons, indicating independent regulation even in cases of apparent “maintenance”. This phenomenon is not restricted to these tTF genes: the genome-wide chromatin landscape undergoes a dramatic transformation during the first few hours of a neuron’s life after a GMC’s terminal division. Together, these results effectively sever direct continuity of neuronal gene regulatory programs with those of their stem cell ancestors and suggest that “mitotic bookmarking” (51, 70) by tTFs in tsTF loci is not a commonly used mechanism, given the highly divergent enhancer accessibility in stem cells and neurons. They also argue that temporal identity is not directly inherited in neurons, but rather integrated with spatial origin and Notch status during a specific critical period to instruct the expression of the unique combination of tsTFs in each neuron. The cis-regulatory logic underlying this integration remains difficult to decipher due to the highly transient nature of these regulatory events, which will be a subject of future studies.

Overall, our findings support the notion that tsTFs function as ultimate integrators of developmental history in newborn neurons, which appear to occupy a special regulatory state that enables them to decode that history into a stable terminal selector combination. These combinations then remain locked in for the subsequent stages of differentiation and instruct all type-specific gene expression in neurons. Our findings also clarify a feature that has been largely implicit in earlier discussions of this framework: being a “terminal selector” is best understood as a property of a TF/cell-type pair rather than of a TF gene. The same factor may serve as a continuously expressed tsTF in one neuronal type and as a transient, “subroutine” factor in another. And within a single cell type, the targets of a given tsTF are often regulated by shifting permutations of partner TFs, with downstream programs substantially shaped by developmental stage and the broader hormonal and signaling milieu.

## Materials and Methods

In order to perform simultaneous RNA and ATAC sequencing on very large numbers of developing optic lobe cells, we followed a genetic multiplexing strategy (42) based on the wild-type strains of the Drosophila Genetic Reference Panel (71) (DGRP). We crossed males from 12 different DGRP lines (40, 189, 320, 382, 383, 391, 441, 406, 492, 505, 852 and 897) to Canton S virgins to acquire F1 hybrids. We selected female white pupae (P0) and reared these to the required stage at 25°C; and for adult dissections, we only used 0-day old (<24h after eclosion) flies. For each of the 4 stages we sequenced (P0, P24, P48 and Adult), we dissected a single fly from each of these crosses, which were then processed together for all subsequent steps of nuclei isolation and sequencing. Natural genetic variation among the DGRP strains (72) enables *post hoc* assignment of each sequenced cell to its origin strain based on its unique SNPs (**Fig. S2**). This approach provides two chief advantages: i) extremely robust identification of doublets, allowing us to ‘overload’ each microfluidic well, reducing per cell library preparation costs; ii) preservation of biological replicates, i.e. for all analyses that are performed at the cluster level rather than single cells, cells belonging to the same DGRP line at each stage (representing a single fly, except for the “P48-1” library which was a separately performed pilot experiment that contained the same dgrp strains) can be ‘pseudobulked’ separately, increasing statistical power. All relevant QC metrics per stage, library, dgrp strain and cell type are presented in the Supplementary Figures. The details of nuclei isolation, library preparation, sequencing, and all data analysis are provided in the SI Methods.

Detailed procedures for all other experiments we performed (*i.e.,* for cell-type annotations, in **Figs. S8-9**) and their analysis, as well as the source details for all *Drosophila* strains, antibodies and other reagents are also provided in the SI Methods.

## Acknowledgements

We would like to thank all members of the Desplan and Özel Labs for helpful discussions, and Filippo Riva, Jay Unruh, Tom Kleist and Jason Morrison for technical assistance. We thank Robb Krumlauf, Filipe Pinto Teixeira Sousa, Nikos Konstantinides and Saumya Jain for critical reading of the manuscript. This work was supported by the Stowers Institute for Medical Research, K99/R00-NS125117 and a Klingenstein-Simons Neuroscience Fellowship to M.N.Ö., R01-EY017916 and R01-EY013010 to C.D., and R21-NS140885 to C.D. and M.N.Ö.

## Author Contributions

M.N.Ö. and C.D. conceived the project. M.N.Ö. performed all single-cell sequencing experiments. M.N.Ö., M.T., Y.C.C., O.B.T., R.C., R.R., H.H. and H.L. analyzed the sequencing data. A.T., R.C., C.L. and Y.-C.D.C. performed all other experiments and analyzed the data. M.N.Ö. and M.T. wrote the manuscript. All authors edited the manuscript.

## Data Availability

Raw and processed multiome data have been uploaded to GEO: GSE305940 and will be publicly available upon acceptance for publication. All other original data underlying this manuscript can also be accessed after publication from the Stowers Original Data Repository: https://www.stowers.org/research/publications/LIBPB-2595

## Supplementary Table Legends

**Table S1: Cluster Annotations**

Summary table of all 259 clusters, including previous and updated cluster identities, cell type annotations, metacluster assignments, temporal and spatial origins, Notch status, and any relevant comments.

**Table S2: Cluster markers (RNA modality) at P24, P48 and Adult stages**

Table of RNA gene markers enriched in each cluster at P24, P48, and Adult stages. Markers were identified using Seurat’s *FindAllMarkers* on stage-specific subsets of the Multiome dataset, including only positively enriched genes (log2FC > 1, adjusted p < 0.01) and excluding non-neuronal clusters. See *Methods* for more details.

**Table S3: Cluster markers (ATAC modality) at P24, P48 and Adult stages**

Table of ATAC peak markers enriched in each cluster at P24, P48, and Adult stages. Markers were identified using Seurat’s *FindAllMarkers* on stage-specific subsets of the Multiome dataset, including only positively enriched genes (log2FC > 1, adjusted p < 0.01) and excluding non-neuronal clusters. See *Methods* for more details.

**Table S4: Gene-enhancer links at P0, P24, P48, and Adult stages**

Table of gene–enhancer links identified across developmental stages, showing peak associations identified by correlation of gene expression with nearby accessible regions and highlighting the dynamic usage of enhancers across P0 (OPCprogenitor), P0 (OPCneuronal), P0 (all), P24, P48, and Adult stages, as well as all stages combined. Links were identified using Seurat’s *LinkPeaks* function, with a search window of 50kb upstream and downstream of the TSS. See *Methods* for more details.

**Table S5: Motif enrichment analysis of neuronal enhancers**

Table summarizing the motifs enriched in consistent and stage-specific neuronal enhancer peaks. *De novo* motif discovery was performed with the MEME suite’s STREME tool using sequences derived from DARs identified in **Table S3**, and candidate regulators were assigned by comparing discovered motifs against the CisBP-*Drosophila* database with Tomtom. See *Methods* for more details.

**Table S6: SCENIC+ networks**

Table containing 28 metacluster-specific gene regulatory networks (GRNs) inferred with SCENIC+ using multiome and supporting scRNA-seq datasets from P24, P48, and Adult stages (tabs). Some related metaclusters were merged prior to network inference to improve interpretability and statistical power. See *Methods* for more details.

## SI Methods

### Single-cell multiome sequencing of whole optic lobes

For nuclei isolation from the adult and developing *Drosophila* optic lobes for multiome sequencing, we prepared Wash Buffer to include: 10 mM Tris-HCl (pH 7.4), 10 mM NaCl, 3 mM MgCl_2_, 0.1% Tween-20, 1% bovine serum albumin (BSA), 1 mM dithiothreitol (DTT) and 1 U/µl of Protector RNase inhibitor (Sigma) in nuclease-free water. Nuclei lysis buffer was prepared to include all ingredients of the wash buffer, with the addition of 0.1% Nonidet P40 substitute and 0.01% Digitonin. 20x Nuclei buffer (10x Genomics) was diluted to 1x in nuclease-free water and supplemented with 1 mM DTT and 1 U/µl RNase inhibitor. All solutions were maintained on ice throughout the protocol.

For each experiment, we selected female white pupae (P0) and reared these to the required stage at 25°C. For the adult dissections, we only used 0-day old (<24h after eclosion) flies. The optic lobes were dissected and maintained in ice-cold Schneider’s media until lysis. They were washed 3 times with DPBS before being transferred into a dounce homogenizer (pre-wet pestles with wash buffer) containing 500 µl lysis buffer. The optic lobes in lysis buffer were incubated on ice for 5 minutes, homogenized with 25 strokes with pestle A, followed by another 5-minute incubation on ice, and 30 strokes with pestle B. Without fully removing the pestle B from the homogenizer, both the pestle and the side walls of the homogenizer were then rinsed with 1 mL wash buffer. The entire volume was transferred into a pre-cooled 2 mL Biopur tube. We also recommend mixing the lysis solution using a wide-bore tip in the middle of both incubation steps. Diluted lysis solution was filtered with 40 µm Flowmi cell strainers (in two batches) directly onto a 20 µm PluriStrainer (pre-wet with wash buffer) placed on a second pre-cooled 2 ml tube. After filtering, nuclei were centrifuged (at 4°C) for 5 minutes at 800 rcf using a swing bucket rotor. Excess buffer was discarded without disrupting the pellet, another 1 mL wash buffer was added. After pipette mixing to wash the nuclei, they were pelleted again by another 5 minutes of centrifugation at 800 rcf; excess buffer was discarded, leaving no more than 10 µl of wash buffer. It is critical to minimize the amount of buffer left in the tube at this step, as the BSA in lysis and wash buffers may inhibit subsequent Tn5 transposition. Finally, the pellet was resuspended in diluted nuclei buffer (minimum 50 µl, 100 µl is recommended if the amount of input material allows for sufficient concentration) and were filtered through a 10 µm mini-PluriStrainer (pre-wet with nuclei buffer) into a 1.5 mL DNA LoBind tube. In order to assess lysis efficiency and to determine the nuclei concentration we stained a small aliquot with Calcein AM and EthD-III (Live/Dead assay) and examined/counted them in a hemocytometer using an epifluorescence microscope.

Next, we used Chromium Next GEM Single Cell Multiome ATAC + Gene Expression kits (10x Genomics) and followed the manufacturer’s instructions for producing ATAC and gene expression (RNA) libraries for sequencing. Briefly, the nuclei were subjected to Tn5 transposition for 1 hour at 37°C and were then immediately loaded into a Chip J for generation of nanoliter-scale gel bead-in-emulsions (GEMs) using Chromium Controller. Following reverse-transcription, cleanup and pre-amplification PCR, the samples were split into two for the ATAC library PCR (9 cycles) and for cDNA amplification (7 cycles) followed by fragmentation, adaptor ligation and sample index PCR for the RNA library (11 cycles). We used 20 µl of the amplified cDNA for RNA library preparation (even though 10 µl is normally recommended), and the final resuspension of the ATAC libraries was done using 40 µl of EB buffer, even though 20 µl is normally recommended. We produced 5 different multiomic libraries (technical replicates) for the P48 stage, and 4 libraries for other stages (P0, P24 and Adult), aiming for ∼20,000 cells per library.

The libraries were subjected to paired-end Illumina sequencing using NovaSeq 6000: 50×8×24×49 cycles for ATAC libraries and 28×10×10×90 cycles for RNA libraries. We sequenced all RNA libraries to 80-90% saturation (as reported by ‘Percent duplicates’ in CellRanger), and all ATAC libraries to 75-80% saturation.

### Processing and quality control of the multiome data

The multiome data was processed using CellRanger-ARC v2.0.2. After library demultiplexing (mkfastq), the reads were aligned and counted using the *D. melanogaster* genome annotation version BDGP6.88 (Ensembl). The ATAC reads were initially counted based on a predefined set of 128,464 genomic ranges (75) (ctx) which cover 80%, i.e. nearly all non-coding regions, of the *Drosophila* genome. Multimodal cell-calling algorithm of CellRanger-ARC failed on our data, likely due to high background in the ATAC modality. For each library, we thus also ran (base) CellRanger v7 on the RNA modality alone, which identified 16-20,000 barcodes corresponding to cells per library (before filtering) for stages P0, P48 and Adult, and 23-25,000 cells per library for stage P24.

Gene (RNA) and peak (ATAC) count matrices for the cell barcodes were loaded into R and further processed using the packages Seurat (38) v4.1.4 and Signac (39) v1.11.0. We first filtered out any barcodes with less than 300 (ATAC) fragments in peak regions (ctx), or less than 200 genes detected (RNA). Additionally, we removed the barcodes with the proportion of mitochondrial (RNA) reads exceeding 20% (P0 and P24), 30% (P48) or 40% (Adult). We observed that our data, and especially later stages, contain unusually large amounts of mitochondrial RNA (median: 12% in Adult cells). While this is typically indicative of low transcriptome quality, we also observed that at relatively lower regimes (up to ∼20%) mitochondrial RNA proportion in our data in fact correlates positively with the total RNA counts (UMIs). After filtering, we removed all mitochondrial genes from the gene/cell matrices to prevent these reads from skewing the depth normalization of single cells. Ribosomal RNA content was also unusually high in these data, which is not directly detectable in Seurat but causes low rates (55-65%) of “Reads mapped confidently to transcriptome” as reported by CellRanger (due to rRNA gene duplications). We concluded that these issues, other than reducing the number of usable reads, do not affect any of the downstream analyses. While the underlying reasons remain unclear, we speculate that incomplete lysis of plasma membrane in some cells may have trapped ribosomes and mitochondria together with the nuclei we sequenced. We additionally removed the barcodes with Nucleosome Signal (the ratio of mononucleosomic fragments to nucleosome-free fragments) of more than 1, or with TSS (transcription start site) Enrichment score of less than 1.5 in the ATAC modality.

We next used demuxlet (76) to identify and remove doublets based on DGRP SNPs detected in each cell (RNA modality). To assign individual cells from pooled F1 crosses to their DGRP parental backgrounds, we adapted a previous approach using popscle with helper scripts from popscle_helper_tools. Since we profiled F1 hybrids from DGRP × Canton-S crosses to reduce possible genotype-specific effects, the heterozygous samples required us to focus exclusively on DGRP-specific variants for genotype assignment. First, we lifted over the DGRP2 variant database to the dm6 reference genome using LiftoverVcf provided by Picard v4.0.2.0 with the chain file provided by UCSC Genome Browser: https://hgdownload.soe.ucsc.edu/goldenPath/dm3/liftOver/dm3ToDm6.over.chain.gz

We identified informative SNPs by filtering DGRP variants using custom functions from popscle_helper_tools. We retained only positions that were: (1) homozygous for the alternative allele in at most 2 of the selected DGRP lines, (2) not missing genotype information in any line, (3) not homozygous reference in all lines, and (4) not heterozygous in any line. VCF processing was performed using bcftools v1.14 and vcftools v0.1.16.

To exclude Canton-S-specific variants from our analysis, we generated a Canton-S variant mask by calling variants from a published Canton-S optic lobe developmental single-cell transcriptomic atlas (23). We performed variant calling using bcftools mpileup and bcftools call v1.14 on the aggregated BAM files from this atlas against the dm6 reference genome. We then masked all SNP positions overlapping with these Canton-S variants using bedtools v2.29.2, as these positions would be uninformative for distinguishing between F1s from different DGRP backgrounds.

To ensure reliable genotype assignment, we generated coverage tracks using deepTools v3.5.0 and filtered genomic regions requiring a minimum read depth of 10. To minimize the impact of ambient RNA contamination on genotype assignment, we quantified gene expression using HTSeq v0.13.5 and identified the top 100 genes by expression level. These highly expressed regions were excluded from SNP calling using the bedtools *complement* and *intersect* operations under the assumption that highly expressed transcripts are more likely to contribute to the ambient RNA and carry mixed genotype signals. Cell barcodes passing quality control were used to filter the BAM files using samtools v1.14 and popscle_helper_tools scripts, retaining only reads from valid cells. The filtered and sorted BAM files were processed through popscle dsc-pileup to generate allele count matrices at informative SNP positions. For demultiplexing, we converted homozygous genotypes in the DGRP database (1/1) to heterozygous calls (1/0) in the VCF file to reflect the expected F1 genotypes. Finally, we performed demultiplexing using popscle demuxlet with the genotype-aware mode, testing multiple alpha parameters (0 and 0.5) to optimize the assignment of cells to their DGRP parental backgrounds while identifying potential doublets. The doublet rates ranged from ∼15% (P48) to ∼24% (other stages) per library. A low number of cells were misclassified to DGRP line_307 and line_508. When using the DGRP assignments for downstream analysis (e.g. creation of metacells, outlined below), these two assignments (and the cells assigned to them) are dropped from the analysis.

Using Seurat v4.1.4 and Signac v1.10.0, barcodes representing singlets and passing all filters have been merged into a single Seurat (.rds) object for each stage. ATAC data was normalized with *RunTFIDF* using default parameters. After feature selection with *FindTopFeatures*, with default parameters, we used latent semantic indexing (LSI) to acquire 150 reduced dimensions using the function *RunSVD*. As described in the Signac tutorial “Analyzing PBMC scATAC-seq” (https://stuartlab.org/signac/articles/pbmc_vignette.html), we observed that the first LSI dimension always correlated very strongly with sequencing depth and thus was excluded from all further analyses that use the LSI reduction. For the RNA data, after selection of top 2000 variable genes using *FindVariableFeatures*, we applied both standard log normalization using *LogNormalize* as well as *SCTransform*, a GLM-based method that was proposed to provide better variance stabilization especially for highly expressed genes (77). We used the SCTransformed data for dimensional reduction of the RNA modality using *RunPCA*, (npcs=150) and all downstream steps that utilize the PCA reduction (below). Whereas for plotting gene expression values or calculating differentially expressed genes between clusters, we used the log-normalized expression values.

### Classification of the multiome data

We independently took supervised and unsupervised approaches to assign multiomic barcodes to specific cell types. The supervised classification was performed using an artificial neural network model that we had trained on the cluster identities of our previously published scRNA-seq dataset (23). This is a multi-task network that accepts expression data from different stages through independent input layers based on the top marker genes at each stage. We thus used the P15 markers for classifying the P0 data, P30 markers for the P24 data, P50 markers for the P48 data, and the Adult markers for the Adult multiome data. These models use the log-normalized expression values of the corresponding marker genes to automatically assign each cell to one of the 206 clusters in our published atlas.

For unsupervised classifications (clustering), we used the Weighted Nearest Neighbor (WNN) algorithm (38) from Seurat v4.1.4, which uses both PCA and LSI reductions to build a joint nearest neighbors graph taking advantage of both modalities using *FindMultiModalNeighbors*, (reduction.list=list(”pca”, “lsi”), dims.list = list(1:150, 2:150)). These neighbor graphs were used both for calculating UMAP reductions with *RunUMAP* (nn.name = “weighted.nn”) for visualization (**Fig. 1**), and for unsupervised clustering of each stage using *FindClusters (*graph.name=”wsnn”, resolution=7). The highest number of clusters, as expected, were achieved at the P48 stage. Due to the continuous trajectory structure of the P0 dataset (**Figs. 1d, 4a-b**), clustering results are only useful for sub-setting the data into specific trajectories for further analysis. Thus, at this stage, we intentionally used a lower clustering resolution of 2.

We performed a comprehensive comparison of the clustering results at P48 and the classifications of the same cells by the P50 neural network. WNN clustering identified subgroups within the following clusters of our published atlas(23): 3 (Photoreceptors), 20, 21, 23, 24, 28, 39, 57 (LC25), 58, 65, 67, 79 (LC10b), 83, 87 (LC20), 91 (Mt8), 100 (LC16), 101, 128 (Dm2), 129 (Mi21), 136 (Dm11), 147 (LC4), 149 (LC22/LPLC4), 158 (Pm3b/Tm23), 159 (TmY15/17), 162, 170, 178, 184, 222 (LC6), 223 (TEv). 2-3 subclusters were identified in each of these clusters, with the exception of cluster 24 (6 subclusters) and cluster 100 (5 subclusters), which we have already shown to contain LC15,16,24,25 neurons (24) (and one more, unknown, neuron). Heterogeneities within the old clusters 129, 158, 159 were also described in other recent studies (25, 26). By examining the differentially expressed genes between the subclusters, we validated that all of these splits represented biologically significant (i.e. not technical) subgroups. Given the clearly superior cell-type resolution of these multiome data over our published scRNA-seq data (23), we decided to abandon our previous classifications altogether and build a new reference based on the multiome clusters. In cases where these clusters were already annotated to specific cell-types, these annotations were re-evaluated based on the markers of each subcluster and accordingly adjusted (detailed in **Table S1**).

WNN clustering in addition identified subgroups within the (old) clusters 62 (Tm6), 85, 102, 125 (Tm4) and 181. Divisions within the Tm4 and Tm6 clusters represented the dorsoventral subtypes we previously described for these cell-types (23). However, the distinctions between these subtypes are very subtle and hard to resolve, especially in stages other than P48; we thus decided to keep Tm4 and Tm6 as single clusters. Clusters 85 and 102 were highly heterogeneous mixtures of central brain derived neurons; we thus merged all multiome clusters corresponding to these (old) clusters into one cluster, 141. Similarly, the division of old cluster 181, while likely meaningful, proved unreliable and was thus merged back into a single cluster (195). Lastly, through our previous (unpublished) sub-clustering attempts, we have been aware that (old) clusters 59, 95 and 96 also contained subgroups, although these were not resolved by the WNN clustering of the entire P48 dataset. We manually split these clusters (into two) by individually sub-clustering them.

Using the (corrected) clusters of the P48 multiome data as reference, we next classified the P24 and Adult datasets. We implemented a label transfer procedure to take advantage of both modalities for this purpose. First, we determined the differentially expressed genes using the *FindAllMarkers* function (assay=’RNA’, only.pos= TRUE, logfc.threshold= 0.5, max.cells.per.ident= 500) and differentially accessible regions (assay= ‘ATAC’, only.pos= TRUE, test.use= ‘LR’, latent.vars= ‘atac_peak_region_fragments’, logfc.threshold= 0.5, min.pct= 0.05, max.cells.per.ident= 500) for all clusters at P48. For the RNA modality, we kept only the ‘top100’ (by logFC) markers for each cluster. Second, we set the variable features of the ATAC assay to the differentially accessible regions and recalculated the LSI reduction using *RunSVD* and the weighted nearest neighbors function from Seurat *FindMultiModalNeighbors* (reduction.list= list(”pca”, “lsi”), dims.list= list(1:150, 2:150), k.nn= 31) based on this. We then transferred the labels from the P48 dataset to the P24 and Adult datasets based first on the ATAC modality, using the functions *FindTransferAnchors* (reference.reduction= “lsi”, reference.neighbors= “weighted.nn”, reduction= “lsiproject”, dims=2:150) and then *TransferData* (weight.reduction= “lsiproject”). Independently, we also transferred labels based on the RNA modality: *FindTransferAnchors* (reference.assay= “SCT”, query.assay= “SCT”, reduction= “cca”, reference.neighbors= “weighted.nn”, features= top100, normalization.method= “SCT”, dims= 1:150) and *TransferData* (weight.reduction= “cca”). The predictions based on both modalities were largely in agreement; we resolved any inconsistencies based on the prediction score (i.e. confidence) calculated by Seurat: if for a given cell different identities were predicted by RNA and ATAC label transfer, the prediction with the higher score was kept (in case of ties, ATAC-based predictions were kept). Finally, we compared the transferred labels to results of the unsupervised clustering we performed at Adult and P24 stages and validated that no additional subdivisions were present. The same procedure was then used for transferring labels from the P24 to P0 dataset. However, since the P0 data contains progenitor cells that are not present in the later stages, using known markers, we separated out the cells corresponding to neuroepithelia (NE), neuroblasts (NB), ganglion mother cells (GMCs) of OPC and IPC, as well as the lamina precursor cells (LPC) based on the unsupervised clustering we performed at P0 (**Fig. 4**).

### Multi-task neural network classifier training and prediction

We trained stage-specific neural network classifiers (Adult, P24, and P48) on log-normalized gene expression profiles, with cluster identities encoded using one-hot encoding. Each model consisted of an input layer (dimension equal to the number of marker genes, top 10 by cluster, per stage), a dropout layer (20%), a dense hidden layer (200 units, ReLU activation, L2 regularization), an additional dropout layer (50%), and a softmax output layer predicting 259 cluster labels. Models were optimized using stochastic gradient descent with Nesterov momentum (learning rate = 0.05, momentum = 0.9), categorical cross-entropy loss, and balanced class weights. Learning rates were reduced stepwise every three epochs. Early stopping and model checkpointing were applied to prevent overfitting.

To encourage cross-stage generalization, models were trained sequentially by passing weights across stages in the order Adult → P24 → P48 and back to Adult, repeating for 14 runs of 30 epochs each. Training and validation splits were predefined for each dataset (Adult: 42,648; P24: 58,390; P48: 54,368 training samples). Performance was monitored at each stage, and the best-performing models (based on lowest validation loss and highest accuracy) were retained for downstream analysis.

For prediction on new datasets, expression matrices were normalized by subtracting the stage-specific mean expression values used during training. Predictions were generated using the trained stage-appropriate model. To estimate confidence, we applied Monte Carlo dropout during inference: 500 stochastic forward passes were performed with dropout layers active, and predicted class labels were recorded for each pass. Confidence was calculated as the proportion of bootstrap predictions consistent with the model’s maximum-probability class assignment. Final outputs included the predicted cluster label, softmax score, and bootstrap-derived confidence score for each cell.

The trained classifiers enable consistent reclassification of previously published scRNA-seq datasets of the optic lobe in alignment with our clustering. All trained models, encoders, example scripts, and prediction utilities are available at: https://github.com/ktreese03/single_cell_classifier

### Identification of cluster markers in RNA and ATAC modalities

To define stage-specific and consistent markers of neuronal clusters, we used the merged multiome dataset after removing 3 clusters of TE neurons (absent in adults), cluster 141 (heterogeneous), and all non-neuronal populations. For each developmental stage (P24, P48, and Adult), we identified differentially expressed genes (DEGs) using Seurat’s v4.1.4 *FindAllMarkers* function on the RNA assay with stringent thresholds (log₂FC > 1, adjusted *p* < 0.01, one-vs-all comparisons, maximum 500 cells per cluster). Genes were then categorized as stage-specific, shared between two stages, or common to all three. To identify stable markers, we further required that a gene be captured in the same cluster across stages, allowing ≤2 stage-specific absences to account for sparsity. Similarly, differentially accessible regions (DARs) were identified using the ATAC assay with *FindAllMarkers* (logistic regression test, latent variable = total ATAC counts, log₂FC > 1, adjusted *p* < 0.01, min.pct = 0.05, maximum 500 cells per cluster). The same filtering strategy was applied to remove low-quality clusters, and DARs were grouped as stage-specific, shared, or consistent across development. This approach yielded 2,256 neuronal DEGs and 53,637 DARs across P24, P48, and Adult, of which ∼30% of DEGs and ∼40% of DARs were consistent markers of at least one neuronal cluster across stages (**Tables S2-3).**

### Metaclustering of the multiome data

The very high diversity of cell types in our dataset (259 clusters, 233 of which are neurons) makes it useful (e.g. for peak calling, below) to have higher-order groupings of these clusters based on similarity, coined metaclusters. Since we have consistent cluster assignments of all cells across P24, P48 and Adult stages, metaclusters were determined for these 3 stages together and separately for the P0 stage.

For the later stages, we performed hierarchical clustering using Seurat v4.1.4 of our cell types at all 3 stages, using various similarity measures based on both modalities. The following procedure was then selected empirically as the resulting tree (**Fig. S7a**) best represented the known developmental relationships among optic lobe neurons: *FindAllMarkers* function was used on the RNA modality with default parameters to determine marker genes for all clusters (separately for neuronal and non-neuronal clusters). Only the genes found as markers in all 3 stages were used as features (rather than the default variable genes) to calculate PCA reduction on the RNA modality using *RunPCA* on the P24 stage. We then ran the *BuildClusterTree* function on these P24 data using the first 100 PCs (for neurons) or the first 20 PCs (for non-neurons) to build hierarchical trees. We manually selected nodes based on known cell type relationships in the literature, to group neuronal clusters into 37 metaclusters, and the non-neuronal clusters into 5 metaclusters (**Fig. S7a-b**). Note that the cells belonging to a given cluster are thus assigned to the same metacluster across P24, P48 and Adult stages.

For the P0 stage, we used the unsupervised clustering performed at this stage (described above) to determine and merge similar clusters into metaclusters (**Fig. S7d**). These were largely decided manually by examining cluster trees using the function (assay = ‘RNA’, reduction = ‘pca’, dims = 1:50) as well as their organization within the UMAP trajectories (**Fig. S7d**). 127 P0 clusters were thus grouped into 19 metaclusters.

### Peak calling

We used the MACS2 (43) package via Signac’s (v1.13.0) *CallPeaks* function to perform peak calling independently on the ATAC reads belonging to the 61 metaclusters we determined above. We note that the paired-end (BEDPE) mode of MACS2 is not suitable for ATAC-seq, where the fragment ends (i.e. cut sites) represent the real signal rather than the fragment centers (as in ChIP-seq). Many workflows thus recommend the single-end (BED) mode with the parameters extsize= 200 (or similar) and shift=-extsize/2, which is valid but effectively ignores half the cut sites present in the fragments file. We thus first converted the “atac_fragments” file from CellRanger-ARC into an “atac_cuts” file, whereby cut sites (the edge positions of each fragment) are re-written as different fragments (i.e. different lines) of length 1 bp. In the merged Seurat (.rds) Multiome object, this “atac_cuts” file was then set as the fragments path and MACS2 was run through the Signac wrapper *CallPeaks* with the following parameters: effective.genome.size= 1.5e+08, additional.args= “--keep-dup all --tsize 100”, group.by= “metacluster.idents”, combine.peaks= FALSE. Metacluster specific peaksets were then saved as separate .bed files. For P24-Adult peak sets (M1-M42), the average (mean) peak length was ∼572bp and the median number of peaks in each set was ∼24,433. For the P0 peak sets (M43-M61), the average (mean) peak length was ∼578bp and the median number of peaks in each set was 21,085. Among all peak sets (M1-M61), the average (mean) peak length was ∼574bp and the median number of peaks in each set was 24,009. Peaks overlapping known problematic genomic regions (blacklist regions) were removed prior to further analysis.

Finally, we attempted two methods to generate a consensus peak set from the P0-Adult peaks: First, we converted the metacluster-specific peaks into a pyranges dictionary and used the *get_consensus_peaks* function (parameters: peak_half_width=250) of the pyCistopic v2.0 package from the SCENIC+ framework (59) to obtain 75,502 fixed-length (500 bp) consensus peaks. Second, we attempted to adapt ArchR’s (78) iterative peak merging method within their *addReproduciblePeakSet* function. Specifically, we utilized the second step, where ArchR merges peak sets into a union peak set. All default parameters from ArchR’s method were retained. To identify reproducible peaks within each group, individual peak summits were extended ±250 bp, and non-overlapping peaks were selected based on peak score. Reproducibility for each peak was calculated as the proportion of replicates (metaclusters) in which the peak was present, and peaks were retained if they were present in at least 50% of the replicates. Reproducible peaks were annotated with distance to the nearest gene, transcription start site (TSS), GC content, and peak classification (promoter, intronic, exonic, or distal).

Following identification of group-specific reproducible peaks, ArchR’s *addReproduciblePeakSet* function was used to merge these peak sets into a single union peak set. During this step, peaks were iteratively merged across groups to select a non-overlapping set, prioritizing peaks with higher scores. This method resulted in 80,019 fixed-length (501bp) peaks.

Re-counting each dataset with CellRanger-ARC revealed that, although the ArchR-derived set increased total peak counts by ∼6%, average counts per cell increased by <2%. Therefore, we proceeded with the pyCistopic-derived peaks, supplementing 1,285 non-overlapping ArchR peaks. The ATAC assay of the Seurat/Signac object on GEO is based on these consensus peak counts.

### Metacells

A chief strength of the multiomic data is the ability to perform direct cross-modality correlations (e.g. between enhancer accessibility and nearby gene expression), but the high sparsity of both modalities often limits the power of these correlations when performed at the single-cell level. Thus “metacell” approaches, i.e. half-measures between single-cells and classical “clusters”, have become popular for analyzing multiome data (79). As the vast majority of our clusters at P24 and later represent biologically equivalent (or extremely similar) groups of cells, these analyses could be performed at the cluster (pseudobulk) level; but it is still often desirable to take advantage of the statistical replicates that are inherently present in single-cell data.

To generate metacells for the P24, P48, and Adult stages, we leveraged biological replicates as determined by DGRP demultiplexing of the multiome dataset. Each stage initially contained cells from 10–11 distinct DGRP lines, but the distribution of cells across these lines was highly uneven. To create more balanced metacells, we manually merged DGRP lines for each stage such that lines with relatively few cells were combined with lines containing more cells, reducing the number of replicate groups to 5–6 per stage (see below). The merged replicates were recorded in a new metadata field, dgrp.new, in the Seurat Multiome object. Some DGRP lines (line_508 and line_307) were dropped (set to a value of 0) as these contain only a few misclassified cells from the demultiplexing outlined above and are not real replicates. The following drops and merges were implemented:

P24:

- Dropped line_307 and line_508

- Merged:

- line_383 and line_897 as lines_383_897
- line_40 and line_441 as lines_40_441
- line_492 and line_406 as lines_492_406
- line_391 and line_505 as lines_391_505
- line_382 and line_320 as lines_382_320

P48:

- Dropped line_508

- Merged:

- line_492 and line_852 as lines_492_852
- line_391 and line_320 as lines_391_320
- line_40 and line_383 as lines_40_383
- line_505 and line_897 as lines_505_897
- line_441 and line_382 as lines_441_382
- line_406 and line_189 as lines_406_189

Adult:

- Dropped line_508

- Merged:

- line_492, line_505 and line_897 as lines_492_505_897
- line_406 and line_852 as lines_406_852
- line_383 and line_441 as lines_383_441
- line_40 and line_320 as lines_40_320
- line_391 and line_382 as lines_391_382

After assigning these merged DGRP replicates, highly heterogeneous clusters (specifically clusters 0, 141, and 144) were excluded from the metacell calculations. For each remaining cluster, metacells were defined as unique combinations of cluster identity, developmental stage, and dgrp.new groupings, resulting in a total of ∼4000 metacells across P24, P48, and Adult stages. Histograms of the number of cells per metacell confirmed that these assignments achieved a more uniform distribution across metacells while retaining biologically meaningful variability (median ∼20 cells per metacell).

Implicitly, the metacells generated by this approach contain many more cells for abundant cell-types as compared to the rarer ones, which in some cases may have metacells based on a few, or even just one cell. However, we argue that the potential caveats of this variance imbalance are offset by the more even representation of different cell types in the metacell space, maximizing biological variation.

To generate metacells at the P0 stage, we used the Python package SEACells(80) v0.3.3 (https://github.com/dpeerlab/SEACells). The first dimension was removed from the LSI reduction embeddings to ensure that it was not retained for SEACells computation, since the first dimension is known to be highly correlated with technical variation as stated previously. The P0 Seurat (.rds) Multiome object was then converted to an Anndata (.h5ad) object by first using the SeuratDisk v0.0.0.9021 function *SaveH5Seurat* (default parameters), then the *Convert* function (dest= “h5ad”). Additionally, the wsnn graph generated for the P0 cells using Seurat’s *FindMultiModalNeighbors* (reduction.list = list(”pca”, “lsi”), dims.list= list(1:80, 2:80), k.nn= 40)) was extracted as a matrix using the *writeMM* function from the Matrix package v1.7-0. This was selected empirically, as other k.nn variables were tested in order to determine the best metacell arrangement.

The Anndata object and the wsnn matrix were imported using Python v3.11.5 and SEACells were calculated by following the tutorial at: https://github.com/dpeerlab/SEACells/blob/main/notebooks/SEACell_computation.ipynb

We used the default parameters for all functions as described in the tutorial, with the following exceptions: The data was preprocessed in Seurat v4.1.4 (as described above) and not in Scanpy; the n_SEACells value was set to 1085 to obtain roughly 50 cells per metacell; the build_kernel_on value was set to “X_lsi” to use the ATAC modality (which is more informative than the RNA modality in our data); and instead of the *model.contruct_kernel_matrix* function, we used the *model.add_precomputed_kernel_matrix* function to utilize our pre-computed wsnn neighbor graph. This method resulted in ∼1000 metacells, with a median of ∼50 cells per metacell. Metacells were validated using the default SEACells quantification metrics (purity, compactness, and separation), as well as manually in our data to ensure as few metacells as possible are composed of cells from multiple clusters.

Ultimately, the two groups of metacells were combined into one metadata field within the multiome object called “Metacells”.

### Calculation of peak-gene linkages

Peak–gene links were inferred separately at P0, P24, P48, and Adult stages, and also all stages together, in R v4.4.1 using Signac’s v1.13.0 *LinkPeaks* function. For each stage, the merged Seurat (.rds) Multiome object was subsetted to remove inconsistent clusters (Adult: 0, 141, 144; P24/P48: 110, 111, 112 (TEs)), and pseudobulk RNA and ATAC profiles were computed per metacell using *AverageExpression* (RNA parameters: assays=”RNA”, return.seurat=”TRUE”, group.by=”Metacells”, ATAC parameters: assays=”ATAC”, slot=”counts”, group.by=”Metacells”). Consensus chromatin assays from the averaged counts were created with *CreateChromatinAssay*, and peak annotations were copied from the original object. Genome coordinates were harmonized with *seqlevelsStyle*, and region-level statistics were calculated using *RegionStats* based on the BSgenome.Dmelanogaster.UCSC.dm6 genome.

Peak–gene links were computed with *LinkPeaks* using a ±50 kb window around each TSS, correlating ATAC counts with normalized RNA expression. For each stage, links were saved as RDS object, exported as a BED file, and converted to tables with absolute z-scores (|z-score|) for ranking.

For P0 progenitor (PG) and neuron subsets, similar pseudobulk averaging, consensus ATAC construction, and peak–gene linking were performed. Genes of interest (e.g., *erm*, *ey*, D) were specifically examined for progenitor–neuron conserved regulatory peaks. High-confidence links (|z-score| > 2) were exported as BED files for downstream motif analyses, including transcription factor enrichment.

### Motif enrichment

To identify cis-regulatory features of neuronal enhancer peaks, we performed motif discovery and enrichment analysis using STREME from the MEME Suite (v5.5.3). Peak regions were converted from BED to FASTA format with bedtools’ v2.30.0 *getfasta* function using the *Drosophila melanogaster* reference genome (BDGP6.46). Each peak set was analyzed independently with STREME, which identifies short, statistically enriched motifs relative to either shuffled input sequences (**Fig. 2e-g**) or user-supplied background sequences (**Fig. 2h-j**).

To assign candidate regulators to the enriched *de novo* motifs, we used Tomtom (MEME Suite v5.5.3) with default parameters to compare all discovered motifs against the CisBP-Drosophila database (2025 release). Tomtom reports significant motif matches based on position weight matrix similarity, with adjusted p-values used to control for multiple hypothesis testing. Final motif sets were summarized to highlight both significantly enriched known motifs and *de novo* motifs with strong similarity to established *Drosophila* transcription factor binding sites (**Table S5).**

### ChromVAR

To estimate transcription-factor (TF) motif activity from chromatin accessibility alone, we applied ChromVAR(58) to the consensus-peak ATAC assay of the multiome object using Signac. The ATAC assay was annotated with motif matches for all Drosophila TF motifs from the CisBP database. For each motif, ChromVAR computes a per-cell bias-corrected accessibility deviation, i.e. the difference between the observed number of fragments in peaks containing that motif and the number expected from the average across all cells, together with a deviation z-score; both are corrected using background peak sets of equal size matched for GC content and mean accessibility, so the resulting motif-activity score reflects the gain or loss of accessibility at motif-containing peaks while controlling for technical biases (e.g., Tn5/GC bias and sequencing depth). Motif activities were computed with RunChromVAR against the *Drosophila* genome (BSgenome.Dmelanogaster.UCSC.dm6, with seqlevelsStyle set to “Ensembl” to match our peak coordinates) using default parameters. TF RNA expression was taken from the log-normalized SCTransform (”SCT”) assay. Excluding stage P0, we then computed, across neurons from the P24, P48, and Adult stages, the Pearson correlation between each motif’s ChromVAR activity and the matching TF’s SCT expression at the metacell level (on values aggregated with Seurat AverageExpression grouped by the “metacells” field). The analysis was restricted to the 72 optic-lobe terminal selector TFs (36) (tsTFs) with an available CisBP motif; for each tsTF, the most strongly correlating motif (positive or negative) across clusters was selected by maximum absolute correlation, and a full tsTF-by-motif correlation matrix was also computed. **Figure S12** shows the metacell-level correlations between tsTF RNA expression (rows) and the activity of these best-correlating motifs (columns).

### SCENIC+

Gene regulatory networks (GRNs) were inferred using SCENIC+ (59) v1.0a2. Multiome data from P24, P48, and Adult stages were combined with previously published scRNA-seq datasets of optic lobes from comparable developmental stages (23, 42). The supplemental RNA-only cells were included solely in the TF–target co-expression step, which does not require cis-regulatory information. Specifically, TF–target gene correlations were computed using GRNBoost2 (arboreto v0.1.6) with default parameters on the combined multiome and RNA-only datasets. All other analysis steps requiring cis-regulatory information were performed exclusively on the multiome cells.

Network inference was performed at the metacluster-level, rather than across all optic lobe cells, to capture context-specific TF–target relationships. To improve interpretability and statistical power, several related metaclusters were merged prior to network construction: M25/M29/M30 (main IPC neurons); M13/M28 (early-born N^ON^ neurons of the main OPC); M27/M35 (N^ON^ neurons of the main OPC from the Ey window); M34/M36 (mid-born N^ON^ neurons of the main OPC); M15/M17 (late-born N^ON^ neurons of the main OPC); M31/M32/M33 (early-born N^OFF^ neurons of the main OPC); M7/M8/M22 (mid-born N^OFF^ neurons of the main OPC); M23/M24/M26 (late-born N^OFF^ neurons of the main OPC); and M19/M20 (mid-born neurons of the vtOPC). Metacluster M1, which contained a heterogeneous mixture of central brain–derived neurons, was excluded. All other metaclusters were analyzed separately.

As our data were already preprocessed in Seurat/Signac, the RNA assays from these objects were extracted and converted to Anndata (.h5ad) objects using SeuratDisk v0.0.0.9021: first with *SaveH5Seurat* (default parameters) and then *Convert* (dest=”h5ad”). Fragments files were extracted from the Multiome Seurat objects using *FilterCells* in R v4.4.1, generating metacluster-specific fragment files for downstream analysis. Preprocessed RNA data from both the Multiome and external RNA-only datasets were integrated and linked to these metacluster-specific fragments via pycisTopic v1.0.2 to create cisTopic objects, which summarize chromatin accessibility patterns across cells and regions. These cisTopic objects were subsequently used to generate multiome inputs for SCENIC+ using *prepare_GEX_ACC*.

Candidate TF–target relationships were then refined using SCENIC+ cisTarget scoring and motif enrichment, requiring overlap between co-expression edges and accessible chromatin regions. To maintain high-resolution regulatory context for each neuronal population, metacluster-specific peak sets (or reduced (merged) peak sets for the combined metaclusters) were used rather than the single consensus peak set. Extended eGRN inference incorporated cistrome information, TF motif rankings, and region–gene linkages, with GSEA-based permutation testing (1,000 permutations), quantile cutoffs (0.85, 0.90, 0.95), and multiple region-to-gene assignment thresholds (top 5, 10, and 15). A correlation threshold (ρ) of 0.05 was applied, except for metacluster M13, where higher thresholds (0.06–0.07) were required to avoid empty outputs. In parallel, direct eGRN inference was performed without motif extension to generate complementary networks.

Motif enrichment was evaluated using the SCENIC+ cisTarget module with the Aerts Lab’s v10 non-redundant public motif collection (v10nr_clust_public) as the motif database. Regulatory regions were scored against metacluster-specific genomic regions extracted from our multiome objects. TF–target edges were retained if they showed significant motif support (adjusted p < 0.05, Benjamini–Hochberg correction) and a minimum co-expression weight of 0.05. Regulon activity was quantified in individual cells using AUCell v1.0a2, with the area under the recovery curve (AUC) computed for each regulon. In total, 28 networks were inferred, of which 23 corresponded to neuronal populations, collectively covering the full diversity of optic lobe cell types. Each network (TF–target edges with various confidence metrics) is provided in **Table S6**.

### SCENIC+ secondary analyses

SCENIC+ regulatory networks were inferred separately for each metacluster (and the merged metaclusters described above) and exported as regulon tables. All downstream analyses for this study were performed in R (v4.3.1), and were performed on neuronal metaclusters (and merged metaclusters) only: M2-M37 **(Fig. 3a**).

To quantify how consistently transcription factors (TFs) regulate target genes across networks (**Fig. 3b**), we first retained only TFs present as regulators in at least 10 networks. For each gene regulated by at least one of these TFs, we first counted the number of networks in which it appeared as a target. For each unique TF–gene pair, we then counted the number of networks in which the TF was inferred to regulate that gene. A regulation ratio was then calculated as: *Networks where TF regulates gene / networks where gene appears*, representing the fraction of networks in which a TF regulates a given gene relative to the total number of networks a given gene is found in. TFs were categorized as Terminal selector, Ecdysone-regulated, or Other, and colored accordingly in the plots. For each TF, the distribution of regulation ratios across its target genes was visualized using boxplots.

To assess how consistently individual genes are associated with the same TF regulator (**Fig. 3c**), the most consistent TF regulator per gene was identified. Specifically, for each gene we selected the TF with the highest regulation ratio (as defined above) was selected. TFs regulating fewer than 10 genes were not considered for this analysis. The distribution of these maximum regulation ratios across genes was visualized using a histogram. Stacked bars represent the number of genes whose most consistent regulator belongs to each TF class (as defined above), and the total height of each bar represents the number of genes whose top regulator falls within that ratio bin.

Next, we aimed to analyze the SCENIC+ results from an enhancer perspective. We first quantified how often an enhancer is reused across networks (**Fig. 3d**). Because each network was generated at the metacluster level and therefore contained metacluster-specific regulatory regions (see “Peak calling” section above), overlapping enhancer coordinates across networks were merged to generate non-overlapping enhancers. Regions were considered part of the same enhancer if their genomic intervals overlapped by a single base pair or more. Each non-overlapping region was then mapped back to the individual regulons they originated from. Enhancer recurrence was quantified as the number of networks in which each non-overlapping enhancer appeared, and the distribution of enhancer reuse across networks was visualized as a histogram.

Finally, we aimed to characterize TF cooperation at the enhancer level (**Fig. 3e**). We considered only enhancer–target gene pairs regulated by two or more TFs within the same network, using the original metacluster-specific enhancer coordinates rather than the merged non-overlapping enhancers. For each enhancer–gene pair with at least two TFs, we determined the composition of TF classes regulating that enhancer. For example, if in one network an enhancer–gene pair is regulated by three TFs (two tsTFs and one ecdysone-regulated TF), the pair contributes one count to the “tsTFs + ecdysone” category. Enhancer–gene pairs were classified into four categories based on the TF classes regulating them: terminal selectors only, ecdysone-regulated only, terminal selectors + ecdysone-regulated, and other combinations. For each network, the proportion of enhancer–gene pairs belonging to each category was calculated. The median proportion across networks was plotted for each TF cooperation category, with interquartile ranges shown as error bars to reflect variability in enhancer counts across networks.

### GO enrichment analysis

To assess the biological processes regulated by each TF, we performed Gene Ontology (GO) enrichment analyses using two complementary frameworks: PANTHER v16 (via the rbioapi R package, v0.8.3) and clusterProfiler v4.12.6. Analysis was performed in R v4.4.1.

First, all unique TF–target gene pairs were compiled from direct eRegulon interactions across all metaclusters from the SCENIC+ results. 103 TFs had at least 50 unique target genes across all networks, for which we independently performed GO enrichment. Gene names were converted to FlyBase IDs, and a non-redundant set of all identified target genes served as the background set.

GO enrichment was performed separately for the two categories: biological process (BP) and molecular function (MF). PANTHER was run using the *rba_panther_enrich* function, with enrichment tested relative to the defined background gene set (organism: *Drosophila melanogaster*, NCBI Taxon ID 7227). Significance was determined using a false discovery rate (FDR) cutoff of 0.05. clusterProfiler enrichment was performed using the *enrichGO* function from the clusterProfiler package (OrgDb: *org.Dm.eg.db*; keyType = “FLYBASE”). Enrichment was assessed relative to the same background universe of target genes, with multiple testing correction performed using the Benjamini–Hochberg procedure. Terms with adjusted p-values ≤ 0.05 were considered significant.

For visualization purposes, only the PANTHER enrichment results were used (**Fig. S14**) to maintain consistency with our previous analyses (23). Fold enrichment values for significant GO terms (FDR ≤ 0.05) were plotted as bar charts, with transcription factors colored according to whether they were annotated as terminal selectors, ecdysone-regulated TFs, or other TFs. Results were saved separately for each ontology, both per TF and per GO term.

### Trajectory analyses at P0

To classify the P0 dataset we transferred labels from the corrected P24 dataset using the multimodal approach we described above. In addition, we performed WNN clustering (resolution=2), obtaining 127 unsupervised clusters (“P0Clusters” metadata field in the Seurat object). This is an intentional under-clustering of the data, since we used these results only for metaclustering (described above) and dividing these data into different lineages which were then analyzed separately.

For the visualization in **Figure 1d** (and **Figs. S7d and S10a-c**), we annotated the progenitor clusters based on the expression patterns of known marker genes (30): Retina (i.e. eye imaginal disc, 2 clusters), Lamina Precursor Cells (LPC, 1 cluster), NE-OPC (1 cluster), NB-OPC (1 cluster), GMC (referring to OPC, 5 clusters), NE-IPC (2 clusters), NB-IPC (2 clusters) and GMC-IPC (2 clusters). 23 P0 clusters represented partially differentiated cells (middle of the trajectories) where the type identities could not be confidently determined. These cells were manually assigned to an identity “0” and grayed out in the UMAP representations. The rest of the P0 cells were labeled and colored based on transferred labels from the P24 dataset. We observed a large number of cells misclassified (by label transfer) as cluster 117 (ENG) or T4/5 neurons. Thus, we also set to “0” the cells classified as cluster 117 but do not group with the genuine ENGs (P0 cluster 48), as well as all cells classified as any of the 8 subtypes of T4/5 but were not within the P0 clusters corresponding to the IPC neurons (below). Other clear cases of misclassification, especially among similar cell-types, could be observed on UMAPs throughout the P0 dataset. Thus, caution should be used when using these cell-type assignments for downstream analyses.

For **Figures 4a-b and S18** (and **Fig. S10d-e**), NB-OPC and GMC clusters (from above) are shared between the N^OFF^ and N^ON^ hemilineages and were included in both UMAPs. We determined that 28 P0 clusters contained N^ON^ progeny of the main OPC (**Fig. 4a**) based on their inclusion of known OPC neurons according to lineage tracing experiments (37) and *ap* expression (except for ENG which seem N^ON^ based on transient Hey and E(spl) expression, **Fig. S10d**, but these glia don’t express *ap*). Similarly, 32 P0 clusters (without *ap* expression) were determined to contain the N^OFF^ progeny of the main OPC (**Fig. 4b**). We maintained the identities transferred from P24 in these UMAPs (as in **Fig. 1d**), but we additionally assigned any clusters containing fewer than 20 cells in each hemilineage to identity “0” and grayed them out. After subsetting, PCA and LSI reductions were re-calculated separately for each hemilineage. We then conducted a grid search involving different combinations of PCA and LSI components to find UMAP representations that best preserve the continuity of neuronal trajectories with the GMC cluster. **Figure 4a** was thus generated using 33 PCA and 29 LSI components and **Figure 4b** was generated using 31 PCA and 38 LSI components.

For **Fig. S16a**, 3 P0 clusters corresponding to the eye imaginal disc were separated. Variable features for both ATAC and SCT modalities were re-determined, followed by re-calculation of PCA and LSI reductions. After WNN clustering of these cells (using 30 PCA and LSI components, resolution=0.8), we annotated them based on previously described markers(81, 82): Photoreceptors R1-6 (3 clusters), R7-8 (2 clusters), cone cells (1 cluster), posterior margin “Cuboidal” cells (1 cluster), “Peripodial” membrane cells (1 cluster) and undifferentiated cells (2 clusters).

For **Fig. S16b**, the LPC and NE-OPC clusters (from above), the 2 P0 clusters corresponding to lamina monopolar cells (L1-5) and the P0 clusters corresponding to Lawf1/2 neurons and epithelial glia (EG), which are produced from common progenitors that derive directly from the OPC NE(83), were separated and plotted together. We did not re-cluster these cells but used the same labels as in **Fig. 1d**, except that any labels found in fewer than 35 cells were manually set to “0” and grayed out. We note that the NE-OPC cluster in this UMAP also produces the NB-OPC (**Fig. 1d**) and all neurons produced from these NB (**Fig. 4a-b**); but it was not plotted again in those UMAPs (below).

For **Fig. S16c**, 16 P0 clusters corresponding to the IPC lineage (determined based on known markers and their continuity on the UMAP) were separated and re-clustered. First, *SCTransform* was re-applied on these cells followed by calculation of PCA and LSI reductions. The UMAP reduction was generated using WNN (20 PCA and 21 LSI components) and clustering was performed using the same neighbor graph with resolution=1.9. Each cluster was then assigned to the identities shown in **Fig. S16c** based on previously published marker genes (84, 85).

In addition, we determined that 9 P0 clusters contained glia that (to our knowledge) do not originate from the OPC, and 27 P0 clusters contained neurons that originate from either the central brain or the tips (Wingless domain) of OPC or IPC. Neurons produced in these regions tend to be very distinct from other regions (24) (e.g. lobula columnar neurons), and at much lower stoichiometry than neurons produced in main compartments of OPC or IPC. These cells are not displayed in any of the UMAPs in **Fig. 4** or **S16**.

### Generation of P0 dendrograms

Using Seurat v4 (R v4.4.1), for each Notch condition, cells corresponding to early neuronal populations were selected using the “P0Clusters” metadata field, excluding the more mature neurons at the tips of the trajectories. Cell barcodes were extracted and used to merge the clusters into their respective “Young Neurons (Non)” and “Young Neurons (Noff)” groups. These are P0 Clusters 41, 31, 32, 12, 62 for NotchOFF and 38, 60, 24, 27 for NotchON. *RunPCA* was rerun on the SCT assay on the entire P0 object with default parameters and npcs=10. SCTransform applies a gene-wise regularized negative binomial regression to remove sequencing-depth–related technical variation and produce variance-stabilized Pearson residuals. Compared to PCA run on log-normalized RNA values, PCA on the SCT assay reduces the influence of library size and other technical artifacts, resulting in principal components that more faithfully capture biological variation among early neuronal populations. PCA was performed on the full P0 object using default parameters with npcs = 10. Similarly, *RunSVD* was rerun on the ATAC assay with default parameters and n = 11. Following this, the NB-OPC, NE-OPC, GMC, Young Neurons (Non), and Young Neurons (Noff) identities were subset from the P0 object, and cluster trees were built with Seurat’s *BuildClusterTree* using the SCT assay / PCA reduction (dims = 1:10) and the ATAC assay / LSI reduction (dims = 2:11), generating the two dendrograms.

### Generation of P0 NB/neuronal coverage plots

In order to visualize the neuronal and NB enhancers of *erm*, *ey*, and *D*, we first subclustered the NBs cluster into distinct temporal windows. Using Seurat’s v4 (R v4.4.1) *FindSubCluster* function (Params: cluster=”NB-OPC”, graph.name=”wsnn”, resolution=0.3, algorithm=1), 3 distinct clusters were identified. These clusters, determined by temporal TF expression, were re-identified as NBs (Early), NBs (Mid), and NBs (Late).

Next, for each TF (*erm*, *ey*, and *D*), we determined a few postmitotic neuronal clusters that did or did not express the TF of interest, making sure to consider whether the cluster came from the same temporal window, as well as Notch status. Once these clusters were determined (Plotted; Dm2 cluster was merged from Dm2a/Dm2b, and Tm9 cluster was merged from Tm9v/Tm9d), Signac’s *CoveragePlot* function was used to plot 10kb US and 10kb DS (except for D, which was plotted with 25kb US and 10kb DS), enhancer regions were manually highlighted.

### Animal Husbandry

All *Drosophila melanogaster* flies were reared at 25°C with males and females used for every experiment except in the analysis of cluster 125 and 166 (split-Gal4 line *Wnt4-p65 ∩ Vsx1-DBD),* and cluster 76 *(split-Gal4 line* Mirr-VP16 *∩ Hmx-DBD),* where only females were used. Pupal dissections were staged by collection of P0 white pupa and waiting for 48 hours (for P48) with vials kept at 25°C. Adult dissections were performed within 48 hours of pupal eclosion. Sample sizes for each experiment are listed within figure legends. Animals were always selected randomly after confirmation of genotype.

Sparse neuron labeling for cell type annotations were performed by crossing *hsFLP2:PEST;UAS-FSF-CD4tdGFP;MKRS/TM6B,Tb,Hu* female virgins to *;;M24-Gal4* (cluster 163/164: Mi21a/b, Dm2a/b),;*AstC-Gal4/Cyo-G;* (cluster 178: MeLo3b), *Vsx1-DBD*; *wnt4-p65*/*CyO* (cluster 166, Dm3c)), CG44325-DBD; *Dip λ -VP16/cyo* (clusters 75 and 79: Y12 and Tlp12), *Mirr-VP16, Hmx-DBD / Tm6c* (cluster 76: Y13), or;;CCHa2/TM3,Sb (clusters 44 and 93: LC20b and MeVP1) males with non-Tb P0 heat shocked for 5-10 minutes at 38°C. To annotate cluster 212, *;;UAS-CD4:tdGFP* female virgins were crossed to *w;Dll-VP16,ex1/CyO-G;Lmpt-DBD#M3/TM6C,Sb,Tb* males. To annotate cluster 203, w;*UAS-CD4:tdTomato;GMR-Gal80* female virgins were crossed to *w;;CCHa1-Gal4/TM6C,Sb,Tb* males, and for sparse labeling *hsFLP2:PEST;;CCHa1-Gal4/TM6B,Tu,Hu* female virgins were crossed to *UAS-FSF-CD4tdTomato;GMR-Gal80* males. Origins of all stocks are reported in the Key Resources Table (below).

### Generation of split-Gal4 lines

#### Dll-VP16

The Dll-VP16 line was generated using CRISPR-mediated T2A split-GAL4 knock-in by WellGenetics Inc. (Taipei, Taiwan). Briefly, the gRNA sequence CACTAGTGATGGATTCGTCT[CGG] was cloned into a U6 promoter-driven plasmid. The donor construct consisted of a T2A-VP16AD-GMR-RFP cassette containing a T2A peptide, VP16 activation domain fused to Zip (VP16AD-Zip), an SV40 3′ UTR, and a floxed GMR-RFP selection marker. This cassette was flanked by two homology arms and cloned into pUC57-Kan for homology-directed repair. The gRNA plasmid(s), hs-Cas9 plasmid, and donor plasmid were co-injected into embryos of the control strain w^1118^. CRISPR-mediated double-strand breaks at the Dll locus were repaired via homologous recombination, resulting in insertion of the T2A-VP16AD-GMR-RFP cassette. F1 progeny carrying the GMR-RFP selection marker were identified and validated by genomic PCR and sequencing.

#### Lmpt-DBD

The Lmpt-DBD line was generated using MiMIC-based recombination-mediated cassette exchange (RMCE). Donor plasmid pBS-KS-attB2-SA(0)-T2A-GAL4DBD-Hsp70 (Addgene #62902) was injected into embryos derived from crosses between the coding intronic MiMIC line Lmpt[MI05730] (BDSC #44862) and a ΦC31 integrase source line.

DIP-lambda-VP16: The DIP-lambda (CG45781)-VP16 line was generated by injecting the donor plasmid pBS-KS-attB2-SA(0)-T2A-dVP16AD-Hsp70 (Addgene #62905) into embryos derived from crosses between the coding CRIMIC line DIP-lambda [CR70096] (BDSC #97155) and a ΦC31 integrase source line.

Embryo injections and recovery of transformants were performed by BestGene (Chino Hills, CA). Progeny lacking the yellow marker (y⁻), indicative of successful cassette exchange, were screened, and correct integration orientation was confirmed by PCR genotyping.

### Antibody generation

A polyclonal antibody against Pdm3 was generated by Genscript (https://genscript.com). Amino acids 801-1292 of Pdm3 was used to immunize guinea pigs: MSTSAVSSTLPQISLRHPDELTAPQMDLKPLELSASTSPPAPPPRHHFGHSLRGSSTVSPKHSPQGRMGGSGGSTTTGMNLSQHHERHDRLERLERQERHERRSHTPTATATRASVSSSSSAGHHGGSLPSGRLSPPSSAPSNSAANSISDRGYTSPLFRTHSPQGHALSLGGSPRLERDYLGNGPSSGTATSTSSCGAPTAAGSSATANVLSSINRLNASNGELTITKSLGAPTATATRASSASPRDDSPGPGPSTSSVSHMQPLKLSPSSRSEPPHLSPNGNDNDNDLLMDSPNEPTINQATTNVVDGIDLDEIKEFAKAFKLRRLSLGLTQTQVGQALSVTEGPAYSQSAICSSALAAQMYAAQLSTQQQNMFEKLDITPKSAQKIKPVLERWMKEAEESHWNRYKSGQNHLTDYIGVEPSKKRKRRTSFTPQALELLNAHFERNTHPSGTEITGLAHQLGY EREVIRIWFCNKRQALKNTVRMMSKGMV.

### Immunohistochemistry

Animals were dissected in cold Schneider’s Insect Medium. Pupal and adult brains were fixed in 4% PFA at room temperature for 25-30 minutes. L3 brains were fixed in 4% PFA at room temperature for 20 minutes. All wash volumes are 500 µL. Brains were briefly washed with 0.3% PBS-Triton (PBST) three times before nutating for 15 minutes. After washing, samples were blocked in 5% donkey serum with PBST for one hour at room temperature before incubating with primary antibodies at 4°C overnight. Uniquely, the experiments using Bifid antibody were fixed for double the amount of time on ice. After the removal of the primary antibody solution, brains were washed three times for 10-15 minutes in PBST before incubation with secondary antibodies at 4°C overnight. The samples were then washed for 10 and 15 minutes in PBST and finally, for 15-20 minutes in PBS before mounting in Slowfade Gold with the desired orientation. All samples were imaged with a Leica Stellaris 8 confocal microscope using a 63x glycerol objective (NA=1.3). All antibodies used, their concentrations, and their origin can be found within the Key Resources table (below).

### HCR *in situ* hybridization

HCR-3.0-style probe pairs for *Glob1* (NM_169679.3, Amplifier: B5, 11 probe pairs) and *Bma* (NM_001382075.1, Amplifier: B4, 33 probe pairs) were designed (86) for HCR in situ hybridization against the CDS of the first splicing variant for *Bma* and against the full cDNA for *Glob1* due to its smaller length. *Vsx1-RB* (NM_001297969.1, Amplifier: B3, 40 probe pairs) probe pairs were designed by Molecular Instruments. All HCR specific wash buffers and hairpin amplifiers were obtained from Molecular Instruments.

For HCR *in situ* hybridization, previously established protocols (87) were followed with adjustments described below. Brains were dissected within cold Schneider’s Insect Medium and fixed in 500 µL of 4% PFA with 0.03% PBS-Triton (PBST) while nutating for 30 minutes at room temperature. Brains were quickly washed three times with 1% PBST and then 3×15 minutes. Brains were then prehybridized in 100 µL of probe hybridization buffer at 37°C while nutating for 30 minutes before the probe solution was added. Samples were incubated in probe solution for 24-48 hours at 37°C. After incubation, brains were washed 4×10 minutes with 100 µL of probe wash buffer at 37°C before they were washed 2×5 minutes with 1000 µL of 5X SSCT at room temperature. They were then equilibrated with 100 µL of amplification buffer at room temperature. During this equilibration, 2 or 4 µL of each 3 µM hairpin stock was aliquoted into individual PCR tubes and incubated in thermocycler for 90 seconds before incubation at room temperature in the dark for at least 30 minutes. Then, hairpins were spun down and a 100 µL mixture with amplification buffer was used to resuspend them before incubation with the brains at room temperature in the dark for 24 hours. Brains were washed 2×5 minutes with 5X SSCT, 2×15 minutes with 5X SSCT, quickly rinsed with PBS-1X twice before mounting with Slowfade Gold and imaged. Unless noted otherwise, all washes were 150 µL and samples were nutating.

The sources and concentrations used for all HCR probes and amplifiers can be found within the Key Resources table (below).

### Image Analysis and Presentation

Images were created and analyzed within Imaris 9.6.1 and adjusted for figures within Adobe Illustrator 2025 and Adobe Photoshop 2025. They are shown as either maximum intensity projections or single slices to show protein expression within cells, or with three-dimensional (3D) reconstruction using Blend mode with glow color settings to visualize more detailed neuronal morphology as indicated within figure captions.

## Key Resources Table

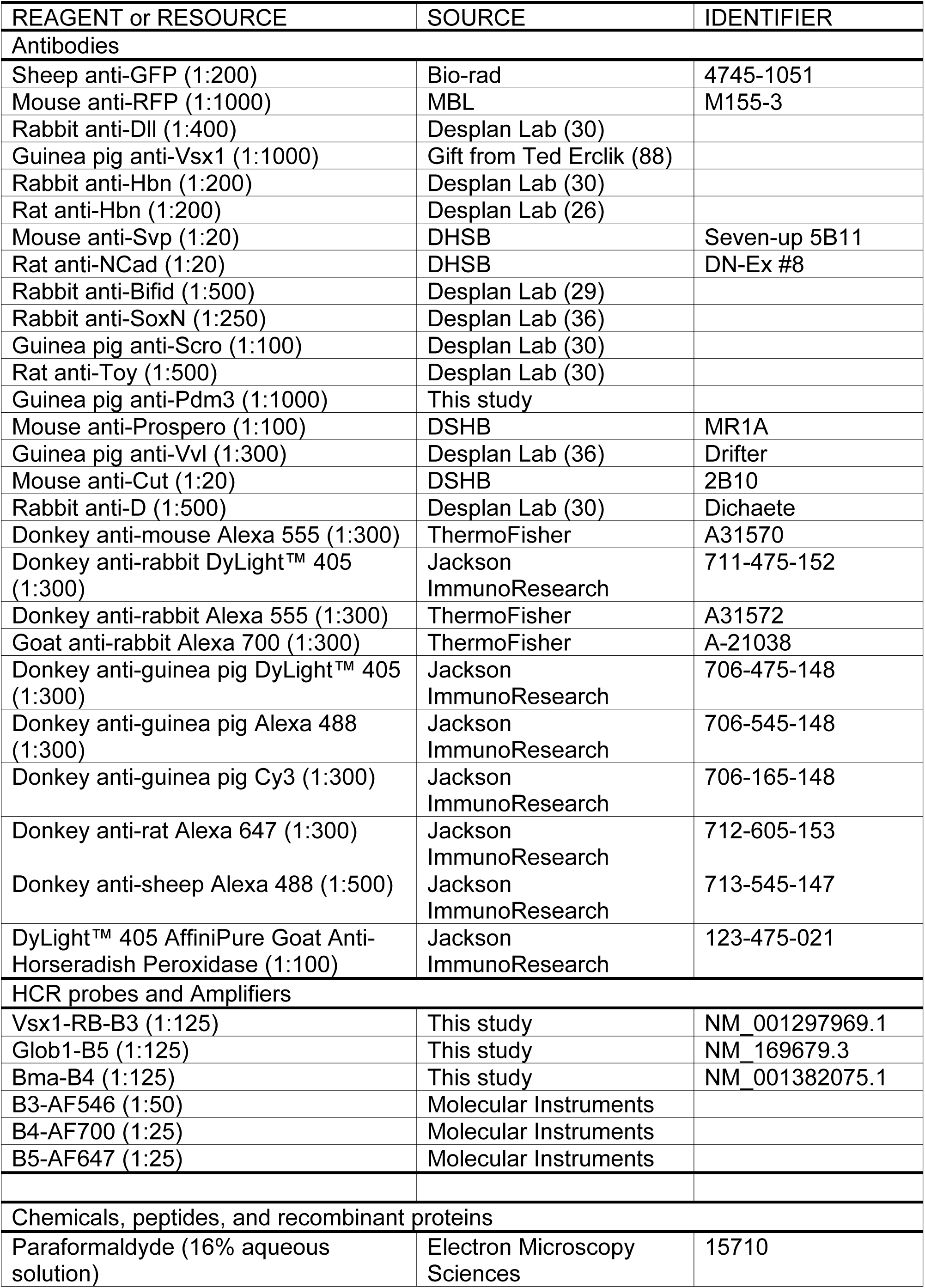
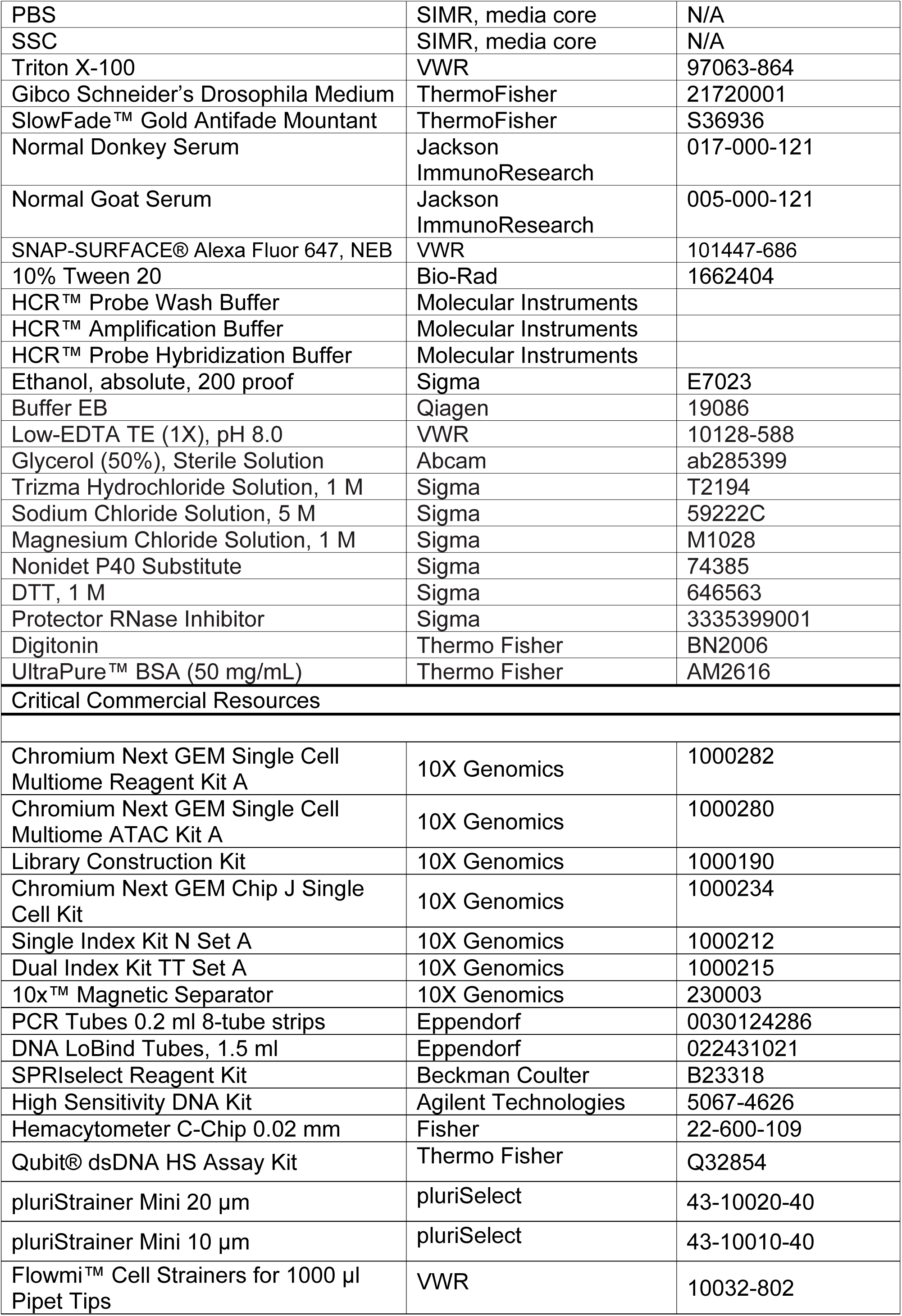
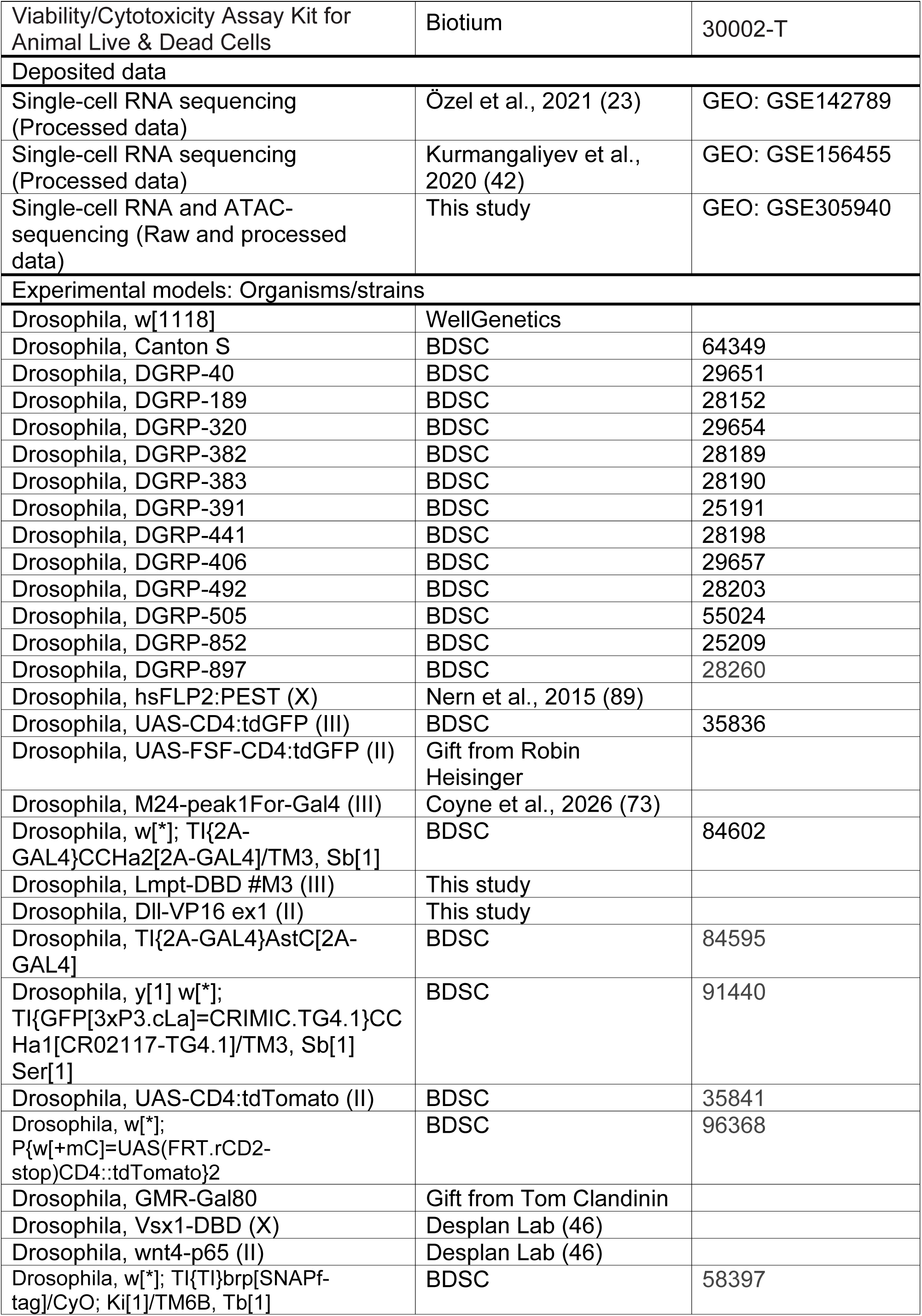
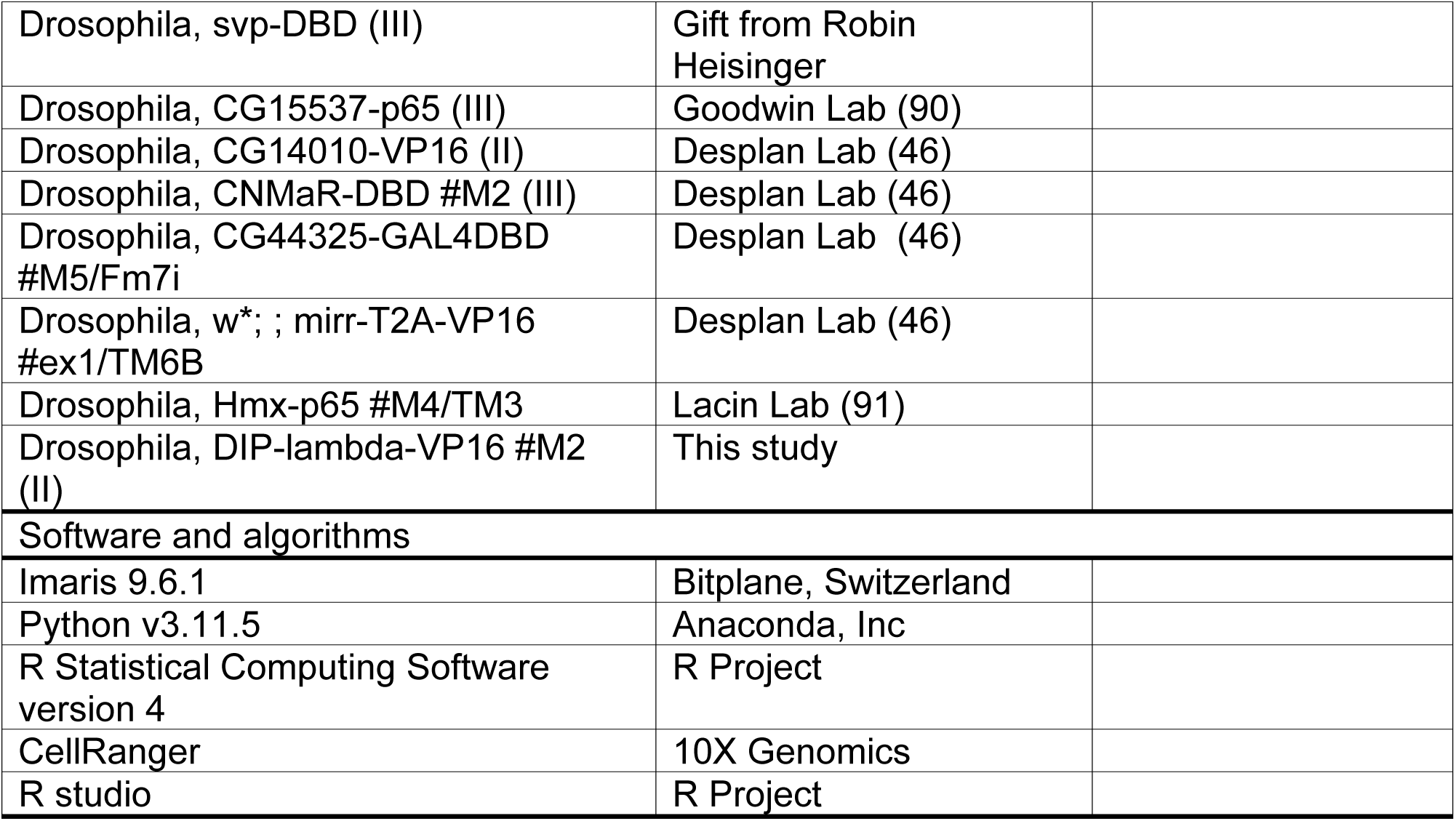

---

**Figure S1:**
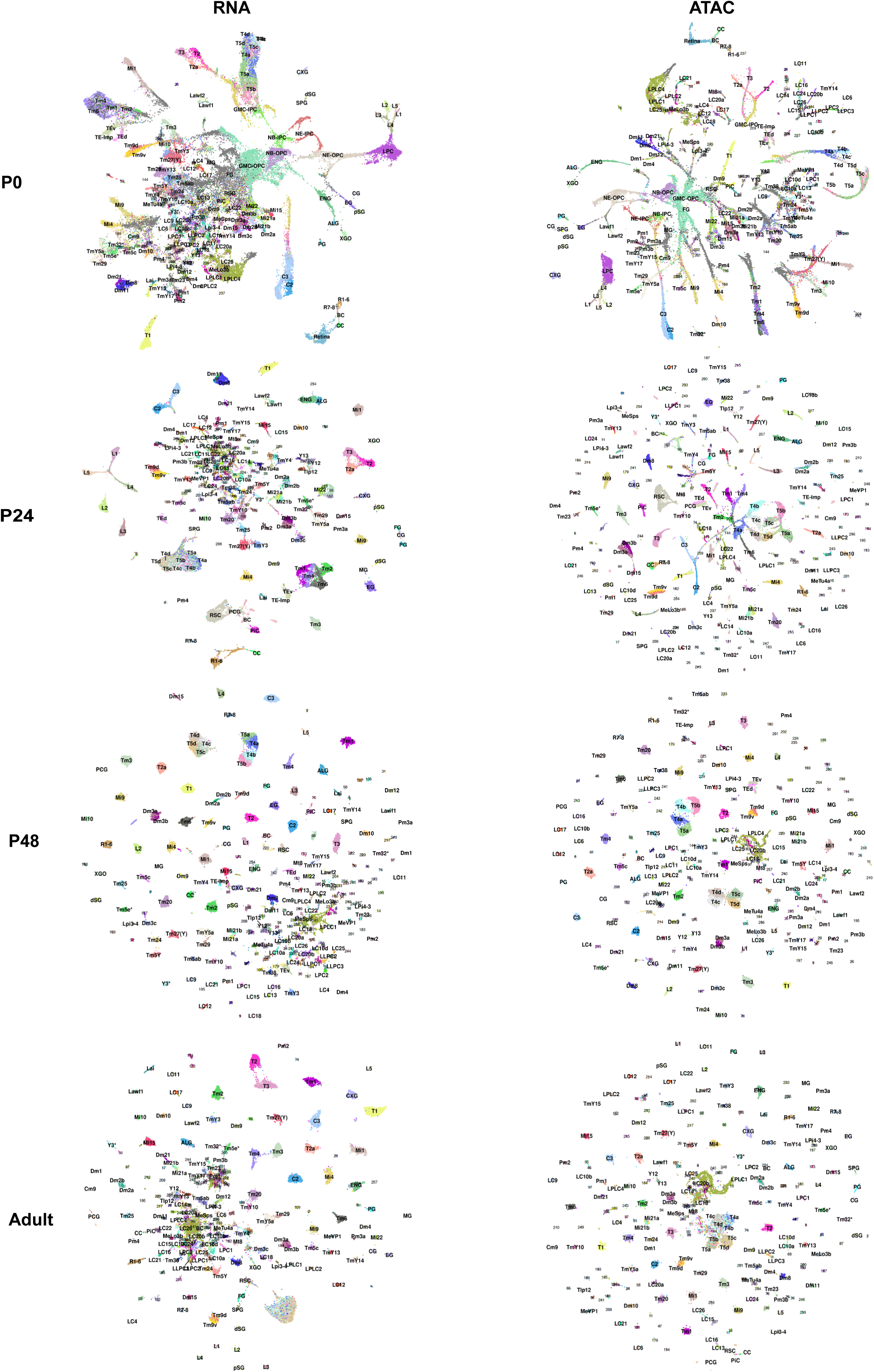
RNA and ATAC UMAP reductions by developmental stage. **a**, For each developmental stage (P0, P24, P48, Adult), UMAP embeddings of all cells are plotted using either RNA or ATAC modalities. Cells are colored by cluster identity (matching the WNN UMAPs in Fig. 1); labeled clusters denote assigned cell-type identity, and unlabeled numeric clusters remain unannotated. P24-Adult reductions were calculated on 150 PCA (RNA) or LSI (ATAC) components, and P0 reductions were calculated on 70 PCA or LSI components.

**Figure S2:**
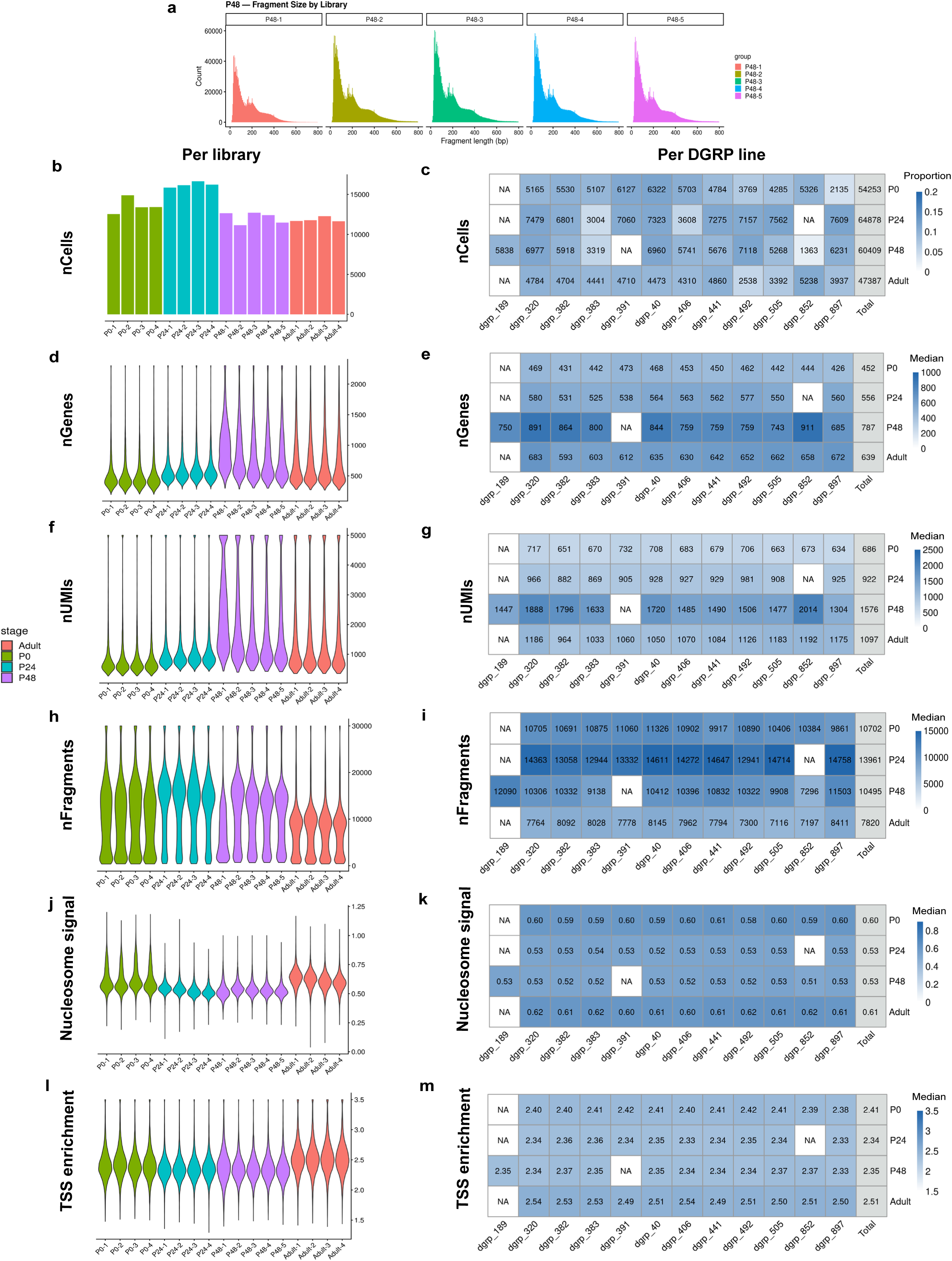
Quality control metrics by library and DGRP strain across development. **a**, ATAC fragment length distributions for P48 nuclei, split by library of origin (P48-1 – P48-5). **b**, Number of cells per library across all four stages, colored by stage (P0, green; P24, teal; P48, purple; Adult, red). **c**, Heatmap of the proportion of each stage’s cells contributed by each DGRP line. Cell values give the corresponding true cell counts, and the “Total” column reports the aggregate cell number per stage. **d**, Violin plots of gene number per library, colored by stage. **e**, Heatmap of median gene number per DGRP line and stage. **f**, Violin plots of UMI count per library, colored by stage. **g**, Heatmap of median UMI count per DGRP line and stage. **h**, Violin plots of fragment count per library, colored by stage. **i**, Heatmap of median fragment count per DGRP line and stage. **j**, Violin plots of nucleosome signal per library, colored by stage. **k**, Heatmap of median nucleosome signal per DGRP line and stage**. l**, Violin plots of TSS enrichment per library, colored by stage. **m**, Heatmap of median TSS enrichment per DGRP line and stage. In **c, e, g, i, k,** and **m**, NA denotes a DGRP line that was not used for the experiments of the corresponding stage. In **e, g, i, k,** and **m**, the “Total” columns show the stage-wide median.

**Figure S3:**
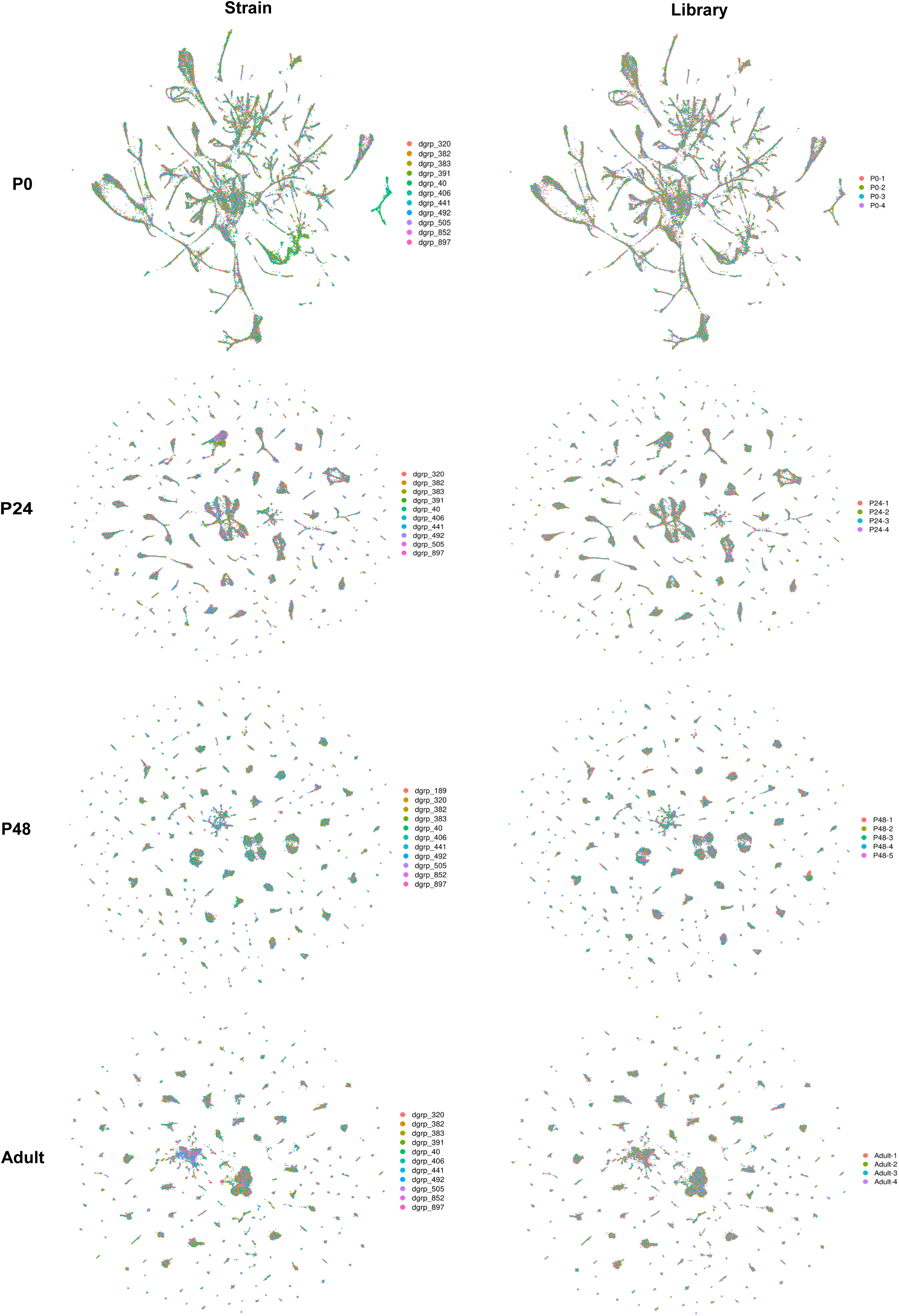
WNN UMAPs by library and DGRP strain across development. **a**, WNN UMAP embeddings at each developmental stage (same as in Fig. 1), with the cells colored by DGRP strain (left) or library (right) of origin.

**Figure S4:**
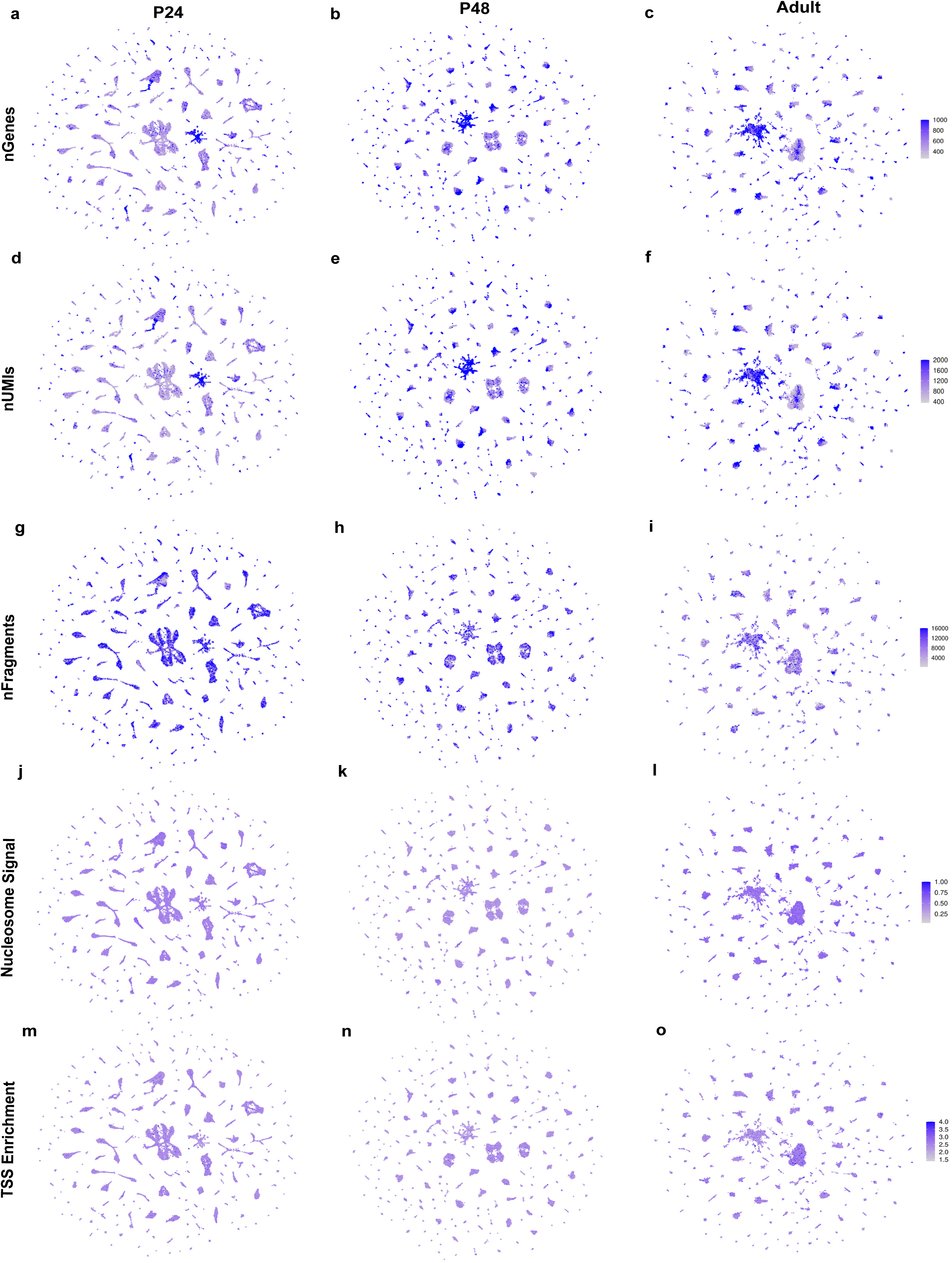
Quality control metrics overlaid on WNN UMAPs at P24, P48, and Adult. **a-o**, Number of genes (**a**-**c**), UMIs (**d**-**f**), fragments (**g**-**i**), nucleosome signal (**j**-**l**), and TSS enrichment (**m**-**o**) values, overlaid on WNN UMAPs of the P24 (**a**, **d**, **g**, **j**, **m**), P48 (**b**, **e**, **h**, **k**, **n**) and Adult (**c**, **f**, **i**, **l**, **o**) stages (same as in Fig. 1). Color scales are capped when necessary to preserve dynamic range of values.

**Figure S5:**
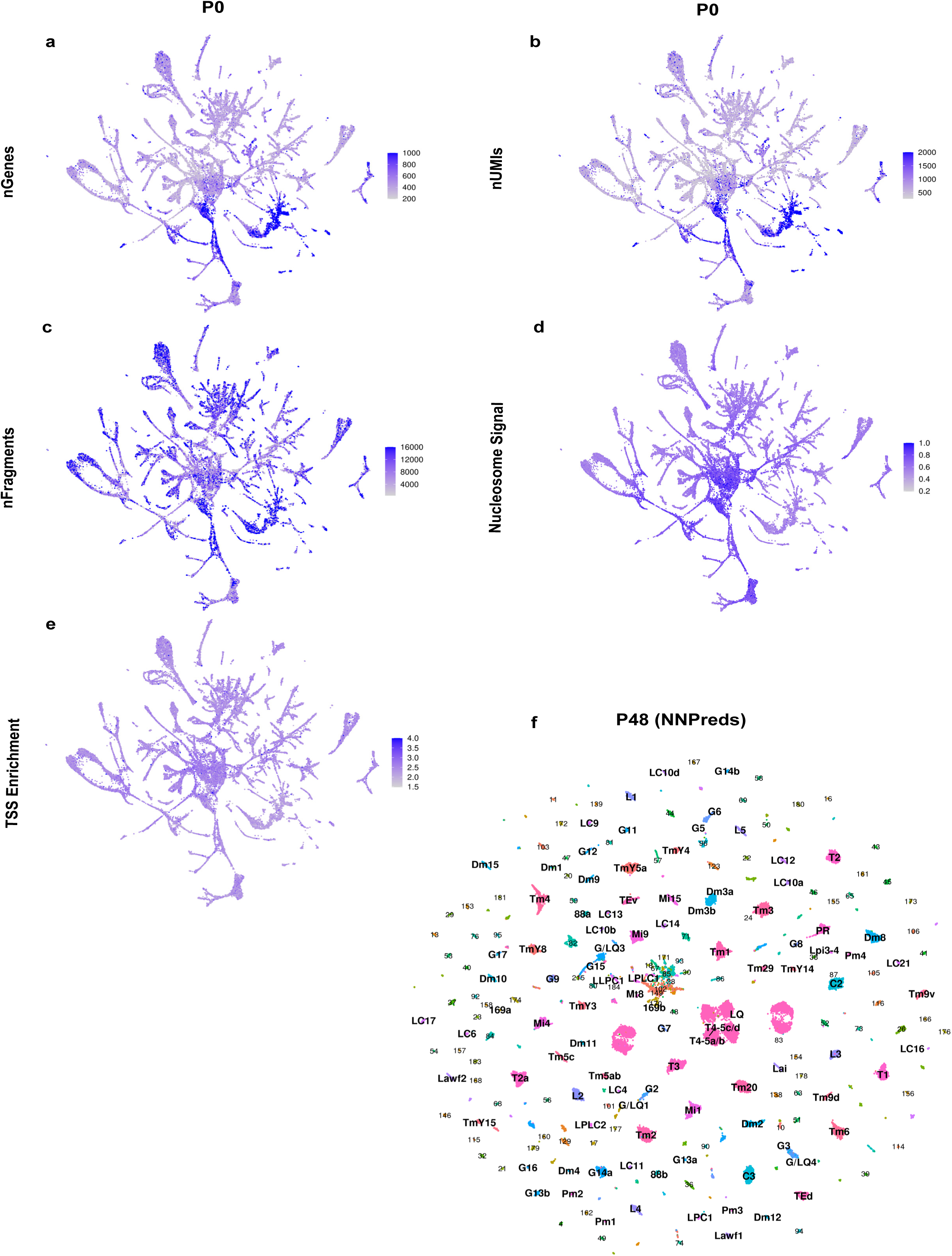
Quality control metrics overlaid on WNN UMAP at P0, and P48 WNN labeled by supervised neural network classifications. **a-e**, Number of genes (**a**), UMIs (**b**), fragments (**c**), nucleosome signal (**d**), and TSS enrichment (**e**) values, overlaid on the WNN UMAP of the P0 stage (same as in Fig. 1d). Color scales are capped when necessary to preserve dynamic range of values. **f**, WNN UMAP of the P48 stage (same as in Fig. 1b), colored and labeled by cluster identity assigned using a neural network classifier trained on the marker gene expression profiles of 206 clusters previously resolved with scRNA-seq (23).

**Figure S6:**
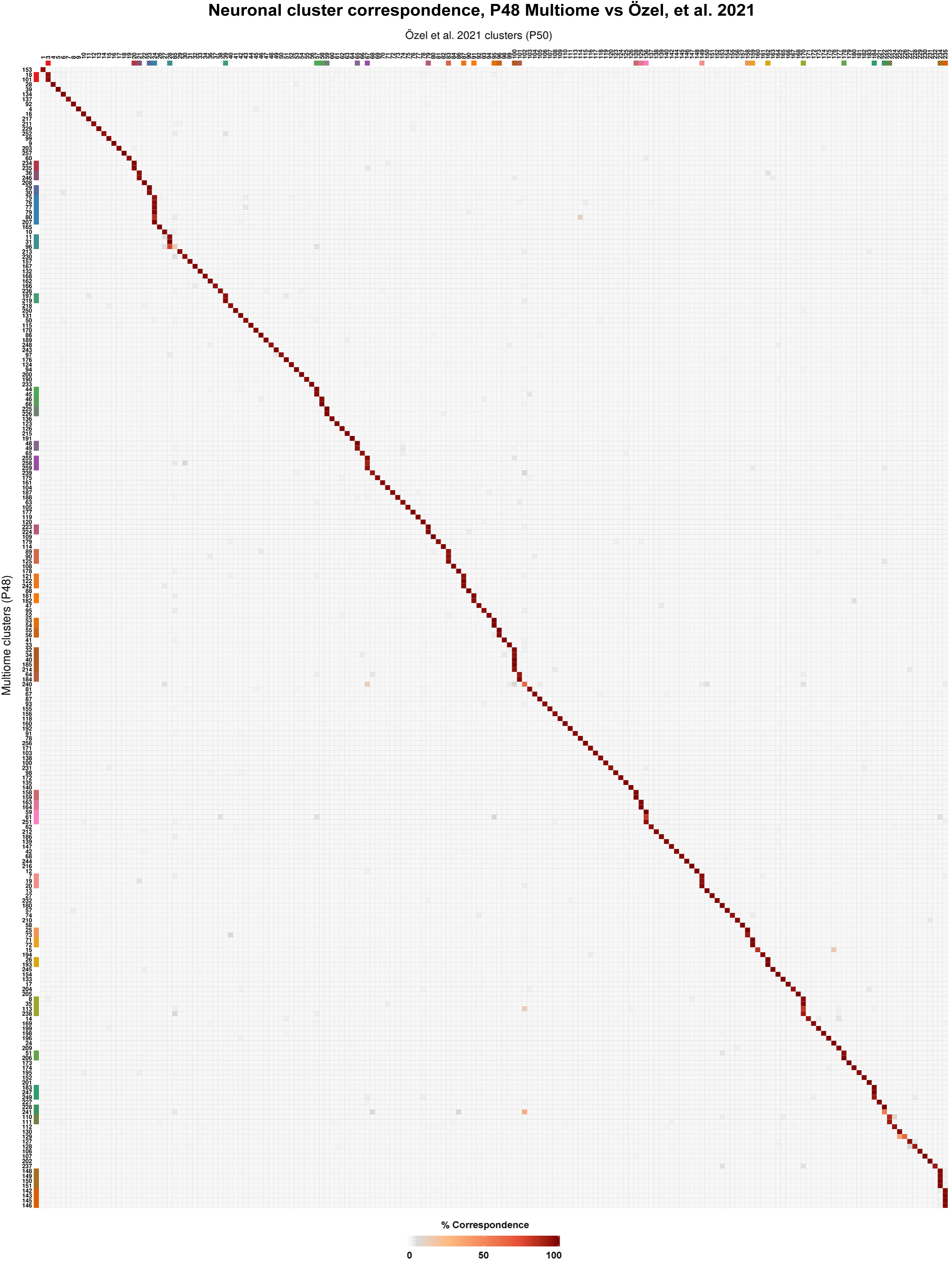
Cluster correspondence between the P48 multiome and the P50 scRNA-seq datasets. **a**, Heatmap depicting the percentage of cells in each of the P48 multiome neuronal clusters (rows) to each of the P50 scRNA-seq atlas (23) neuronal cluster annotations (columns), assigned by label transfer on the RNA modality only. Rows are ordered by their best-matching cluster, which are highlighted along the diagonal. Colored annotation bars mark the previous atlas’ clusters that have been split into two or more clusters in the multiome.

**Figure S7:**
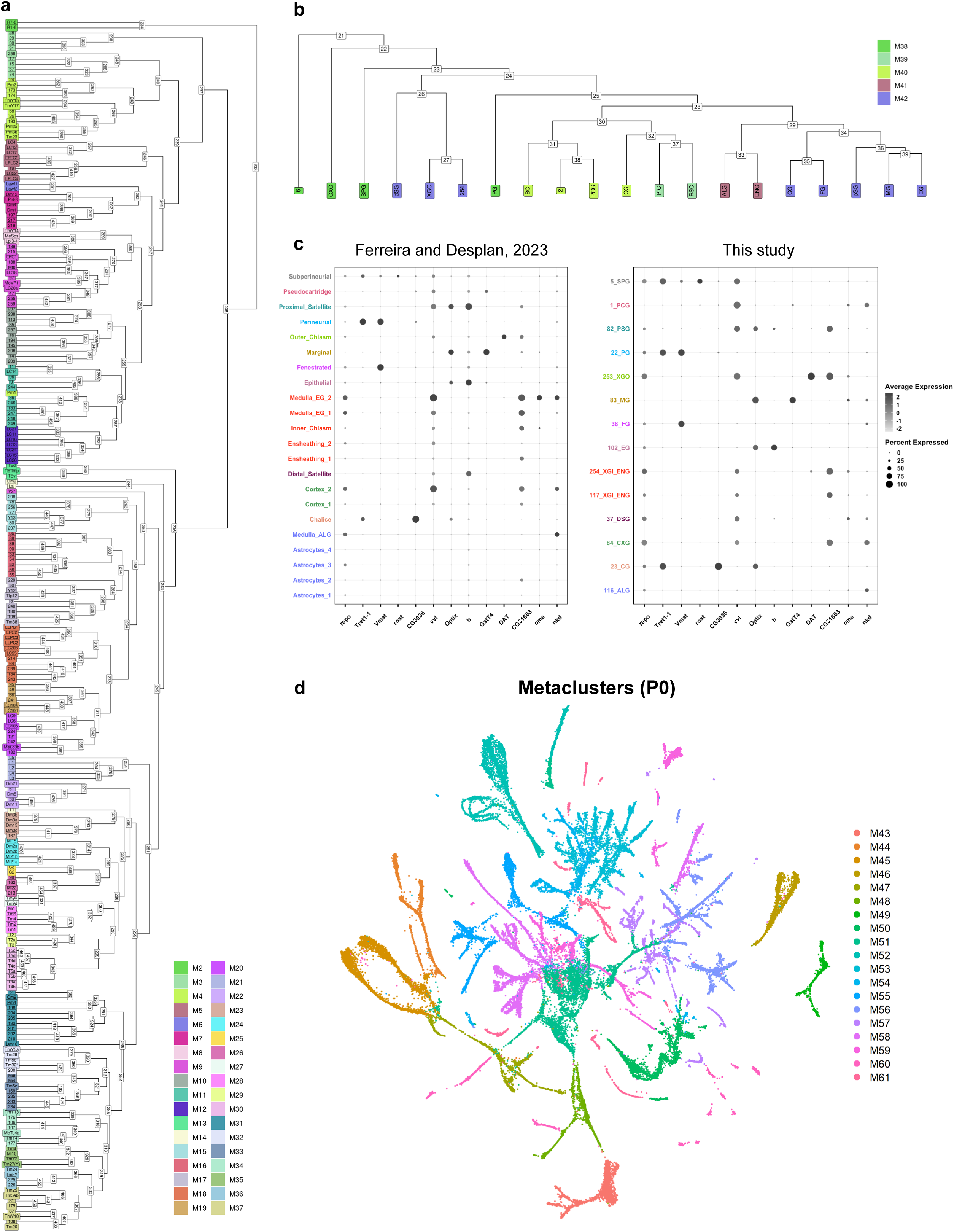
Meta-clustering of the multiome data and the annotation of glial clusters. **a-b**, Hierarchical tree of neuronal clusters (**a**) and non-neuronal clusters (**b**) based on transcriptomic similarity using 100 principal components (**a**) or 20 principal components (**b**) of the RNA modality at P24. The dendrograms were generated using Seurat’s *BuildClusterTree* function. Branch lengths indicate transcriptional distance between clusters. See SI Methods for more details. **c**, Annotations of glial clusters identified in this atlas (right), based on previously described marker genes (44) (left). **d**, UMAP reduction of the P0 dataset (Fig. 1d), colored by metacluster assignment.

**Figure S8:**
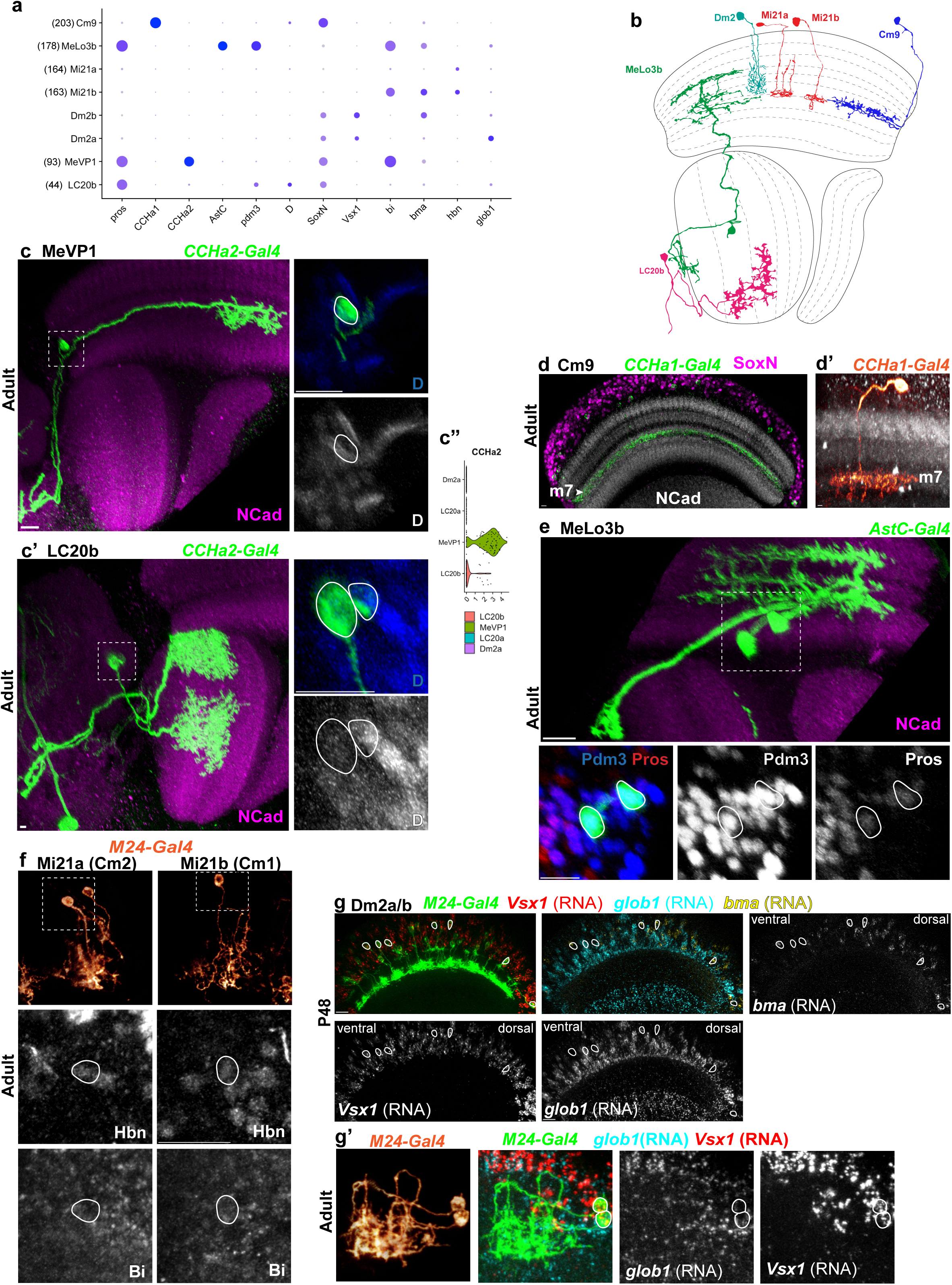
Cell type annotations with single-gene and enhancer-GAL4 lines. **a**, mRNA expression of the relevant marker genes for the indicated clusters in the adult stage. Dot size indicates the fraction of cells expressing the gene and color intensity corresponds to log normalized and scaled expression level. **b**, Cross section of the optic lobe neuropils showing cartoon representations of the newly annotated optic lobe neurons (except MeVP1). **c**, *CCHa2-Gal4* driving CD4:tdGFP (FLPout). This single gene *Gal4* line labels both clusters 93 and 44 at adult. These clusters are distinguished by the expression of D in cluster 44. The medullar arborization and axon path to the central brain matches the connectome (17) cell type MeVP1. In both panels, Anti-GFP (green) and anti-NCad (magenta) are shown as maximum intensity projections in the largest image to the right. The region within the white box is shown as a maximum intensity projection at a higher magnification in the two panels to the left. In the top right multi-channel panel, the neurons are marked with anti-GFP (green). In the bottom right panel, cell bodies outlined in white do not express D. The absence of D indicates that these cells are part of cluster 93. Therefore, we annotated cluster 93 as MeVP1. n =4 brains. **c’**, *CCHa2-Gal4* driving CD4:tdGFP (FLPout). The morphology of the cells that express D matches the connectome (17) cell type LC20b. Of note, LC20a shares a similar dendritic morphology with LC20b but the two cell types are transcriptionally quite distinct, and the cell bodies of LC20a are located dorsally, whereas the cell bodies of LC20b are located ventrally. In the bottom right panel, cell bodies outlined in white express D. Therefore, we annotated cluster 44 as LC20b. n = 4 brains. **c’’**, Violin plot of *CCHa2* expression showing that LC20b neurons express it at much lower levels than MeVP1, but significantly higher than the background level in all other clusters. **d**, *CCHa1-Gal4* in the presence of *GMR-Gal80* driving *CD4:tdTomato*. Anti-RFP (green), anti-SoxN (magenta), and anti-NCad (white) are shown as maximum intensity projections with medulla layer 7 (M7) annotated with white arrow. CCHa1, a neuropeptide, was specially expressed in cluster 203 (and photoreceptors) and labeled medulla neurons that innervate layer M7. n = 5 brains. (**d’**) *CCHa1-Gal4* in the presence of *GMR-Gal80* driving *CD4:tdTomato* (FLP out). Cluster 203 was identified to be Cm9 by sparse labeling with anti-RFP (glow) and anti-NCad (white). n = 5 brains. **e**, *AstC-Gal4* driving *CD4:tdGFP* (FLP out). The morphology and dorsal positioning of the cell bodies match the connectome (17) cell type MeLo3b as shown in **b**. Anti-GFP (green) and anti-NCad (magenta) are shown as maximum intensity projections. The region within the white box is shown as a maximum intensity projection at a higher magnification in the other three panels. The bottom left multi-channel panel shows the cell bodies of these neurons marked with anti-GFP (green), anti-Pdm3 (blue), and anti-Pros (red). The cell bodies outlined in white coexpress Pros and Pdm3, which can be seen on the bottom right panels. n = 4 brains. **f**, *M24-Gal4* (73) driving *CD4:tdGFP* (FLP out) showing sparsely labeled neurons that co-express Dll and Hbn. Based on their morphology, we named these cell-types Mi21a and b that are distinguished by Bifid (Bi) expression. Bifid is a target of the BMP pathway, which implies that the different fates of these subtypes is established by Dpp signaling (29), similar to Dm3a/b (23). Anti-GFP (glow) is shown as a 3D reconstruction with cell bodies shown with increased magnification below with anti-Hbn (magenta) and anti-Bi (white). n = 15/9 optic lobes/brains. **g**, HCR *in situ* hybridization for indicated mRNAs was used to identify Dm2 subtypes at P48. *M24-Gal4* drives *CD4:tdGFP* (FLP out) with *Vsx1* (red) and endogenous GFP (green) marking Dm2s at P48. Visualized as maximum intensity projections, cells that express Vsx1 were circled in white with *glob1* (blue) and *bma* (yellow) shown on dorsoventral axis. The same sample is shown to the right with single channels images for both *glob1* and *bma* with the ventral medulla on the left and dorsal on the right. n = 5/3, optic lobes/brains. **g’**, Sparsely labeled Dm2 subtypes in adult brains were visualized using *M24-Gal4* driving *CD4:tdGFP* (FLP out) with endogenous GFP signal (green). HCR *in situ* hybridization against *Vsx1* (red) and *glob1* (blue) is shown as maximum intensity projections of a representative image. Two Dm2 neurons with differential *glob1* expression are shown as 3D reconstructions of the endogenous GFP signal (top left). Single channel images for *glob1* and *Vsx1* are shown below with cell bodies circled in white. n = 4/3, optic lobes/brains. Scale bars: 10 μm. Developmental stage is indicated to the immediate left of the panels.

**Figure S9:**
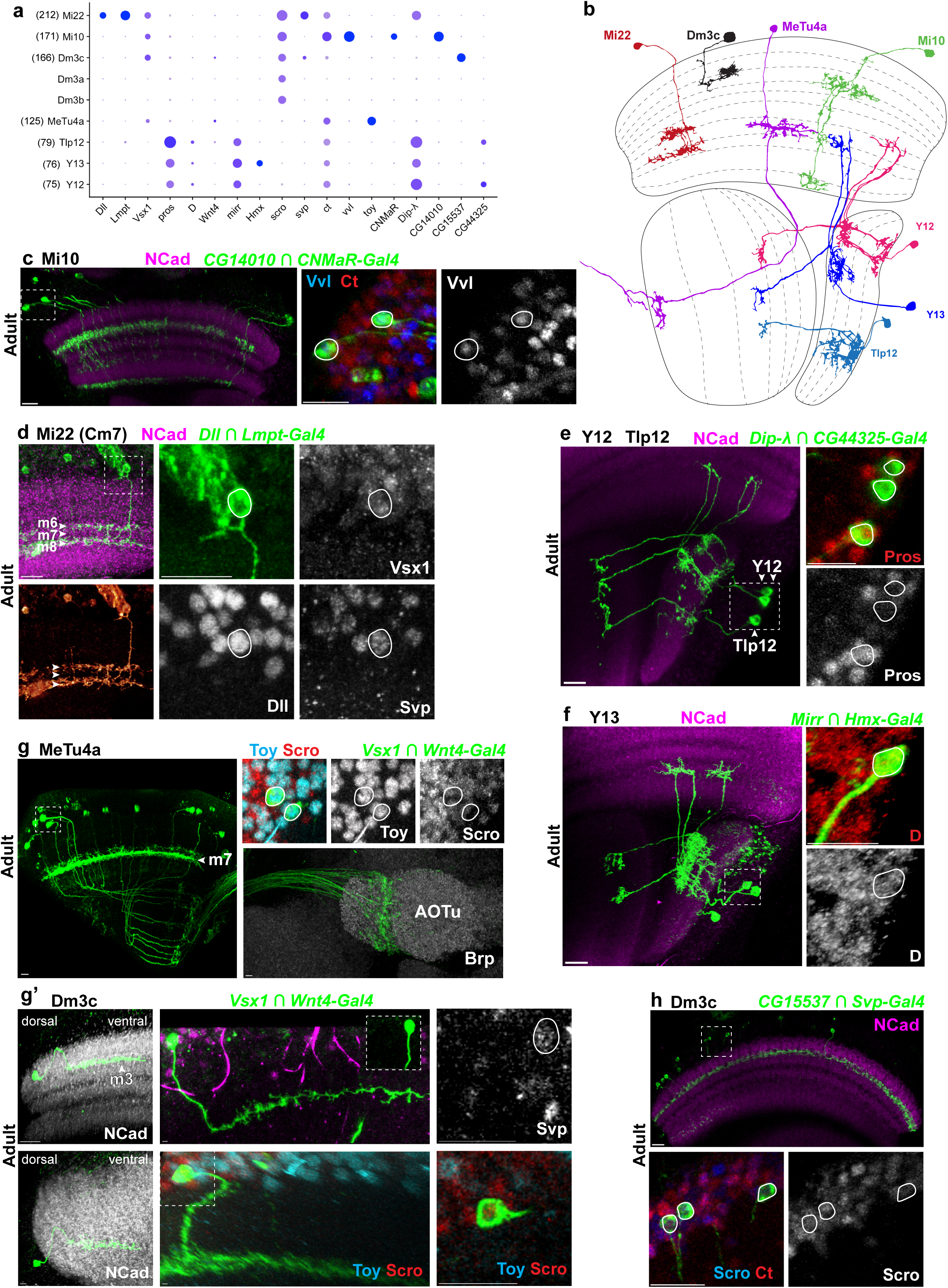
Cell type annotations with split-Gal4 lines. **a**, mRNA expression of the relevant marker genes for the indicated clusters in the adult stage. Dot size indicates the fraction of cells expressing the gene and color intensity corresponds to log normalized and scaled expression level. **b**, Cross section of the optic lobe neuropils showing cartoon representations of annotated optic lobe neurons. **c**, *CG14010-VP16 ∩ CNMaR-DBD* driving *CD4:tdGFP*. Anti-GFP (green), anti-NCad (magenta), anti-Vvl (white), and anti-Cut (white) are shown as maximum intensity projections. The neuron to the far right shows the full axonal tract, and the morphology matches FlyWire (16) Mi10 as shown in **b**. The region within the dashed white box is shown as maximum intensity projections at a higher magnification in the right panels. The middle multi-channel panel shows the cell bodies of these neurons marked with anti-GFP (green), anti-Vvl (blue), and anti-Ct(red). The expression of Vvl is shown in white on the right panel to validate cluster 171 identity. n = 4 brains. **d**, *Dll-VP16 ∩ Lmpt-DBD* driving *CD4:tdGFP*. Anti-GFP (green) and anti-NCad (magenta) are shown as maximum intensity projections with medulla layers 6-8 (m6, m7, m8) annotated with sequential white arrows. In the lower left panel, anti-GFP (glow) is shown as a 3D reconstruction of the same neuron. The region within the dashed white box with the cell body circled in white is shown as maximum intensity projections at a higher magnification (right) with separate panels for anti-GFP, anti-Dll, anti-Vsx1, and anti-Svp whose co-expression validates cluster 212 identity. Based on their morphology, we named this cell type Mi22 which matches connectome (17) cell type Cm7 as shown in **b**. n = 10 brains. **e**, *Dip λ -VP16 ∩ CG44325-DBD* driving *CD4:tdGFP* (FLP out). Anti-GFP (green) and anti-NCad (magenta) are shown as maximum intensity projections. This split-Gal4 targets both clusters 75 and 79. These clusters are distinguishable by the levels of Pros expression present. The panels to the right show Pros expression with anti-GFP (green) and anti-Pros (red). The top two cells express very little Pros, indicating that they are part of cluster 75. The bottom cell expresses a much higher level of Pros, indicating that it is part of cluster 79. In the panel to the left, there are two cells in cluster 75 (Y12), and one cell in cluster 79 (Tlp12). The morphology of Tlp12 and Y12 both match the morphology found in the connectome (17) for their respective cell types. Of note, Tlp12 does not arborize in the medulla, whereas Y12 can have up to 3 arborizations in the medulla. These cell types also differ in their lobular plate innervation. n = 5/3 optic lobes/brains. **f**, Mirr-VP16 *∩ Hmx-DBD* driving *CD4:tdGFP* (FLP out). Anti-GFP (green) and anti-NCad (magenta) show three cells of cluster 76 in a maximum intensity projection. Their morphology matches the morphology of Y13 in the connectome (17). Specifically, the innervation pattern in the layers of the lobula plate indicates Y13 and helps distinguish this morphology from other Y-neuron types. The two panels to the right show the region highlighted by the white box on the left. On the top right, there is a multi-channel representation of cluster 76 (Y13) expressing D (red). The bottom right panel shows D in white. n = 5 /3 optic lobes / brains. **g**, *Wnt4-p65 ∩ Vsx1-DBD* driving *CD4:tdGFP* (MeTu4a). Anti-GFP (green), anti-Toy (blue), and anti-Scro (red) are shown as maximum intensity projections with medulla layer 7 (m7) annotated with a white arrow. The region highlighted by the white box is shown in the top right. This split combination was previously used to annotate cluster 166 as MeTu neurons (46). We found in our new atlas that this intersection is also consistent with cluster 125. MeTu4a neurons express Toy but not Scro indicating that cluster 125 is MeTu4a. Cluster 125 was identified as MeTu4a by their presence in the dorsal region of the optic lobe and the location of axon innervation in the anterior optic tubercle (74) (AOTu) shown in the below right image of *Wnt4-p65 ∩ Vsx1-DBD* in the presence of *brp[SNAPf-tag]* driving *CD4:tdGFP* with labeling of Brp (SNAP, white) and anti-GFP (green). n = 6 brains. (**g**’) *Vsx1-DBD ∩ Wnt4-p65* driving *CD4:tdGFP* (FLP out) (Dm3c). Cluster 166 was identified to be Dm3c by sparse labeling. (Left) Sparsely labeled Dm3c subtype was labeled with this split-Gal4 which innervates medulla layer 3 (m3) (white arrow) and projects from dorsal to ventral side, as shown in the horizontal (top) and frontal view (bottom). (Right, top) Sparsely labeled Dm3c neuron (green) co-expressing Svp (magenta) is shown as a maximum intensity projection. The bottom panel shows a sparsely labeled Dm3c neuron (green) coexpressing Scro (red) but not Toy (blue) in a maximum intensity projection. A single Z-slice of the region in the white box is shown within the image. **h**, *CG15537-p65 ∩ Svp-DBD* driving *CD4:tdGFP*. This split-Gal4 combination is specific to cluster 166, and labels Dm3c, which innervates in the m3 layer of the medulla. Anti-GFP (green) and anti-NCad (magenta) in the top image are shown as maximum intensity projections. The region within the dotted white box shows cell bodies with a higher magnification in the lower panels. The lower right panel shows a maximum intensity projection of anti-Scro (white), matching with the scro expression seen in the Vsx1-Wnt4 split *Gal4* line. n = 3 brains. Scale bars: 10 μm. Developmental stage is indicated to the immediate left of each set of panels.

**Figure S10:**
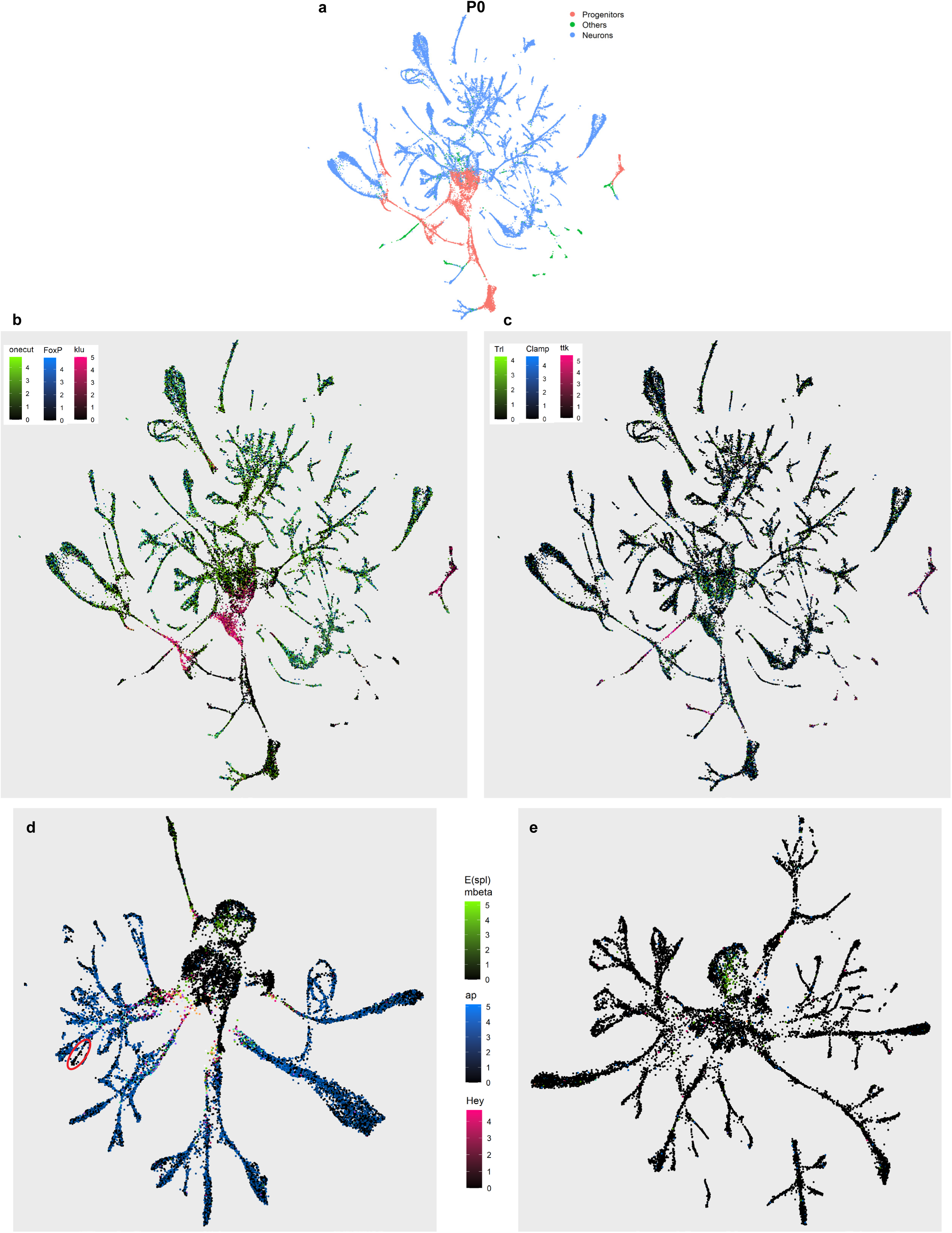
UMAP trajectories of optic lobe neurogenesis. **a-c**, UMAP reduction of the P0 dataset (same as in Fig. 1d), colored by: (**a**) Broad cell type categories (Neurons: blue, Progenitors: red, Others: green), (**b**) Log-normalized mRNA expression of *Onecut*, *FoxP*, and *klu* (green, blue, and red, respectively), (**c**) *Trl*, *Clamp*, and *ttk* expression (green, blue, and red, respectively). **d-e**, UMAP reductions of the Notch^ON^ and Notch^OFF^ hemilineages of the OPC (same as in Fig. 4d-e) and colored based on the log-normalized expression of indicated genes (*Hey*: red, *E(spl)mbeta*: green, *ap*: blue).

**Figure S11:**
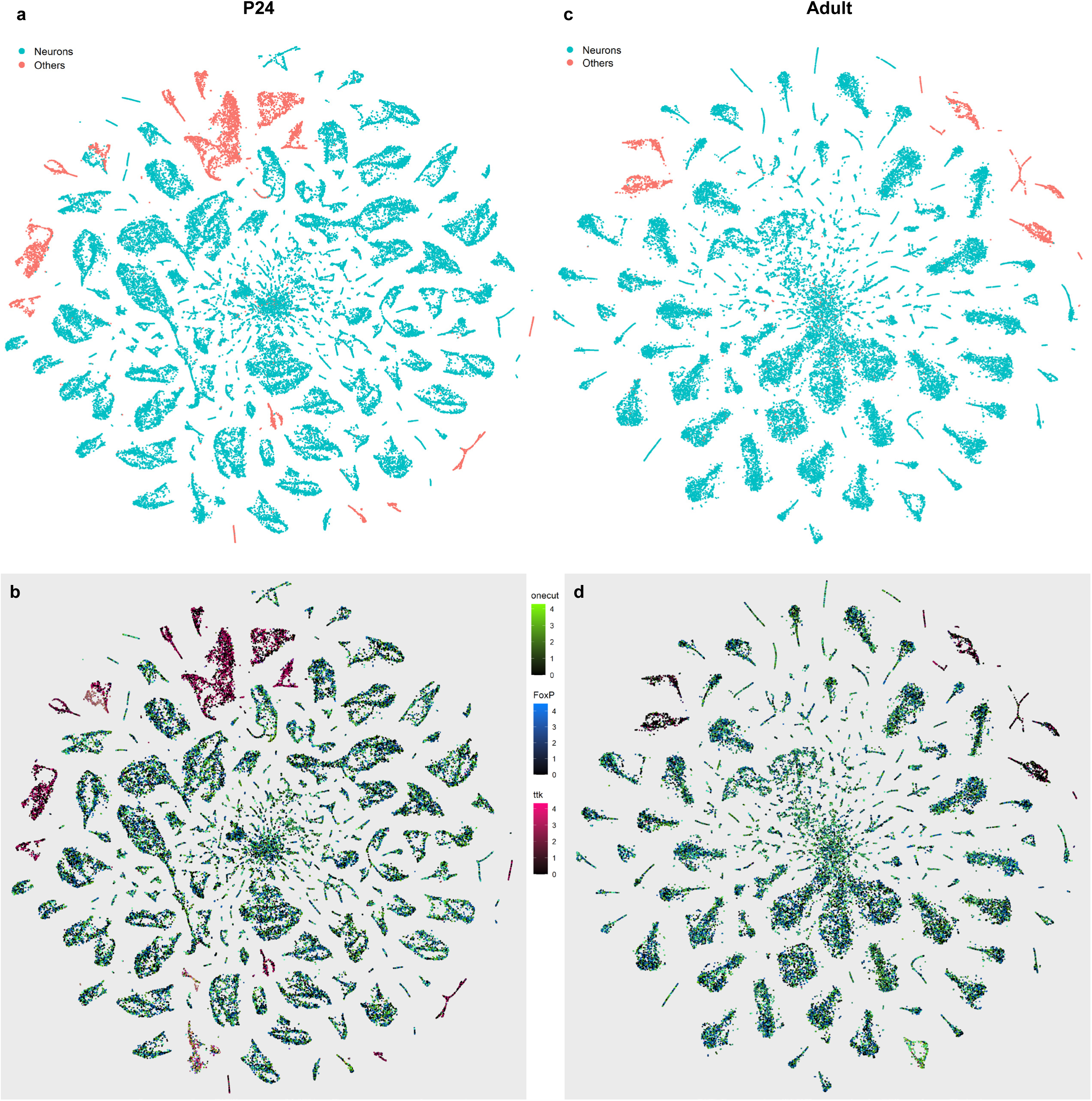
Expression of pan-neuronal genes. **a-d**, tSNE reductions of the P24 (**a-b**) and Adult (**c-d**) multiome datasets based on ATAC modality, calculated using the first 100 LSI components. **a,c**, Broad cell type categories (Neurons: blue, Others: red). **b,d**, Log-normalized mRNA expression of *Onecut*, *FoxP*, and *ttk*.

**Figure S12:**
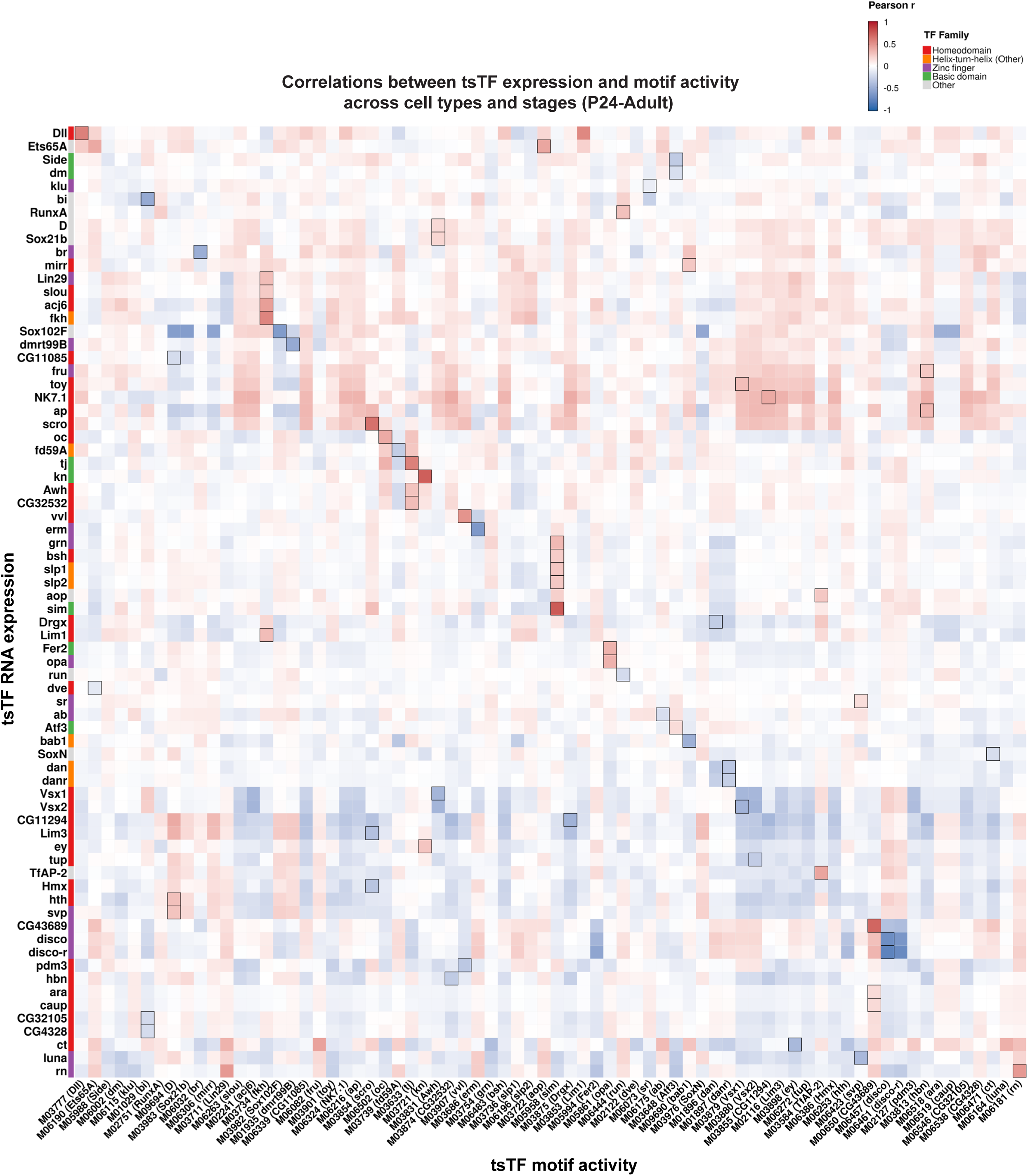
Correlation between tsTF expression and motif activity across cell types and stages. **a**, Heatmap depicting the Pearson correlation (color scale) between the RNA expression of the indicated terminal selector TFs (tsTFs) (rows) and the ChromVAR scores (activity as inferred from chromatin accessibility) of their corresponding motifs (columns). Correlations were computed at the metacell level across neurons from the P24, P48, and Adult stages. Only the most strongly-correlating motif, by absolute correlation, is shown for each of the 72 tsTFs with an available CisBP motif. For each row (TF), its best-correlating column (motif) is outlined with a black square. Rows and columns are hierarchically clustered and displayed in matched order to preserve row-column pairing. Left annotation indicates the TF family (Homeodomain, Helix-turn-helix (Other), Zinc finger, Basic domain, or Other).

**Figure S13:**
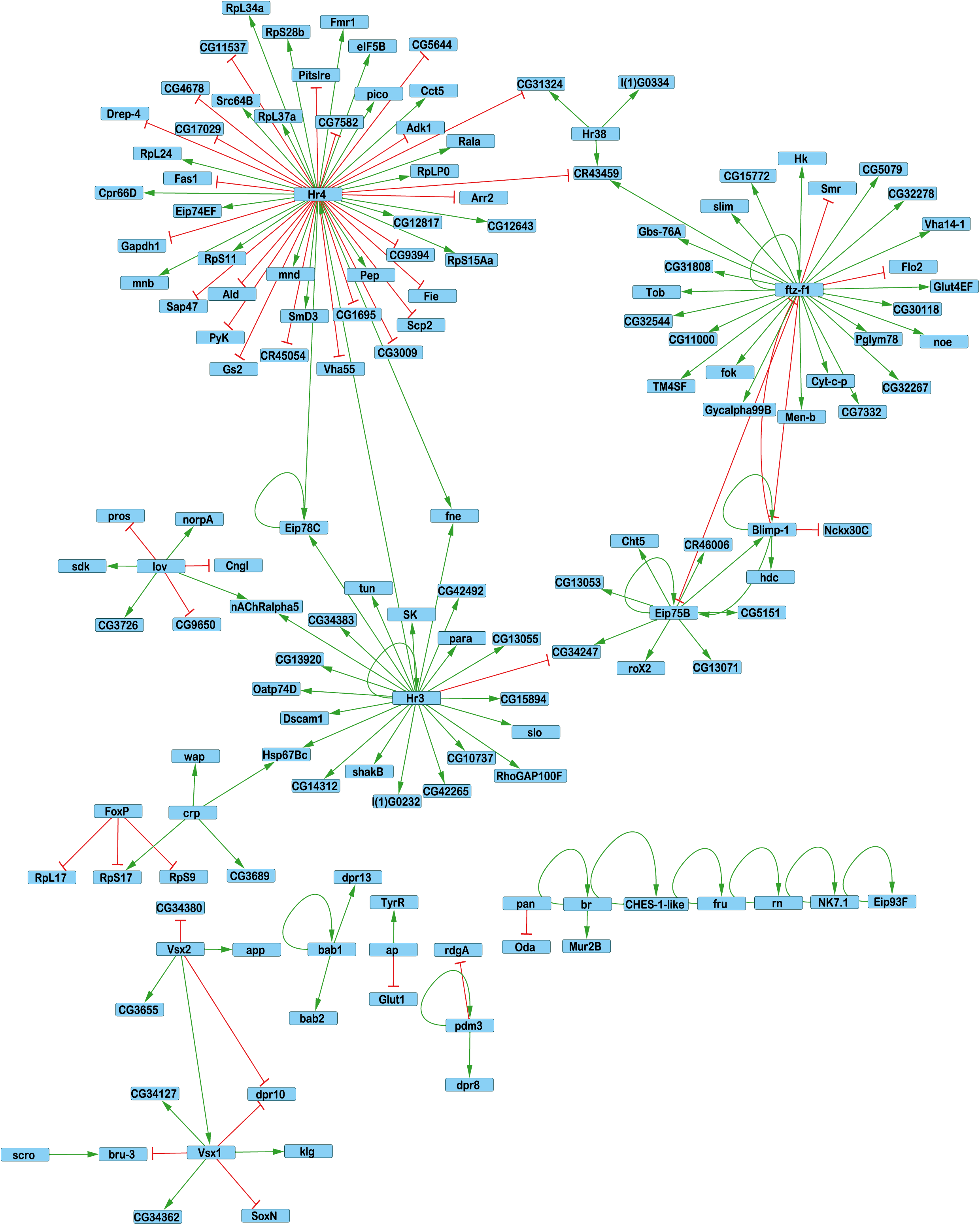
Consistently predicted regulatory interactions by SCENIC+. Cytoscape visualization of regulatory interactions predicted by SCENIC+. Shown are TFs present in at least 10 different networks (excluding M1, and non-neuronal metaclusters M38–M42) and the TF–gene pairs found in more than 50% of the networks that contain the respective TF. Red bars indicate repression; green arrows indicate activation.

**Figure S14:**
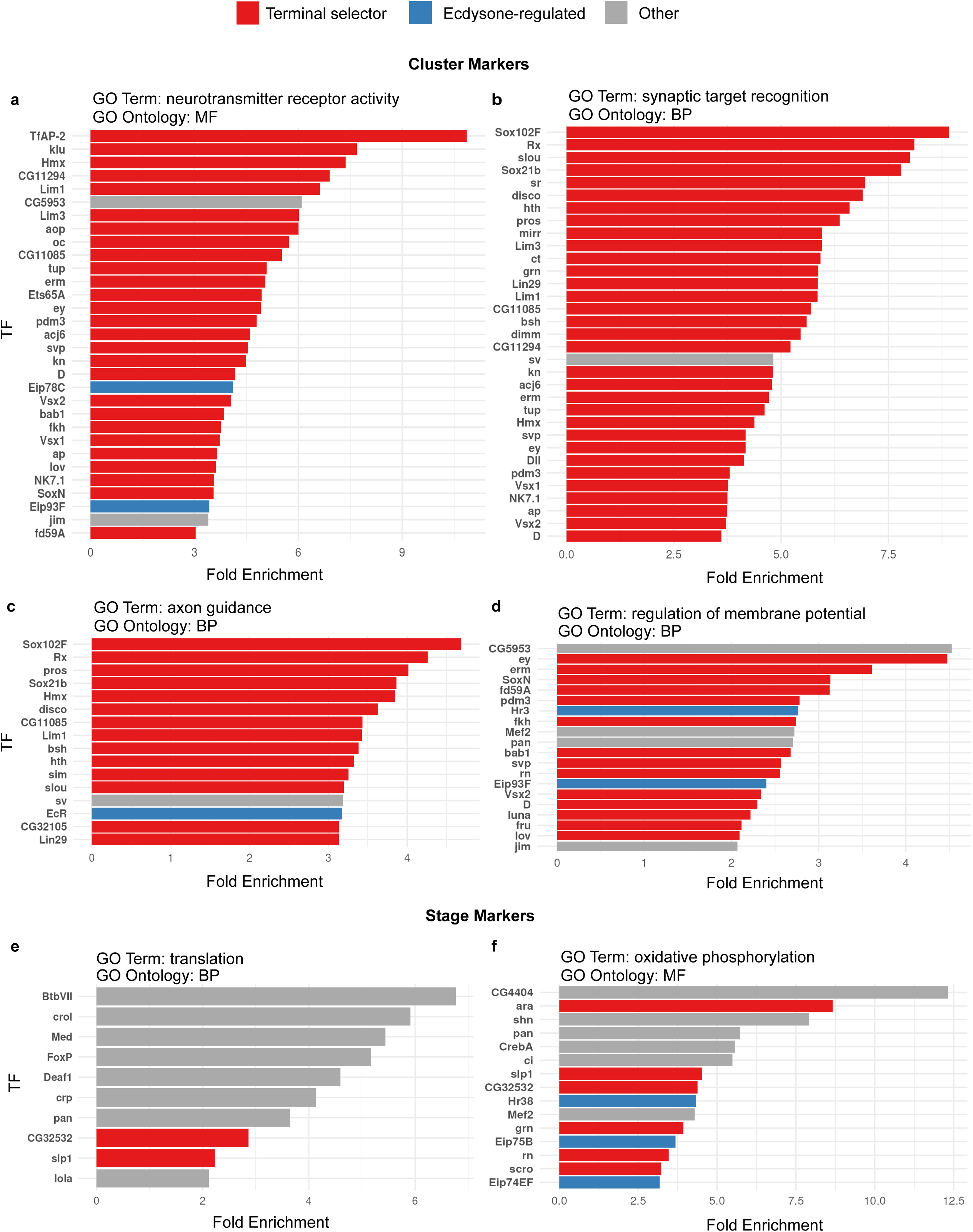
GO enrichment analysis of the predicted target genes for each TF in the SCENIC+ networks. **a–f**, PANTHER GO enrichment terms for target genes of top TFs with at least 50 target genes found across all SCENIC+ networks. Terms are grouped based on prior association with either cluster or stage markers as previously reported(23). Each bar shows the fold enrichment (log₂ scale) of a GO term among each TF’s predicted target genes. TFs are on the y-axis, and bars are colored by TF classification (Terminal selector, Ecdysone-regulated, or Other). Only the significant (FDR < 0.05) molecular function (MF) or biological process (BP) terms are shown. Highlighted terms are (**a**) Neurotransmitter receptor activity (MF); (**b**) Synaptic target recognition (BP); (**c**) Axon guidance (BP), (**d**) Regulation of membrane potential (BP), (**e**) Translation (BP), and (**f**) Oxidative phosphorylation (BP).

**Figure S15:**
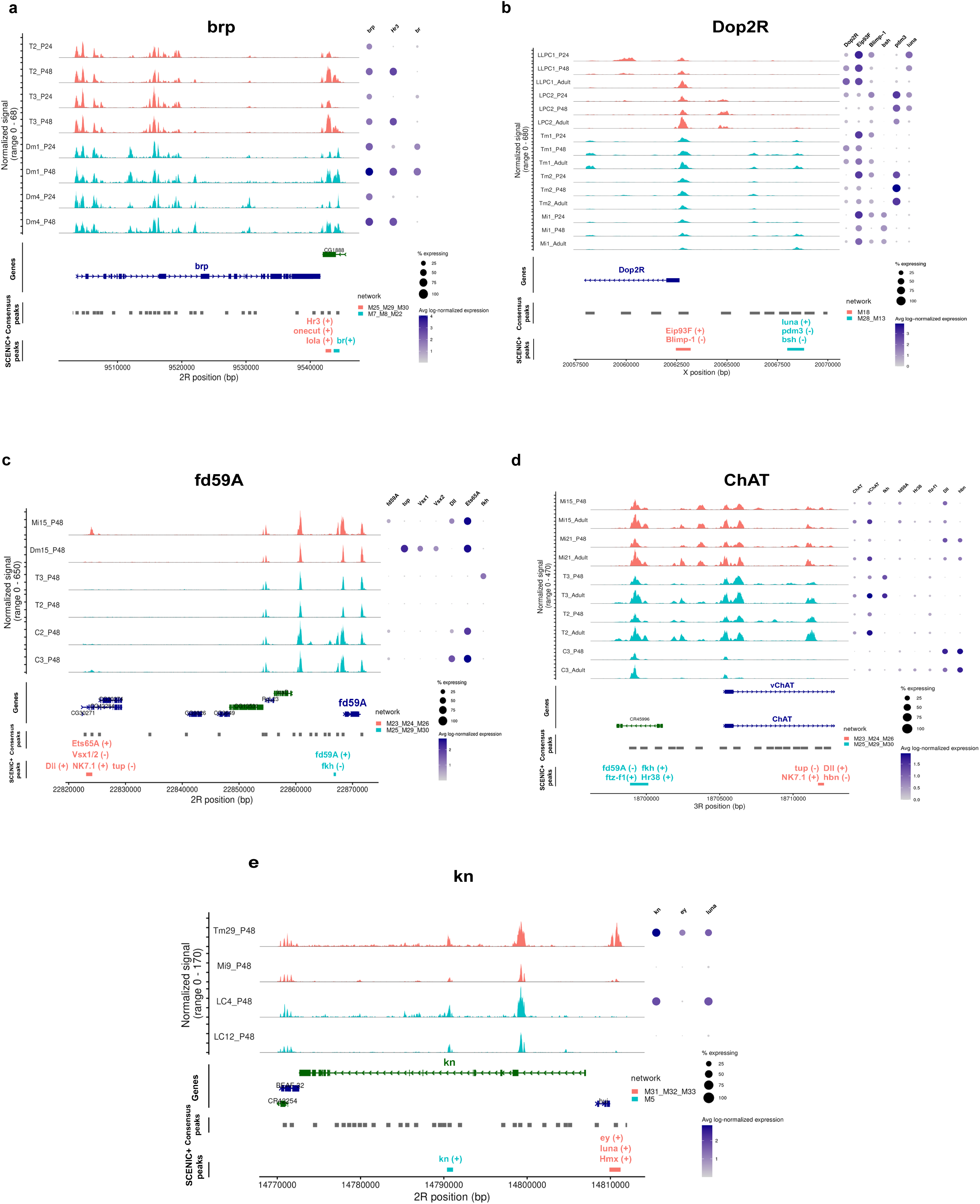
Accessibility and expression profiles of 5 select loci with SCENIC+ predicted regulators. **a-e**, Coverage plots (normalized ATAC signal, range indicated per locus) and matched dot plots for the *brp* (**a**), *Dop2R* (**b**), *fd59A* (**c**), *ChAT* (**d**), and *knot* (**e**) loci, shown for select cell types and stages. Coverage tracks are colored by the respective metacluster that each cell type belongs to. Gray ticks below each gene track denote consensus peaks, and colored bars indicate enhancers with their predicted regulating TF(s) (+, activation; −, repression), colored by the corresponding metacluster network. Dot plots on the right show the percentage of cells expressing each gene (dot size) and its average log-normalized expression (color) in the corresponding cell type and stage.

**Figure S16:**
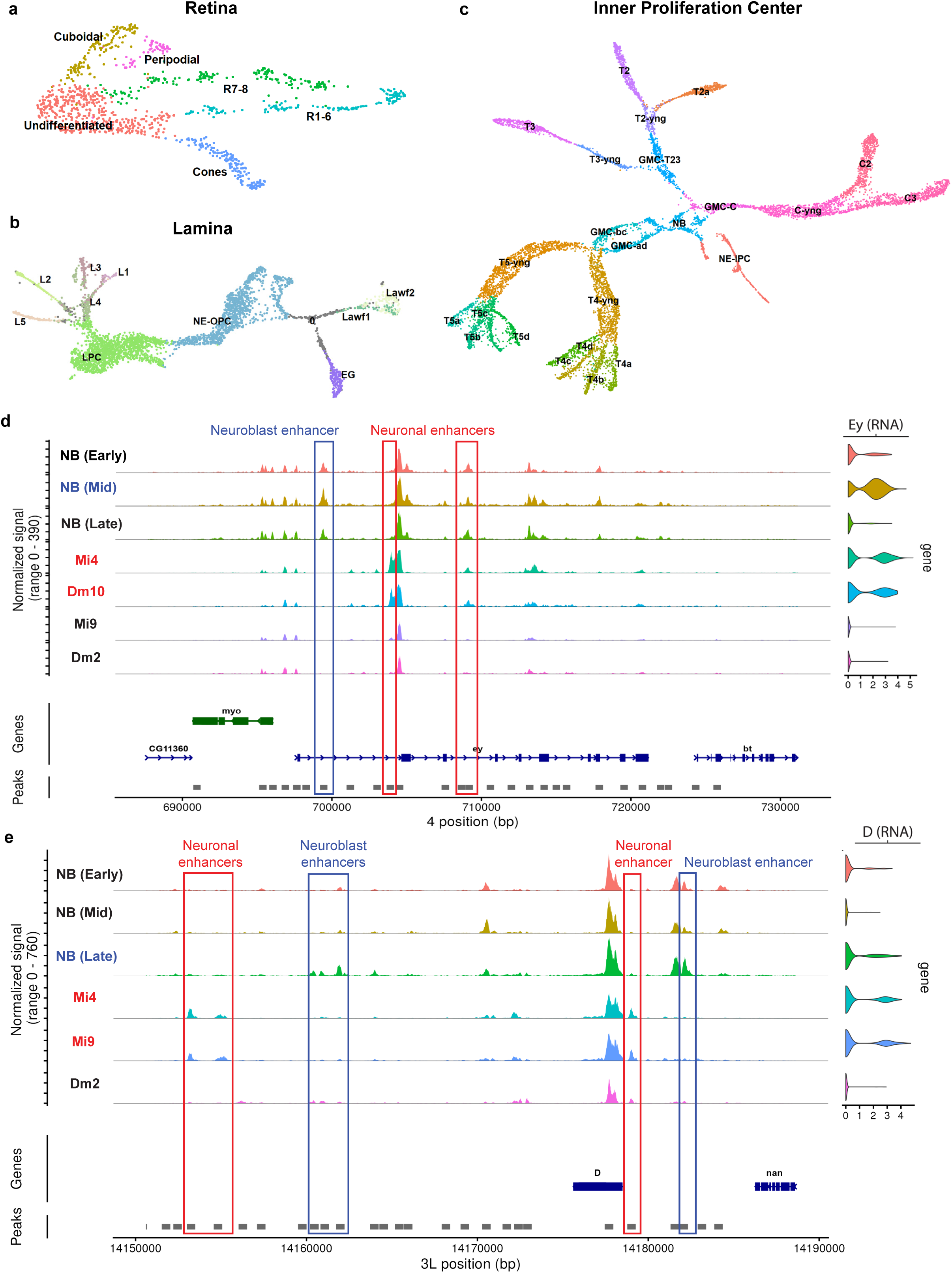
UMAP reductions of other lineages and putative neuroblast and neuronal enhancers of *ey* and *D.* **a-c**, UMAP reductions of single-cell multiomic profiles from the P0 optic lobes (from Fig. 1d). (**a**) Retinal lineage, calculated on 30 PCA and LSI components. (**b**) Lamina (lateral OPC) lineage, calculated on 100 PCA and 90 LSI components. (**c**) IPC lineage, calculated on 20 PCA and 22 LSI components. In **a** and **c**, identities were manually assigned based on WNN clustering and marker gene expression. In **b**, progenitor identities were assigned manually based on clustering and marker genes and neuronal identities were assigned based on label transfer from the P24 dataset (Fig. 1c). Young neurons for which these could not be confidently determined are shown in grey. **d-e**, Coverage plots at P0 showing nearby accessibility for *ey* (**d**), and *D* (**e**) loci, same as in Fig. 4c. NBs were subclustered into their respective temporal windows, and neurons were selected based on the presence or absence of the transcription factor as a terminal selector (tsTF). Blue indicates the NBs expressing the TF, and red indicates the neurons that express the TF as a tsTF. Boxes indicate putative enhancer regions for the respective group, colored accordingly. NE: Neuroepithelia; NB: Neuroblasts; GMC: ganglion mother cells; LPC: lamina precursor cells.

**Figure S17:**
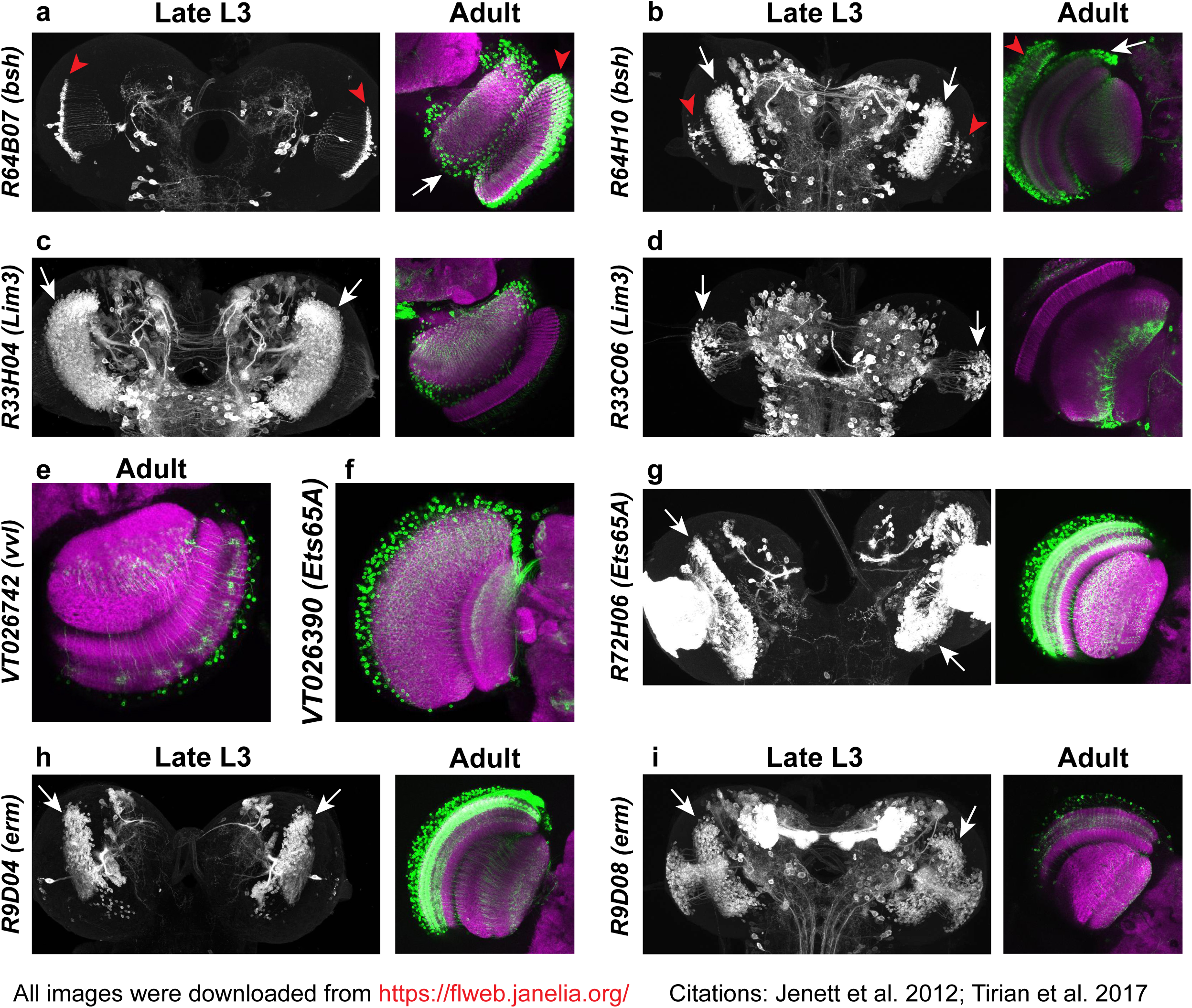
Reporter expression of candidate enhancers for *bsh*, *Lim3*, *vvl*, *Ets65A* and erm. Optic lobe expression of *GAL4* driver lines whose source genomic fragments overlap the candidate enhancers identified for the indicated genes (Fig. 4c–f and Fig. 5b), shown at late L3 (single channel, anti-GFP) and adult (green, reporter; magenta, neuropil) stages: **a,** R64B07 (*bsh*); **b,** R64H10 (*bsh*); **c,** R33H04 (*Lim3*); **d,** R33C06 (*Lim3*); **e,** VT026742 (*vvl*); **f,** VT026390 (*Ets65A*); **g,** R72H06 (*Ets65A*); **h,** R9D04 (*erm*); **i,** R9D08 (*erm*). White arrows indicate reporter expression in the medulla, and red arrowheads in the lamina. Only the adult stage images are available for VT lines (**e**-**f**). All images were downloaded from the Janelia FlyLight database (https://flweb.janelia.org/); the R and VT lines are from Jenett et al., 2012 and Tirian and Dickson, 2017, respectively.

**Figure S18:**
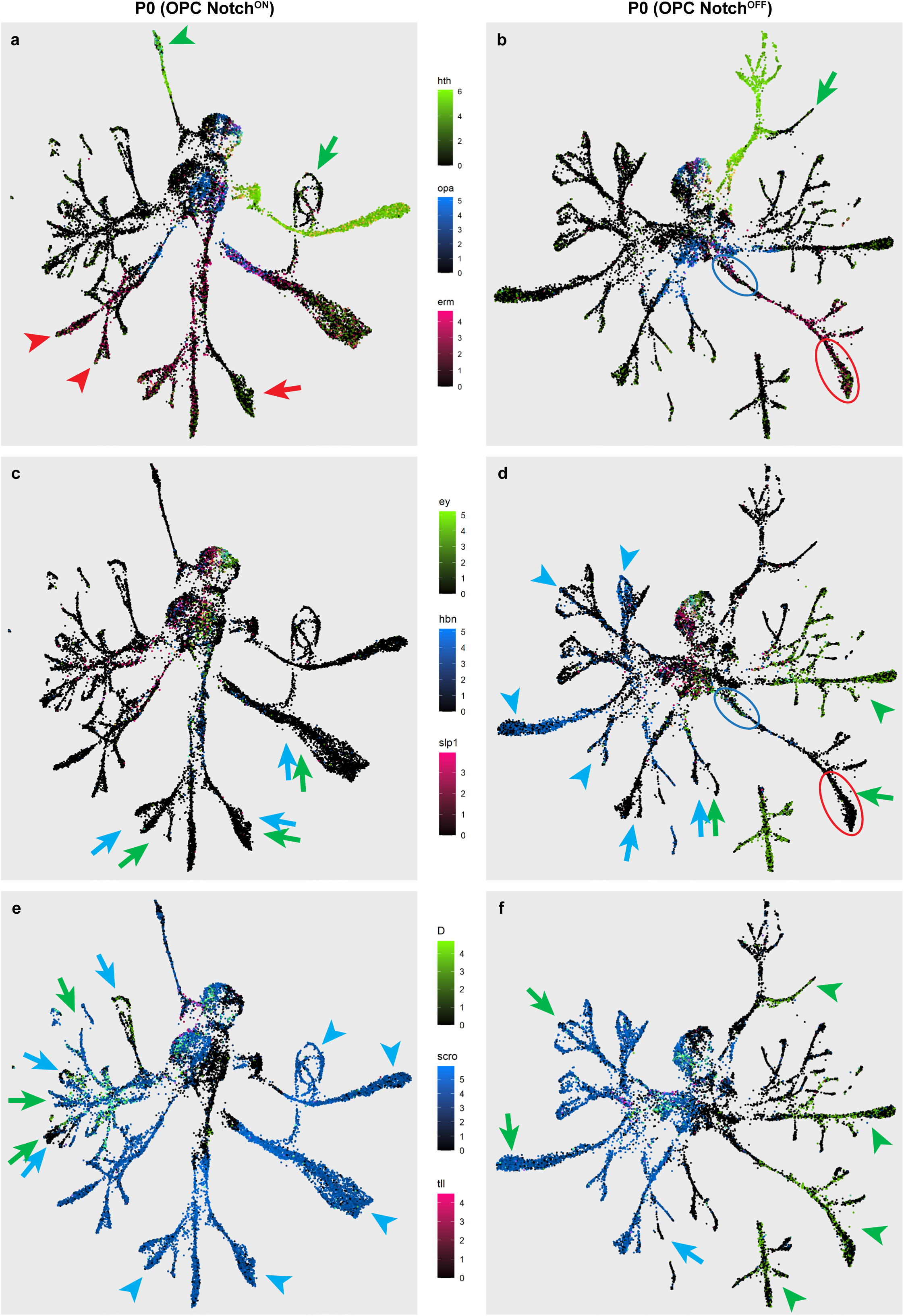
Expression of temporal TF genes during OPC neurogenesis. UMAP reductions of the Notch^ON^ (left) and Notch^OFF^ (right) hemilineages of the OPC (same as in Fig. 4a-b), colored based on the log-normalized mRNA expression of: (**a-b**) *hth* (green), *opa* (blue), and *erm* (red); (**c-d**), *ey* (green), *hbn* (blue), and *slp1* (red); (**e-f**), *Dichaete* (green), *scro* (blue), and *tll* (red). Arrows (colors match the gene legends on each row) indicate neuronal trajectories which do not maintain the expression of the respective tTFs despite originating from these temporal windows. Arrowheads indicate independent activation of these TFs in other lineages.

